# The 2-feature Simultaneous Type/ Serial Token Model of temporal feature conjunctions: Successes are merely fortunate conjunctions

**DOI:** 10.64898/2026.09.07.749788

**Authors:** Mahan Hosseini, Alon Zivony, Martin Eimer, Brad Wyble, Howard Bowman

**Affiliations:** Cognitive Neuroscience, Institute of Neuroscience and Medicine (INM-3), Forschungszentrum Jülich, Jülich 52428, Germany; University of Sheffield, Sheffield S1 4DP, United Kingdom; Birkbeck, University of London, London WC1E 7HX, United Kingdom; Psychology Department, Penn State University, University Park, Pennsylvania 16801, United States of America; School of Psychology and School of Computer Science, University of Birmingham, Birmingham B15 2TT, United Kingdom; Functional Imaging Lab, Imaging Neuroscience, Queen Square Institute of Neurology, University College London, 12 Queen Square, London WC1N 3AR

**Keywords:** temporal feature binding, illusory conjunctions, distractor intrusion errors, selective attention, simultaneous type/ serial token model

## Abstract

Resolving the binding problem, i.e., how we integrate perceptual features in space and time to perceive objects, people and events as coherent wholes instead of masses of free-floating features, has been a central goal of cognitive scientists for decades. Conjunction errors occur whenever features are mis-bound leading to illusory percepts that were not actually presented. A specific kind of Rapid Serial Visual Presentation experiment induces *temporal* illusory conjunctions, called distractor intrusion errors. Despite these errors being well-established, current theories and models provide limited explanations. We present the 2-feature Simultaneous Type/Serial Token (2f-ST^2^) Model, the latest addition to the family of ST^2^-models, which simulates the most important behavioural and electrophysiological distractor intrusion findings. These include response distributions, reaction time patterns, and evoked electrophysiology responses. Importantly, we simulate these results across diverse empirical paradigms, implementing a range of stimulus sets, temporal dynamics, and involving multiple responses after each trial, without needing to modify our model’s architecture or core parameters. Our findings are discussed in relation to theories of selective attention and working-memory binding. We argue that the 2f-ST^2^ model is more parsimonious than the leading theory of distractor-intrusions (the focussed-attention/sophisticated-guessing model), since it uses only a single binding mechanism, implying that correct bindings are simply fortunate conjunctions. The 2f-ST^2^ model further makes predictions for stimulus-competition, subjective-visibility and human electrophysiology. In summary, the ST^2^-model is a general framework for understanding how perception, temporal attention and working-memory interact, and the 2f-ST^2^ model extends the framework to distractor intrusions, thus explaining temporal feature binding.

## Introduction

A central question for perception research is how we perceive coherent integrated objects in our environments. How is it that we perceive “redness” and “cup-ness” together when we see a “red cup”? How this “binding problem” is resolved in the brain has challenged scientists for many decades (Feldman, 2013; Treisman, 1998). What is clear is that the precision of our perceptual system in binding features together is of critical importance to our survival, governing our capacity to navigate and act upon our environment. For example, when witnessing a crime, in a flurry of moving figures, correctly associating who wore what and who held what object is of critical importance for future eye-witness testimony.

The brain has to deal with two dimensions to arrive at coherent percepts: space and time. That is, in general, it perceives spatially collocated and temporally coincidental features as a single coherent whole. The importance of such binding for the functioning of our perceptual systems is evident from cases in which it goes wrong. In healthy subjects, *illusory conjunctions or conjunction errors* (Botella et al., 2001; Botella & Eriksen, 1992; Henderson & McClelland, 2020; Klein et al., 2023; Pelli et al., 2004; Treisman & Gelade, 1980, Zivony & Eimer, 2020, 2021) occur whenever subjects incorrectly combine the features of two or more presented stimuli, whereas chronic impairments of feature binding are a central symptom of certain neuropsychological disorders, such as Schizophrenia (Su et al., 2015) and visual neglect (Battelli et al., 2007). We focus here on the problem of temporal association of features, since theories of spatial associations tend to have been more explored than their temporal counter-parts (Scholte & De Haan, 2025). A standard way to understand a mechanism is to explore it at the limits of its normal functioning (e.g. Particle Accelerators in Physics). In perception research, this can be done by presenting multi-feature stimuli very rapidly one after the other at the same spatial location, in so called Rapid Serial Visual Presentation (RSVP; Lawrence, 1971; Potter, 1976). Certain kinds of RSVP experiments induce a particular type of *temporal* illusory conjunctions, called *distractor intrusion* errors (Botella, 1992; Botella et al., 2001; Botella & Eriksen, 1992; Chennu et al., 2011; Zivony & Eimer, 2021), in which a distractor item is erroneously reported instead of the target. These types of errors are important, because they offer a window into the architecture and temporal dynamics of the cognitive and neural mechanisms that are responsible for the integration of visual features across time. Understanding the processes that produce such distractor intrusions can therefore provide critical, more general insights into how cognitive systems such as selective attention and working memory operate to generate veridical representations of visual objects in space and time. In this article, we will first introduce the experimental procedures that have been used to study distractor intrusions. Next, we will discuss current empirical and theoretical accounts of why such errors occur, and highlight the limitations of these accounts. In the main parts of this article, we describe our own computational model of selective attention and working memory and its recent adaptations that were specifically introduced to account for distractor intrusions. We present a number of simulations using our model that have successfully replicated a wide range of empirical results from experiments with human participants, including both behavioural and electrophysiological data. We conclude that this model can provide a general framework for understanding how perception, temporal attention and working memory operate in dynamic visual environments.

### Distractor Intrusion Errors in Rapid Serial Visual Presentation (RSVP) Experiments

Distractor intrusion errors are a result of an erroneous binding of (typically) two feature dimensions of two separate visual stimuli (one being a distractor) into a combined percept of one stimulus that has not been presented in the RSVP stream (Botella, 1992; Chennu et al., 2009; Zivony & Eimer, 2024). In typical RSVP experiments in which intrusion errors can occur, participants search for a target based on a feature that differentiates the target from the distractors (e.g. the colour red), and report another of the object’s features (e.g. its identity). Figure 1 (inspired by Figure 1 in Botella et al. (2011)) depicts the central stimulus set of a typical RSVP stream of coloured letters, with digits next to stimulus frames indicating item-positions with respect to the target, which is at position 0. Participants’ task would be to report the red letter after the stream concluded, making the target frame the red ‘S’. In distractor intrusion experiments, we refer to the stimulus-feature that differentiates the target from the distractors as the key feature (here, the colour red) and the stimulus-feature that needs to be reported as the response feature (here, the S-shape). Distractor intrusion errors arise if participants report the identity of a (typically) neighbouring distractor stimulus instead of the ‘S’.

**Figure 1.**
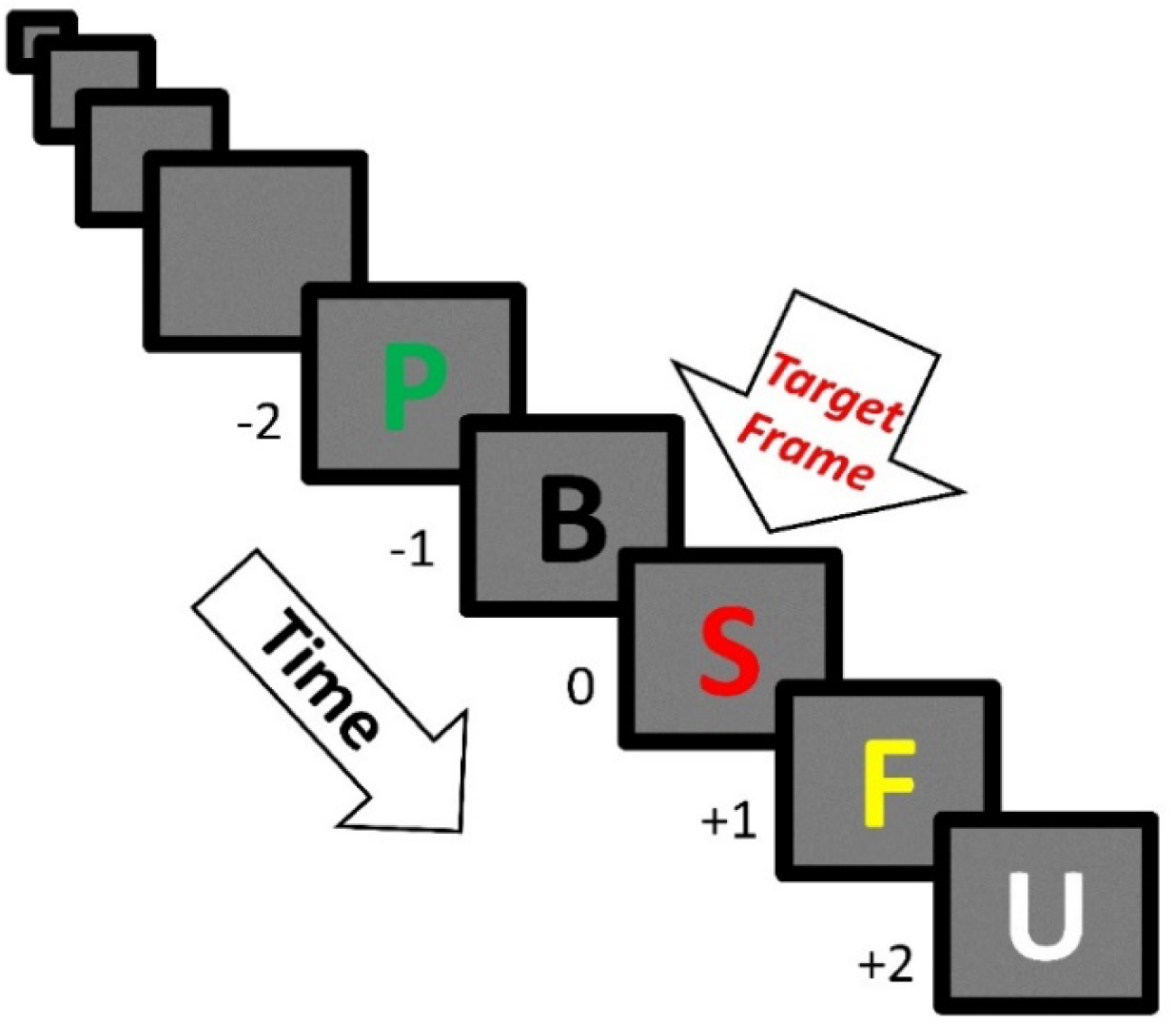
Example RSVP Stream that could generate distractor intrusion errors. Time unfolds from top to bottom. In this example, the task would be to report the red letter and we refer to the frame as the set of features appearing at each position in the sequence. Frames are numbered relative to their position before (negative) or after (positive) the ‘S’ stimulus that is the target. Intrusion errors are made if participants erroneously report a distractor stimulus as being red instead of the target. For example, a +1-intrusion error is made if the ‘F’, which immediately follows the red ‘S’ target frame, was reported as being the red letter. Likewise, a −2-intrusion error is made if the ‘P’ was reported as being red.

Whilst distractor intrusions have been documented in many studies, very few theoretical accounts and even fewer formal accounts of distractor intrusions have been put forward. A first observation is that the required system distinguishes between different features in a compositional sense that allows them to recombine. One possibility for such an architecture is a *separated-feature structure* in which the processing of RSVP items must split into multiple pathways, one for each feature dimension; see Figure 2A. Then, differences in the temporal dynamics of these two pathways desynchronise the representation of features that were presented together. These separate pathways then recombine (see Binding in Figure 2A) at a stage that is likely to be an immediate or close precursor to conscious perception. The “loss of representational synchrony” between the two feature-pathways is what leads to mis-bindings at the Binding module and distractor intrusion errors being experienced. Figure 2B illustrates how key and response feature would arise in the separated-feature structure, with key feature detection determining which response feature is bound; see large horizontal dashed arrow.

**Figure 2.**
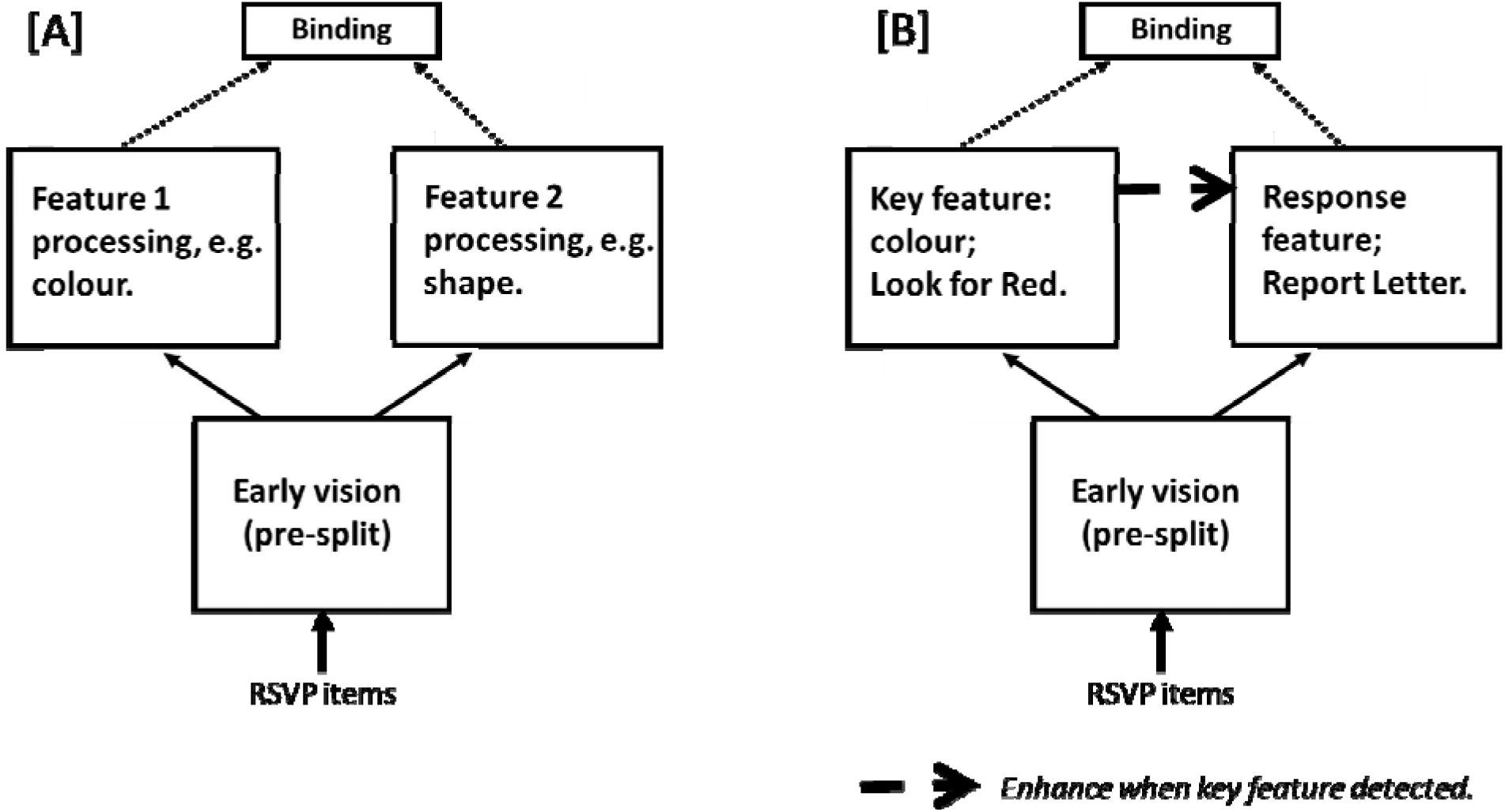
Separated-feature structure suggested by intrusion errors. [A] Early vision processes of a given RSVP item are shared, before processing splits into multiple pathways, one for each feature dimension present in the stimulus. A final binding process then binds the items’ features into an integrated representation that is perceived. Loss of representation synchrony between the two features during the split is an obvious contributor to intrusion errors. [B] In a typical intrusion error experiment, task instruction enforces one pathway to implement the *key feature* and the other pathway the *response feature*, with the detection of the key feature determining the response feature to bind in the final stage (see large horizontal dashed arrow) under ideal conditions.

Key feature detection is a classic example of attentional selection. The system is searching through time for a target key feature and attention is deployed when that feature is detected. Thus, latency variance in key feature processing is an index of temporal variance in attentional selection. Furthermore, it is widely agreed that attention plays a major role in feature binding. Specifically, theories of attention have long suggested that attending to a specific location ensures that the features occupying that space will be bound together, e.g. feature integration theory (Treisman & Gelade, 1980; Kristjánsson & Egeth, 2020). Evidence for this claim comes from studies showing that when attention is engaged elsewhere, binding can go awry, resulting in illusory conjunctions of features (Treisman & Schmidt, 1982; and Chun, 1997, which exhibits this phenomenon in time). However, as previously indicated, spatial attention cannot fully account for binding. To perceive coherent objects in a changing world, it is not enough to track whether features are spatially collocated—whether features are temporally co-occurring must also be tracked. In other words, to resolve the binding problem, our perceptual system must deal with two dimensions: both space and time. Relative to spatial attention, there has been less focus on the role of temporal attention—attention to specific points in time.

Many accounts of temporal attention view attentional selection as a temporally discrete process that separates perception to distinct stages (Seibold et al., 2023). In contrast, a subset of theories (the ST^2^ model (Bowman & Wyble, 2007), the episodic-ST^2^ model (Wyble et al., 2009), Olivers and Meeter’s (2008) boost and bounce model, and Reeves and Sperling’s (1986) model), suggest that attention is a modulatory process that unfolds over time. Such theories have recently been termed “diachronic” theories of attention (Zivony & Eimer, 2021). A key feature shared by these theories is the notion that detecting the target’s key feature initiates transient attentional enhancement (TAE), a relatively brief (phasic) enhancement that substantially strengthens sensory representations, propelling some into working memory, with the N2pc being proposed as the likely electrophysiological analogue of this deployment (Eimer, 2014).

In contrast to the key pathway, the response pathway reflects processing sufficient for encoding into working memory and conscious experience. This could involve competition for access to binding, if multiple responses are enabled, perhaps especially for access to conscious experience (Bowman et al., 2022).

Over the last couple of decades, several theoretical accounts as well as computational models explaining intrusion errors have been presented. We will introduce each of these next.

### Botella et al.’s (2001) focussed attention-sophisticated guessing model

The first model proposed to account for distractor intrusion errors in a temporal stream is the *focussed attention-sophisticated guessing* model of Botella et al. (2001). Apart from a precursor to the model being presented here (Chennu et al., 2011), the *focussed attention-sophisticated guessing* model is the one extant *computational* model of distractor intrusion errors. Botella et al.’s (2001), model proposes two distinct mechanisms which, although related, are not direct analogues of the key and response pathways shown in Figure 2. In Figure 2, the pathways differed based on which *feature is processed,* whereas the two mechanisms in Botella’s model differ based on *which response is generated*. The suggestion was that these mechanisms produce two qualitatively different kinds of processing: guesses based upon a *sophisticated guessing module* and “by-construction” correct responses based upon a *focussed attention module*. Botella et al. (2001)’s model consists of four main elements: module K for the key feature, module R for the response feature, a focussing mechanism and a sophisticated guessing mechanism. Modules K and R act in parallel, extracting key and response features, respectively. The focusing mechanism activates attentional processing once a task-relevant key feature has been detected, which the authors call t_c_, the critical moment. This mechanism, however, fails at times, especially if presentation rates are fast. When this happens, sophisticated guessing acts as a backup by selecting the item with the highest module R activation at the critical moment, t_c_. Crucially, correct reports occur *either* after successful completion of the focussing mechanism *or* after sophisticated guessing happens to select the correct item in module R, as its representation had the highest activation at time t_c_. The latter case is described by the authors as a *fortunate conjunction* (conjunctions being synonymous with intrusions), as correct reports were given after guessing. In contrast to correct reports, distractor intrusions are never reported by the focussing mechanism, but exclusively happen when sophisticated guessing fails to select the correct correspondence (Botella et al., 2001).

One important piece of evidence in favour of this account comes from a speeded reaction time (RT) intrusion task. Unlike most studies of distractor intrusions, participants had to report the target as accurately and as quickly as possible. The results of this study showed that pre-target intrusion errors were made later than correct reports, despite the fact that pre-target intruder items were present *earlier* than target items. This finding was argued to suggest a mixture of short and long trials involving correct reports, supporting the authors’ dual-mechanism model, as fast and slow correct reports are proposed to be the result of successful focussing and sophisticated guessing, respectively (Botella et al., 2001). We will call this phenomenon the *counterintuitive reaction time finding*, and it will provide a critical test for the computational model we propose, since thus far no other model or theory is able to replicate or explain this finding.

We highlight two issues with Botella et al.’s (2001) model. First, it is less parsimonious than one that only assumes distinct key and response pathways as in Figure 2. In particular, *focussed attention* is effectively an additional mechanism to the general structure of Figure 2 Second, and most problematically, in the *focussed attention-sophisticated guessing* model, if one takes the structure in Figure 3 literally, correct responses obtained via the focussed attention pathway are generated *without indication of how response features are accessed*, since these are represented in Module R, which is not accessed with focussed-attention. This seems to suggest that there is a pathway (the focussed attention pathway) that can keep the key and response features together throughout the cortical stages of stimulus-processing. Physiologically, it is difficult to imagine what would support such an error-free pathway, since different feature dimensions are processed in distinct regions in visual cortex (Op de Beeck et al., 2008; Zeki, 1978). One might suggest that a mechanism, such as, oscillatory synchrony (Tallon-Baudy, 2003) binds the features and only does this successfully for the focussed attention route. For example, the association of physically separated neurons in the brain could be maintained by those neurons oscillating at the same frequency and in phase with one another, giving a physiological marker that neurons are “bound”. However, does any 100% accurate neural mechanism exist, given the fundamentally stochastic nature of neural processes?

**Figure 3.**
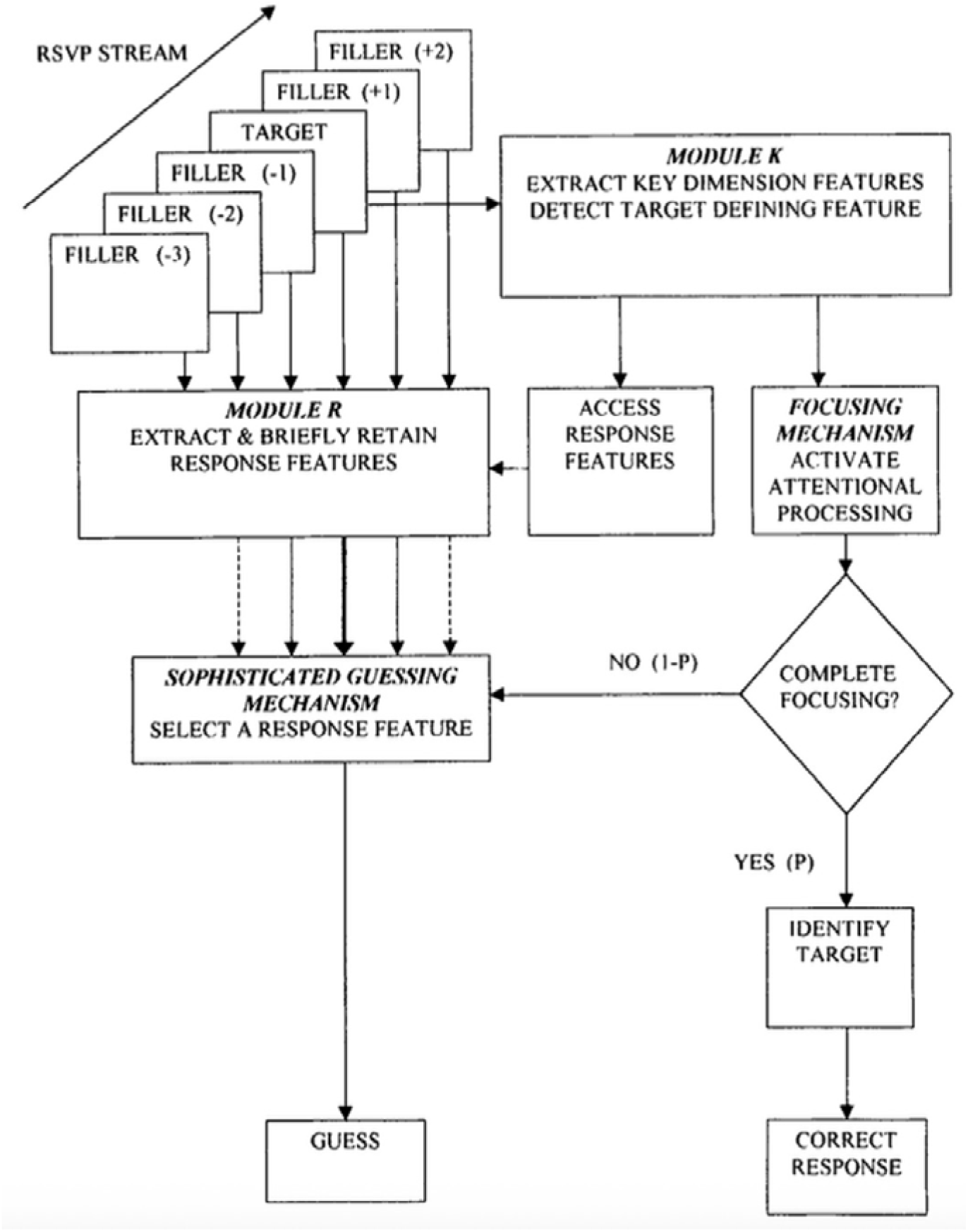
Botella et al. (2001)’s model of distractor intrusions, involving four main elements: module K, module R, a focussing mechanism and a sophisticated guessing mechanism. Modules K and R extract key and response features, respectively. The focusing mechanism activates attentional processing once a task-relevant key feature has been detected. In cases in which the focusing mechanism fails, the sophisticated guessing selects the item with the highest module R activation.

This challenge provides a central motivation for the model we are proposing here. Specifically, we will present a model in which there is no sharp qualitative distinction between correct responses and errors and all of the computational mechanisms will be formally specified. Instead, we will argue that *all* correct responses are simply *fortunate conjunctions*, while errors are simply *unfortunate conjunctions*. Nonetheless, we will show that this model can generate the counterintuitive reaction time finding that partly motivated the *focussed attention-sophisticated guessing* model. Specifically, the model we propose here, was presented after the Botella model in an initial version (Chennu et al, 2011) and generalises the Simultaneous Type/ Serial Token (ST^2^) model first presented in Bowman and Wyble (2007) to temporal feature binding. Next, we present the ST^2^ model in its original form, in preparation for the additions we will make, when we describe the full two-feature version of the ST^2^ model in section “The 2-feature Simultaneous Type/ Serial Token Neural Model”.

### Simultaneous Type/ Serial Token Models (Bowman & Wyble, 2007)

The ST^2^ model of temporal attention and working memory (Bowman & Wyble, 2007), and its derivative, the eST^2^ model (Wyble et al., 2009), simulate a large spectrum of phenomena, including, the attentional blink (AB) (Bowman & Wyble, 2007); lag-1 sparing (Bowman & Wyble, 2007); spreading the sparing (Wyble et al., 2009); repetition blindness (Wyble et al., 2009); perceptual episodes (Wyble et al., 2011); delayed consolidation during the AB (Bowman et al., 2008); P3 event related potential effects during the attentional blink (Chennu et al., 2009); loss of temporal precision of the P3 during the AB (Chennu et al., 2009); the experiential blink (Jones et al., 2020; Pincham et al., 2016); and the meta-experiential blink (Bowman et al., 2022).

A key concept in the ST^2^ model is the distinction between *types* and *tokens*, which, in cognitive psychology, has its origins in the repetition blindness findings of Kanwisher (1987). Types reflect the featural representation of a stimulus, encompassing all featural properties of it, whereas tokens indicate the episodic context in which a stimulus occurs, for example by recording the temporal sequence of stimuli. For example, if a person saw the letter ‘K’ followed by ‘M’, a type for the letter ‘K’ would represent all of its identifying information such as its visual form, the fact that it is a letter and also a consonant. Token information would indicate that the letter ‘K’ occurred once, and that it appeared *before* the letter ‘M’. If a person perceived two distinct K’s presented in sequence, there would typically be one type, corresponding to the K’s identifying information, and two tokens, one corresponding to each of the two instances. Types reflect information that is instance-invariant, i.e. that is always present when that item (e.g. letter) occurs; tokens are instance-specific and thus fundamentally episodic.

The ST^2^ model presented in Bowman and Wyble (2007) is shown in Figure 4. The model is divided into two stages: a first that (abstractly) models the (ventral) visual-processing stream and a second that models encoding into working memory. See also Chun & Potter (1995) for a similar account of such a two-stage model. In later treatments of the theory (Chennu et al., 2009), we have associated the conscious experience of a stimulus/type with this encoding into working memory. Although, see Bowman et al. (2022) and Jones et al. (2020) for a more nuanced position on this association.

**Figure 4.**
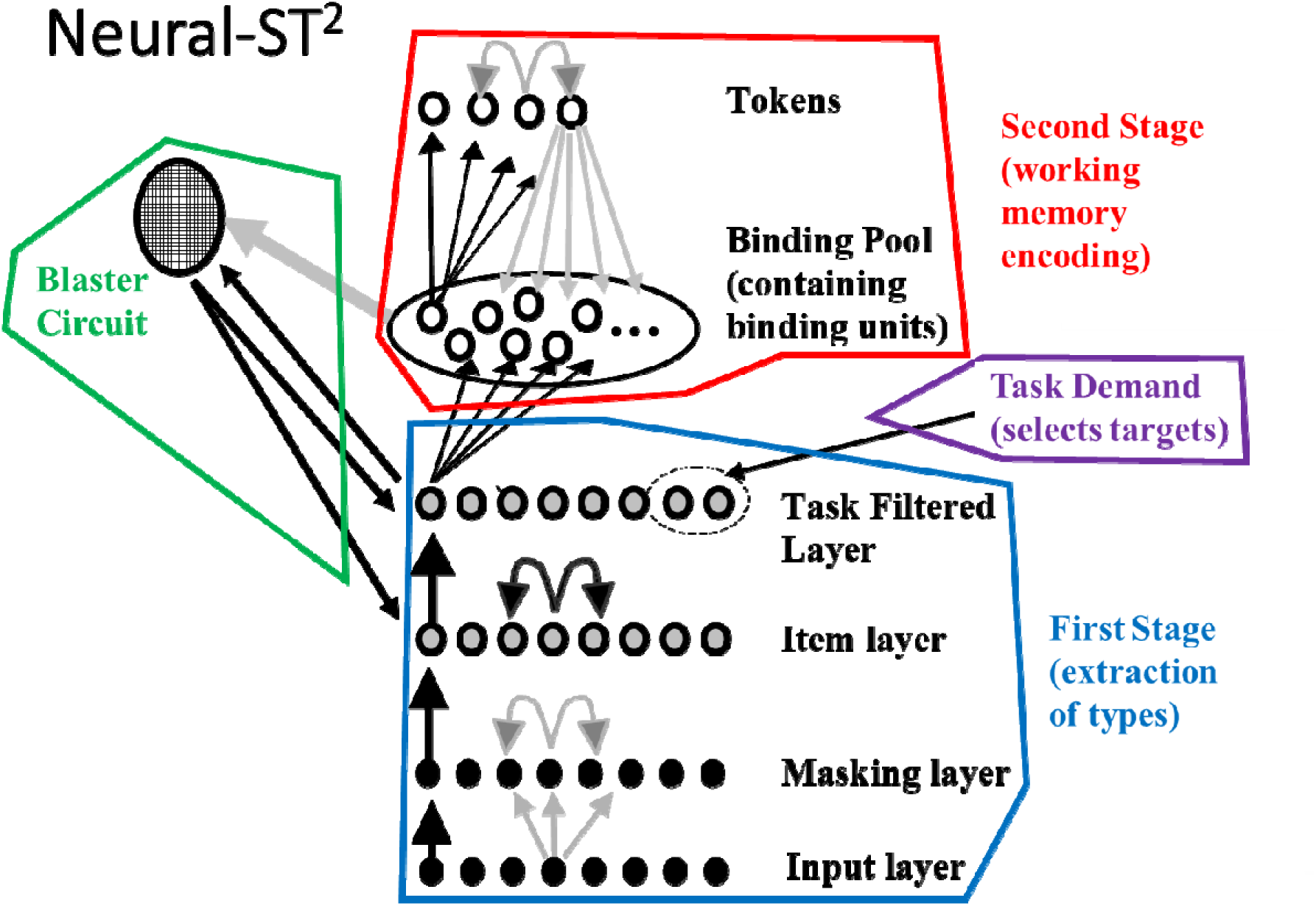
Depiction of original ST^2^ model, as presented in Bowman and Wyble (2007). The four main components of the model are demarcated (see online version for colour demarcation). For details of different kinds of nodes and kinds of arrows see Bowman and Wyble (2007). These details are not central to the intuitive understanding of the ST^2^ model required at this point in this paper. The first stage provides an abstract model of the visual processing pathway, with representations sufficiently active at the final (Tasked Filtered) layer entering the second stage and thereby being encoded into working memory. This Stage 2 process of encoding involves associating final layer of Stage 1 representations (i.e. a type) to the currently available token, thereby associating a type with an episodic marker, indicating that the type has occurred and where in the sequence of type occurrences it happened. The task demand system ensures that only task relevant types can pass into Stage 2, and the blaster implements transient attentional enhancement. This transient enhancement effectively acts as a gate: amplifying Stage 1 representations for a brief period of time following the detection of a target. This detection occurs when a type in Stage 1 is consistent with the task set and thus fires the blaster.

The first stage is simultaneous, such that, multiple stimuli can be active at the same time, with little interference between them. This simultaneity is required to explain cases where closely presented stimuli can be encoded into working memory with little cost to accuracy, such as lag-1 sparing (Bowman & Wyble, 2007), spreading the sparing (Wyble et al., 2009) and perceptual episodes (Wyble et al., 2011). The type stage is similar in some respects to the conceptual short-term memory or CSTM, described by Potter (1976), in that incoming information can be very rapidly processed to extract its identifying information, but without a subsequent stage of memory encoding this information will almost immediately be forgotten.

The conscious experience and encoding into working memory performed by the second stage of the model involves three mechanisms. Firstly, only salient stimuli progress from 1^st^ to 2^nd^ stage, with a task demand unit enforcing task instructions, e.g. that red items are targets. Secondly, a transient attentional enhancement mechanism, called the blaster, enhances representations in Stage 1 when a salient stimulus is detected, thereby “opening the gate” to Stage 2. This transient enhancement implements temporal selection, i.e. a temporal spotlight. Finally, a pool of binding units bind the features of the representation experienced, encoding them together into working memory. This paper considers how these basic ST^2^ mechanisms can be enhanced by adding key and response pathways in Stage 1, in order to model distractor intrusions.

Consistent with the perspective that the P3 represents a conscious perceptual event (e.g. Pincham, Bowman, Szucs, 2016) and with previous work, we associate the P3 with activation of the late stages of the model (Chennu, Craston, Wyble & Bowman, 2009; Craston, Wyble, Chennu & Bowman, 2008; Bowman, Jones, Pincham, Fleming, Cleeremans & Smith, 2022; and Jones, Pincham, Gootjes-Dreesbach & Bowman, 2020). In this paper, we make a further association between the firing of the blaster and the N2pc. As elaborated on later, this is consistent with the perspective that the N2pc reflects the deployment of attention (Kiss et al., 2008).

As previously highlighted, the ST^2^ model, as presented in Bowman and Wyble (2007), was extended in preliminary form (Chennu et al, 2011) to model experiments with multiple feature dimensions and how errors of binding could lead to intrusion errors. We enhanced that preliminary model further to give us the model that is the centre-piece of this paper. This further refinement is presented in section “The 2-feature Simultaneous Type/ Serial Token Neural Model”.

### Additional Theories of Distractor Intrusion Errors

Zivony and Eimer (2020, 2021) proposed a theoretical account, which is closely related to the ST^2^ models, and in which a transient attentional enhancement (TAE) mechanism is deployed once sufficient evidence about the target’s key-feature has accumulated (similar to the ST^2^ model’s task-relevant type activating the blaster circuit). When Zivony and Eimer’s (2020, 2021) TAE is deployed, it enhances stimulus-representations before gradually dissipating (similar to the blaster circuit enhancing late Stage 1 activation levels in ST^2^ models). In Zivony and Eimer’s (2020, 2021) RSVP experiments, category-matching distractor items (i.e., intruder-items), which share the response dimension of the target item, were exclusively presented in the frame *immediately following* that of the target item. As a result, only +1 post-target intrusion errors were possible. Crucially, temporal variability of the TAE across trials can strongly affect visual awareness and perceptual reports. Post-target intrusion errors are argued to occur if attentional engagement is substantially delayed. This is because the delayed TAE enhancement particularly benefits the intruder item causing it to win the competition with the target item, being encoded into WM, and thus reported.

The authors supported this notion with an examination of the temporal dynamics of the attentional system after post-target intrusion errors (Zivony & Eimer, 2021). The N2pc ERP-component was measured as an index of attention being allocated to visual objects carrying task-relevant features (Eimer, 1996; Woodman & Luck, 1999). Zivony and Eimer (2021) showed that the N2pc’s onset latency was significantly delayed in distractor intrusion trials compared to correct response trials, providing evidence for post-target intrusions being a result of delayed TAE enhancement. However, the authors modified their initial encoding account (Zivony & Eimer, 2021) after novel empirical data, specifically changing the assumption that *only the winning item is encoded into WM* (Zivony & Eimer, 2020).

Vul and colleagues (Vul et al., 2009) empirically examined a number of alternative ways in which visual selective attention might operate and proposed a contiguous-graded selection process. This account implies that once the correct key (also called selection) feature is detected, a time-window of selection is placed around the stimulus carrying the correct key feature (i.e., the target). Item representations of target as well as proximal distractors, which occur during the postulated time-window of selection, are then temporarily stored in short-term memory. One of these items is subsequently sampled to be processed further, allowing its identification as well as report. Which item is selected for further processing (and, ultimately, response) involves sampling from a probability distribution^1^ across the items in short-term memory. This account was mostly based on their Experiment 1, an RSVP experiment in which intrusion errors were possible and subjects were required to guess the identity of the target stimulus four times (the most likely, then the next most likely and so on) after each trial. Importantly, the authors showed that the distribution of Guess 2 reports was not modulated by which item was provided as guess one. This led to the conclusion that the temporal position of the selection window does not vary considerably across trials and that both guesses are sampled from the *same* probability distribution/ pattern of activation (Vul et al., 2009).

Notably, as previously discussed, Zivony and Eimer (2021) showed that the latency of the N2pc varies with the type of conjunction observed, e.g. the N2pc’s onset latency was significantly delayed in post-target distractor intrusion trials compared to correct response trials. This stands against the simple interpretation of Vul et al.’s finding that the proposed distribution of potential responses is unchanged across trials. If attentional selection (i.e. the N2pc) varies in time, one would expect the positions (relative to the target) included in the response distribution to vary accordingly across trials. This though, leaves the possibility that there is always a distribution, but that the positions included in this distribution (relative to the target) vary across trials.

Next, we will demonstrate that the 2-feature-ST^2^ model accounts for distractor intrusion errors in RSVP. After a presentation of the model’s architecture & dynamics, introducing how our model generates intrusion errors, human response distributions and reaction times, we will explain how electrophysiological (i.e., EEG) data will be replicated. We will first replicate the early work in this field (Botella, 1992; Botella et al., 2001; Chennu et al., 2011) before replicating the recent data of Zivony and Eimer (2021), who compared correct reports with post-target (i.e., +1) intrusion errors. Finally, we show that the evidence Vul et al. (2009) provide for responses being samples from a probability distribution/ pattern of activation across potential responses can also be simulated with the 2-feature-ST^2^ model.

### The 2-feature Simultaneous Type/ Serial Token Neural Model

The 2f-ST^2^ model has previously been presented in a preliminary form (Chennu et al., 2011). The present paper will introduce the current, updated version of our computational model. At certain points, we will refer to the previous version to highlight important differences. When we do this, we will refer to the preliminary version (Chennu et al., 2011) as the “initial 2f-ST^2^ model”.

### Architecture

The architecture and dynamics of the 2f-ST^2^ model (Figure 5) closely mirror that of the original ST^2^ model (Bowman & Wyble, 2007), which we reviewed in section “Simultaneous Type/ Serial Token Models (Bowman & Wyble, 2007)”, again implementing a parallel Stage 1 extracting and processing types; a sequential Stage 2 binding types to tokens; and a blaster circuit. The 2f-ST^2^ model extends the original model’s architecture (Bowman & Wyble, 2007), adopting two pathways in its Stage 1, enabling it to simultaneously extract two kinds of types, corresponding to key and response feature dimensions, for each item in the RSVP stream (see Figure 5A). Stage 2 is extended in a similar fashion, now consisting of two binding pools, one for each feature dimension, which allows a working-memory token to be associated with a pair of types indicating the encoded item’s key (e.g., colour) and response (e.g., shape) feature. Types and tokens are fundamental concepts in the 2f-ST^2^ model as well as in its predecessor models (Bowman & Wyble, 2007; Wyble et al., 2009); see also Kanwisher (1987).

**Figure 5.**
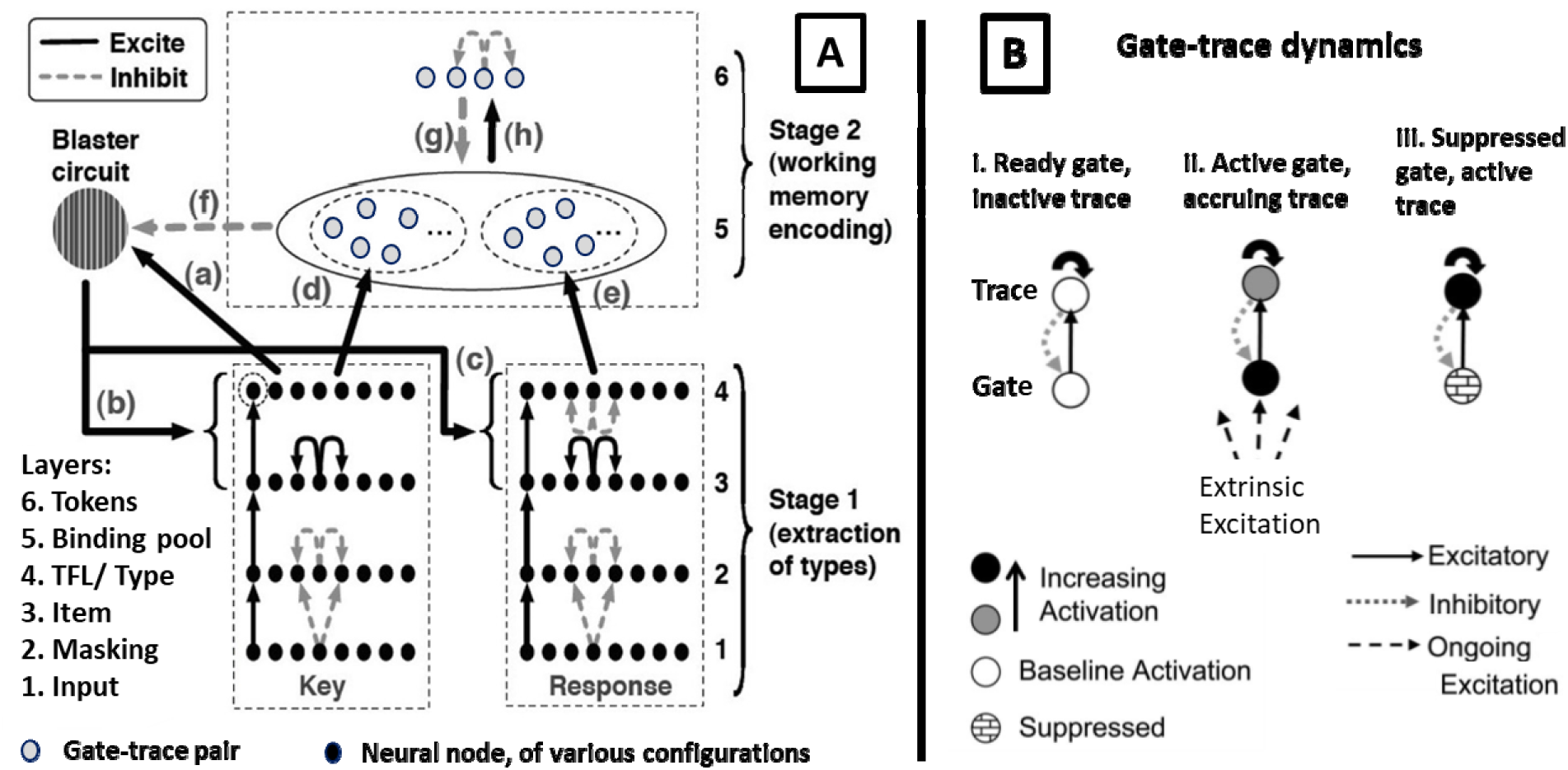
2f-ST^2^ model. [A] The 2f-ST^2^ Architecture showing two Stage 1 pathways, Blaster Circuit, Binding Pool and Tokens. Stage 1 implements a key and a response pathway in which respective feature dimensions are extracted and processed. Task-demand picks out task-relevant neurons, see dashed circle around leftmost unit in final layer of key pathway. The key type meeting task-demands (unit in dashed circle becoming active) triggers the blaster circuit in the TFL/ type layer of the key pathway. The blaster enhances activation levels of both pathway’s item and TFL/ type layers. Finally, the currently active token is bound to a pair of key and response types in Stage 2, meaning that the item was encoded in WM. Adapted from Chennu et al. (2011). TFL denotes Task Filtered Layer. [B] Trace-gate mechanism: binding pool and tokens (panel [A]) are gate-trace pairs, which are initially in a “ready” state (i), can be externally excited (ii), and then enter an allocated state (iii). Adapted from Bowman & Wyble (2007).

The key pathway extracts items’ key features and hence conceptually resembles Botella et al.’s (2001) Module K. Importantly, the RSVP stream’s task demands (dashed circle around the left key TFL/ type node in Figure 5A) are implemented in this pathway, specifically in its TFL/ type layer (where TFL denotes Task Filtered Layer). The item that carries the task-relevant key feature (e.g., the colour red in Figure 1) activates the blaster circuit through the connection (a) of Figure 5A. Therefore, distractors are modelled as items that lack the key-feature demanded by a given task (e.g. corresponding to black or yellow items) and hence do not activate the blaster circuit. Once the blaster is activated, similarly to the original ST^2^ model (Bowman & Wyble, 2007), nodes of the last two Stage 1 layers (item and TFL/ type) are enhanced. Importantly, in the 2f-ST^2^ model, the blaster enhances key and response item and TFL/ type layers simultaneously and to the same extent. This is illustrated by connections (b) & (c) in Figure 5A. The blaster enhancing later Stage 1 activation levels resembles the TAE processes considered in Zivony and Eimer (2020, 2021). Similar to the original ST^2^ model (Bowman & Wyble, 2007), the blaster’s enhancement means that the most strongly active node^2^ of each pathway’s TFL/ type layer can become sufficiently active, that they cross an *effective* access threshold and initiate activation in their respective binding pools; see Figure 5A connections (d) and (e).

Once respective binding pool nodes have been sufficiently excited, successful binding between one key type, one response type and one token occurs. The TFL/ type layer response pathway dynamics depend on whether Botella et al.’s (2001) or Zivony and Eimer’s (2020, 2021) experimental paradigm are to be modelled. Specifically, all response types can be bound to the token via their respective binding pool units when replicating Botella et al.’s (2001) experimental paradigm. However, when modelling Zivony and Eimer’s (2020, 2021) paradigm, only the target and the +1-distractor can be active in this layer, as the remaining items are target category-nonmatching distractors. In both cases, lateral inhibition between category-matching items (i.e., all items in Botella et al.’s (2001) paradigm and the target & +1-distractor in Zivony and Eimer’s (2020, 2021) paradigm) in the response TFL/ type layer (see lateral inhibition in right hand Layer 4) generates a competition between types to excite their respective binding pool units. The target versus intruder competition examined by Zivony and Eimer (2020) can be seen to be consistent with this aspect of the 2f-ST^2^ model. Which response type is ultimately bound to the currently active token therefore depends on which TFL/ type response node was active the most at the time the blaster enhances late Stage 1 layers. This mechanism further enables the 2f-ST^2^ model to commit distractor intrusion errors, which occur whenever a distractor item’s response type wins this pathway’s competition (explained in detail in the next section).

Tokens in the 2f-ST^2^ model largely work as they did in the original ST^2^ model (Bowman & Wyble, 2007). Specifically, the currently active token inhibits the other tokens’ binding pool nodes whenever the model is encoding (Figure 5A’s connection (g)) and is excited by one key and one response binding pool node during binding/ tokenization (Figure 5A’s connection (h)). The successful binding of a given token to a pair of key and response types thus allows the 2f-ST^2^ model to store episodic, i.e., ordered, representations of items and hence constitutes WM encoding in the model.

Furthermore, as in previous ST^2^ models (Bowman & Wyble, 2007; Wyble et al., 2009), the 2f-ST^2^ model implements gate-trace node-pairs; see Figure 5B. Gate (on) nodes (or neurons) excite trace (off) nodes after being externally excited themselves. Trace nodes in turn accrue the excitation they receive from gate nodes and, after crossing a threshold, inhibit the gate node strongly, suppressing all gate node activation and placing the unit in an allocated state. Trace nodes are important in indicating that an event has terminated, for example that a given token has completed binding to a pair of key and response types.

For an overview of the 2f-ST^2^ model’s key terms and their definitions see the glossary in Appendix 3.

### Parameter Modifications

Large neural network models such as neural-STST exhibit very substantial fitting degrees of freedom, potentially making it possible to simulate any data pattern (Roberts & Pashler, 2000). To limit this flexibility, as with previous work with the STST model, we have endeavoured to preserve the parameter settings of the original model. However, adding a second parallel pathway to the model, with associated adjustments to lateral inhibition and task demand has required us to re-stabilise the model and accordingly adjust some parameters. These changes to Stage 1 and Stage 2 parameters are documented in Appendix 1.

### Dynamics

Task demands were absent in the Response pathway TFL when modelling experiments in which all stimuli could have been targets (as in Botella’s paradigms (Botella et al., 2001; Botella & Eriksen, 1992)). However, a task filter is implemented in the response pathway TFL/ type layer when modelling Zivony and Eimer’s (2021) experiment in which only two stimuli met task demands (i.e., the two digits adjacent in time in Figure 13). To this end, the 0 and +1 response types, modelling the target and following distractor, met task demands and thus had different response task demand unit (TDU) values from the other types^3^. TDUs model task relevance of stimuli by adding a bias to respective TFL/ type layer neurons. Across both pathways, distractor stimuli’s TDUs bias corresponding TFL/ type layer units negatively with a bias of -.3. Response pathway TDUs that met task demands (i.e., 0-& +1-types when modelling Zivony and Eimer’s (2021) experiment) do *not* bias their TFL/ type layer neurons (bias = 0). In the key pathway, task-relevant types had TDU biases of +.003, thus positively biasing their respective TFL/ type layer neuron. However, due to the lateral inhibition present in the response pathway TFL/ type layer, a lower bias value of task-relevant response TDUs was required compared to key TDUs.

To simulate the generation of correct reports and intrusion errors, input-strengths varied across simulation trials for target key features, but not for distractor key features. Variability in target key feature input-strengths across trials induces variability in blaster firing latencies and encoding times. Response feature input strengths varied similarly across trials for the target as well as its proximal distractors (2 positions in either direction, i.e., −2 to +2) and remained fixed for all other distractors.

### Configuration

For the model’s configuration, there are three important delay parameters: τD, τK and τR. τD (random delay) is sampled for each item in an RSVP stream, when it enters the model at the input layer, following a Gaussian distribution with a mean of zero and a standard deviation of 4 time-steps (∼20 ms-equivalent). τK (key delay) and τR (response delay) add a positive or negative time-delay to the processing of the key and response pathway, respectively. Importantly, τK and τR are set when running the model and *remain fixed throughout a complete simulation run of the model.* Both parameters’ ‘default’ value is 0 and changing them enables the replication of a variety of experiments involving manipulations that affect the speed of key or response feature processing.

To obtain more trials than in previous ST^2^ simulations (Bowman & Wyble, 2007; Chennu et al., 2011; Wyble et al., 2009), we ran the model five times for all configurations, re-seeding the random number generator (rng) before each run. However, whenever we re-generated results already presented with the initial 2f-ST^2^ model, we only ran the model once (using the rng seed employed in (Chennu et al., 2011)) to ensure comparability. We also only ran the model once when presenting individual neurons’ traces (e.g., membrane potentials) in Appendix 4, as saving these drastically increases computational load and was not feasible with five runs.

### Intrusion Errors in the 2f-ST^2^ model

We provide a schematic illustration of how the 2f-ST^2^ model generates correct and intrusion responses in Figure 6. Time progresses from top to bottom panels and the left and right columns illustrate how correct and intrusion reports are generated, respectively. The model’s key and response pathways depict individual key and response features, respectively, with key features being represented by black, red, and yellow ellipses and response features being represented by black rectangles labelled B, S, and F. Importantly, we used gradients to represent activation strength of an item’s representation in the model’s hierarchy at a given point in time (throughout a trial), with more transparent gradients meaning lower activation strengths. For example, the small black oval in the second row of Figure 6’s left column is more solid towards its top and more transparent in its bottom area, which means that the black item’s key feature has a rather strong activation in the key TFL/ type layer, but an already decaying activation in the previous (lower) key item layer.

**Figure 6.**
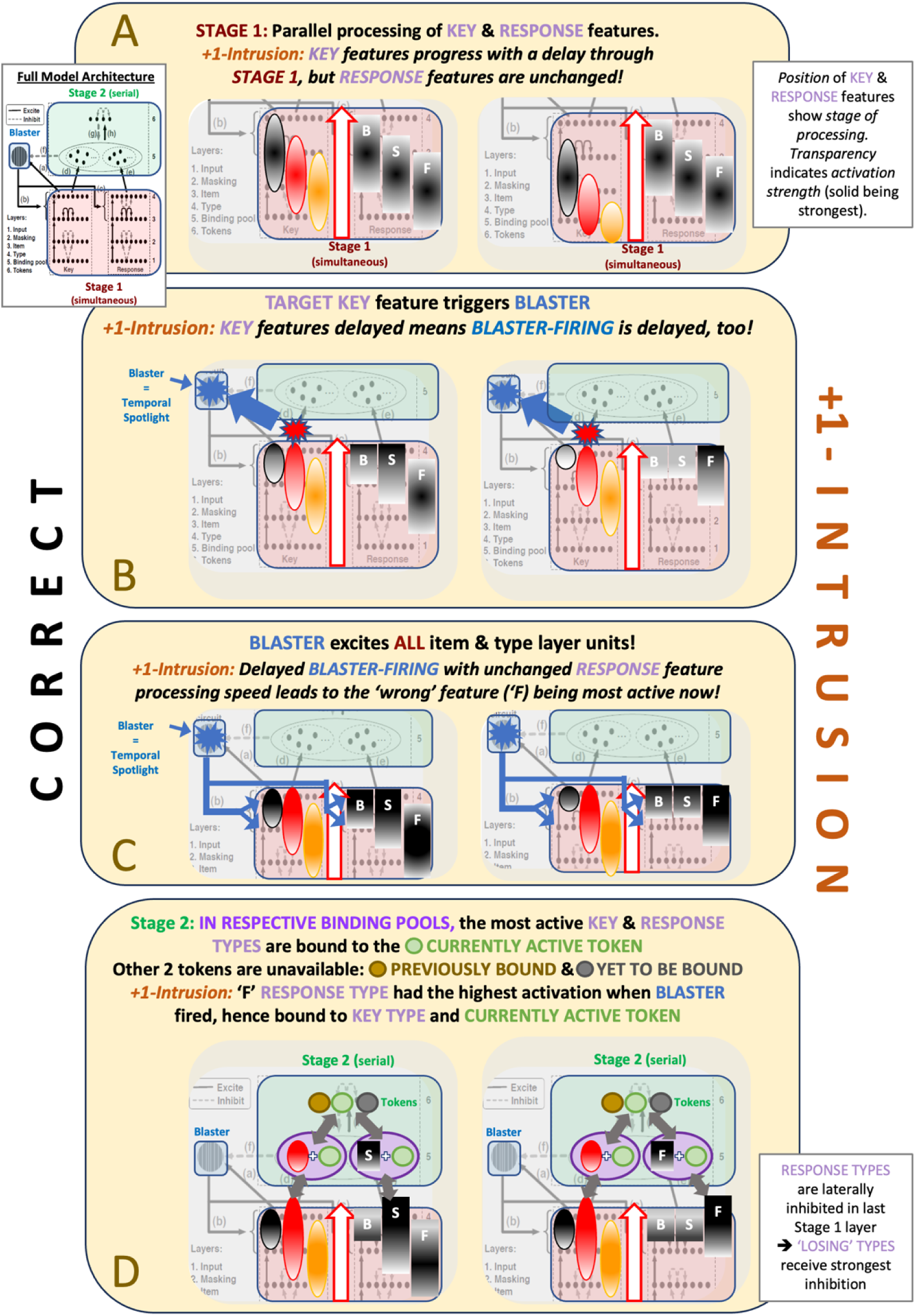
Schematic outline of how the 2f-ST^2^ model generates correct (left column) and +1-intrusion (right column) responses. An overview of the complete model architecture is provided as an inset. Key features are represented by black, red, and yellow ellipses and response features by black rectangles labelled B, S, and F. Panel A illustrates parallel propagation through Stage 1 layers, being typically delayed for the key pathway in (post-target) intrusion trials. Panel B illustrates how this translates to the intruder (F) type being the most active at the time-point of blaster-firing. Panel C illustrates how the blaster enhances *all types* in later Stage 1 layers and how this leads to different types crossing the threshold to the response binding pool. Panel D illustrates how, as a result, different binding pool units are bound to the currently active token (green), leading to a correct response on the left and to an intrusion response on the right side. Panel D also illustrates how response types are laterally inhibited by each other in the last Stage 1 layer (TFL/ type). Lateral inhibition suppresses ‘losing’ (i.e., unbound) types.

The right column shows a +1-intrusion response, i.e., the distractor stimulus presented *immediately after* the target was reported. Concretely, we show the example of coloured letters being presented in the RSVP stream, with a red S as the target (key-feature in left pathway = red ellipse & response-feature in right pathway = S-shape) and a yellow F as the +1-intruder (key-feature = yellow ellipse & response-feature = F-shape). The intrusion is due to delayed key-feature processing, which means the blaster is triggered later (right column, second row of Figure 6), when the F-type of the response pathway is most active (i.e. F rectangle is darker). In contrast, for correct reports (left column), the correct S-type had the highest activation. The bottom two rows show how this difference in response-type activation strengths translates to either one response type (correct-S, left, or intruder-F, right) being bound to the active token and hence encoded into WM. The third row illustrates how the blasters’ enhancement strengthens the activation of *all types* (i.e. *all* gradients get more solid) in the last two Stage 1 layers, leading the *strongest TFL/ type* (last Stage 1 layer) unit to cross the threshold to binding pool entry. The last row illustrates how the corresponding binding pool units are bound to the token, generating a correct and intrusion response in left and right columns, respectively. As shown, binding pool units have a conjunctive tuning, responding to the conjunction of a type and a token. Figure 6’s bottom row further demonstrates that neurons in the response TFL/ type layer laterally inhibit each other. Naturally, the response TFL/ type neuron that has the strongest activation when the blaster fires will laterally inhibit the other neurons in this layer more strongly, leading to the ‘competition’ to enter the response binding pool to have an unambiguous outcome (as illustrated by the rather large contrast in gradients between response TFL/ type neurons at this final stage).

The timing of blaster enhancement with respect to the activation of response units is then critical in the 2f-ST^2^ model. The key and response types that are most active when the blaster fires will usually be bound to the currently active token. τK and τR, the fixed delays to key and response pathways, impact model dynamics by modulating the distribution of activation strengths across response TFL/ type layer neurons, and particularly the most active neuron, at the time of blaster firing. τK accomplishes this by changing blaster firing latency, e.g. a delay in key pathway processing means that the task-relevant key TFL/ type neuron excites the blaster later. τR on the other hand modulates the distribution of activation strengths across response TFL/ type neurons and the neuron most active when the blaster fires. For example, a positive τR value means that response pathway processing commences *later*. Given a fixed blaster-firing latency, delayed response pathway processing would imply that *pre-target* intrusions are more likely.

This is because, compared to a τR of 0, the RSVP stream’s response features had less time to propagate to the later Stage 1 layers of the response pathway and, thus, it is more likely that pre-target TFL/ type neurons are strongly active when the blaster fires. Additionally, τK & τR enable us to adjust key and response feature salience, which are often manipulated in experiments. That is, a very salient key or response feature is argued to be processed easily and swiftly by participants, which corresponds to a low (or negative) value in the respective pathway’s delay.

### Virtual Event-Related Components (vERPs) of the 2f-ST^2^ model

The 2f-ST^2^ model can generate virtual ERP (vERP) components, enabling comparison to human ERPs. vERPs are based on the postsynaptic activation potentials of virtual neurons, which are the sum of a neuron’s presynaptic activation multiplied by corresponding weights. The virtual N2pc (vN2pc) component is computed as the postsynaptic activation of the blaster’s output neuron. The virtual P3 (vP3) is computed as a sum across four layers. Both pathways’ TFL/ type layers, as well as binding units’ gate projections are included in the vP3 and the contribution of all four layers is equal; see Figure 5A. In comparison with previous ST^2^ modelling of virtual P3s (Chennu et al., 2009; Craston et al., 2008), we excluded the item layers from contributing to the P3. This reflects the TFL/ type layers and binding pool most naturally corresponding to the later cognitive processes related to working-memory encoding, which are typically argued to underlie human P3s (Akyürek et al., 2010; Polich, 2007), although see Jones et al. (2020) and Pincham et al. (2016) for a linking of the (breakthrough) P3 to conscious perception. The item layer, on the other hand, reflects cognitive processes that are prior to encoding and hence were excluded.

In previous iterations of ST^2^ models (Chennu et al., 2009; Craston et al., 2008), vERPs have been defined as the amplitude of postsynaptic potentials of virtual neurons across a given layer. However, the human EEG signal does not exactly, or exclusively, capture postsynaptic potentials. Rather, EEG measures electrical dipoles, which are generated by micro-current sources at cell membranes (Nunez & Srinivasan, 2007). The current is affected by sources of mixed signs, e.g., synchronous postsynaptic potentials, local inhibitory synapses as well as passive membrane (return) currents needed for current conservation (Nunez & Srinivasan, 2007). To more accurately reflect that EEG signals typically measure electrical dipoles, we developed an alternative to the original (layer-wide postsynaptic activation of virtual neurons) ST^2^-vERPs. This alternative was computed by first calculating the approximate derivative of original vERPs (i.e. the difference of adjacent points in the time-series), since (modulo a multiplicative constant, the capacitance) current is the time derivative of the voltage, i.e., how it changes through time, before 1-D smoothing the resulting time-series with a gaussian kernel.

These smoothed vERPs carry two advantages. First, they more closely resemble the current recorded by EEG electrodes and, second, they capture important properties implied by the EEG signal being based on dipoles, e.g., that a polarity-reversal (i.e., a negativity following a positivity or vice versa) implies a shift in the direction of net ion-flow. Nonetheless, the original vERPs reflect relevant internal processes and dynamics of the 2f-ST^2^ model. Hence, we will consistently present the original (unsmoothed) vERPs (reflecting post-synaptic potentials) as well as the smoothed difference vERPs side-by-side. See Appendix 2 for the complete procedure involved in smoothing vERPs.

### The 2f-ST^2^ Model replicating Human Data

This section will commence with 2f-ST^2^ replications of early work by Botella and colleagues (Botella, 1992; Botella et al., 2001) as well as Chennu et al. (2011), in which all stimuli shared the same response feature category (e.g., all being digits). We will then replicate the recent work of Zivony and Eimer (2021), in which only the target and the +1-intruder item shared the same response feature category. This range of experiments will be replicated with the same model architecture (Figure 5A), with the addition of response pathway task filtering when necessary (i.e., when replicating Zivony and Eimer’s (2021) experiments) and configuring τK and τR to match empirical manipulations that affected key or response feature processing speeds, where one time-step of τK or τR approximates 5 ms. This section will conclude with replicating the results of Vul et al. (2009) who asked participants to make multiple responses after each trial. Importantly, following the method of the ST^2^ model (Bowman & Wyble, 2007), we have endeavoured to simulate empirical data without fitting core model parameters apart from processing speed of the different pathways. Modulo the cases that we explicitly discuss, we will employ this same strategy in this paper. As a result, it is not realistic to believe that the model will produce quantitively precise fits to the data. Accordingly, the criteria for judging our data fits should be qualitative; i.e. do our simulations qualitatively replicate empirical phenomena?

### Experiments with response feature category matching distractors

For our first simulations, no task-filtering was applied to the response TFL/ type layer of the 2f-ST^2^ model, since in these Botella et al. (2001) tasks, all response features were in principle, able to be bound to the currently available token. Results were obtained after a single run of the model, using the same rng seed as in the initial 2f-ST^2^ model (Chennu et al., 2011) to ensure comparability. Further, we will use a measure called the Average Position of Intrusions (API, Botella et al., 2001) to demonstrate the qualitative fit between the 2f-ST^2^ model and empirical data. The API measure is the sum across all intrusion positions with respect to the target multiplied by the proportion of a given intrusion with respect to all intrusion trials (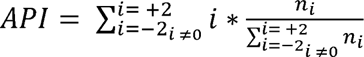 where *i* is distractor position and *n_i_* the number of responses at position *i*). More precisely, API is the expectation with respect to the probability distribution of responses across intrusion positions. Thus, the API quantifies the central tendency of a response distribution, thereby indicating shifts in response tendency that may result from empirical manipulations in human experiments or different values of τK and τR in the 2f-ST^2^ model.

### Replicating Botella et al.’s (2001) response distributions

Botella et al. (2001) employed different levels of word-frequency to modulate key or response feature processing speeds. An example stream of Botella et al.’s (2001) Experiment 1A is presented in Figure 7. Participants’ task was to report the colour of the stream’s animal word. High-frequency (HF) animal words (e.g., dog) and low-frequency (LF) ones (e.g., iguana) modulated key feature processing speed, since HF words are detected faster as belonging to the animal category. Botella et al. (2001) observed a pre-target shift of intrusion responses going from LF to HF words. We replicated Botella et al.’s (2001) Experiment 1A (Figure 9A) setting τK = 0 / τR = 0 for HF and τK = 8 / τR = 0 for LF. A larger τK value means a delayed onset of key pathway processing in the 2f-ST^2^ model, simulating that the *identity* of the animal word (*key* feature) was detected slower when it was less frequent.

**Figure 7.**
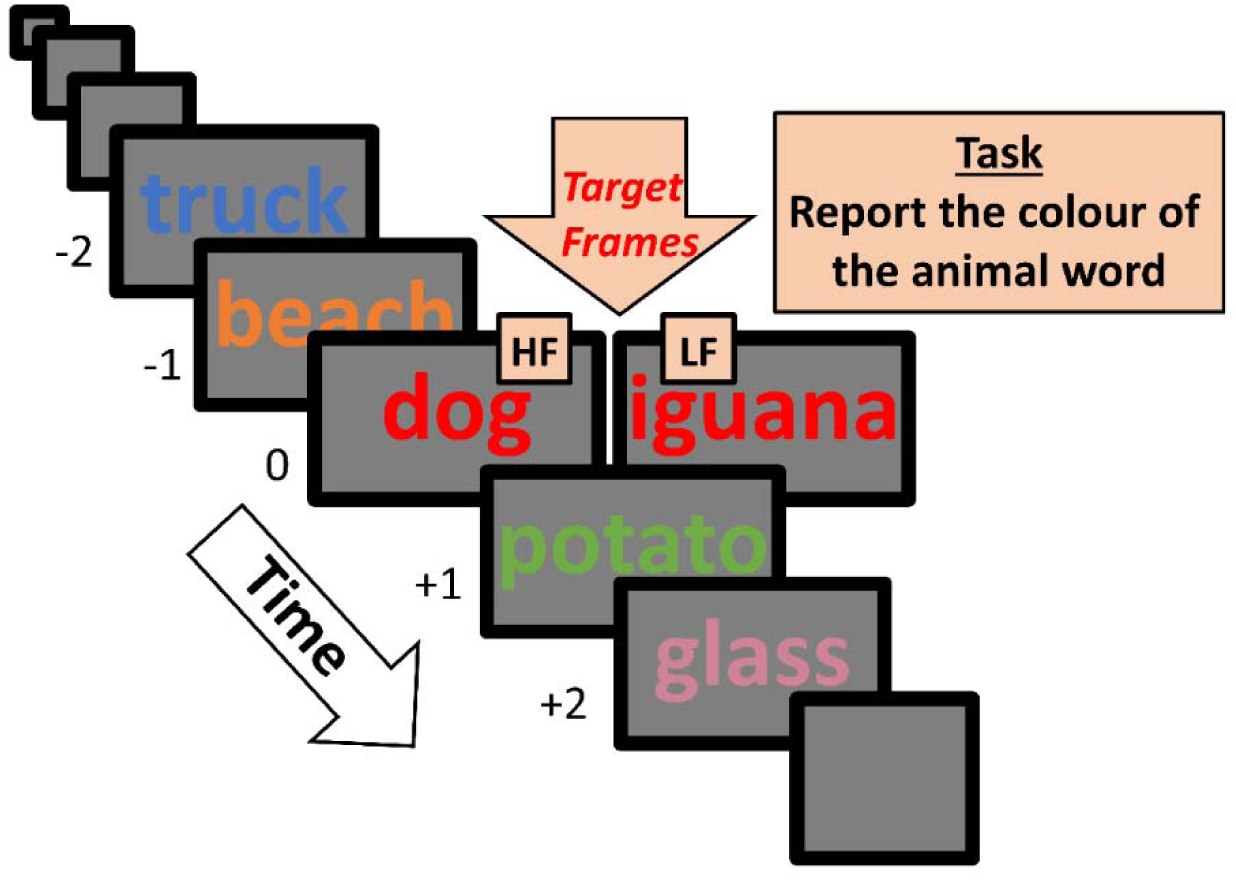
Experimental design of Botella et al.’s (2001) Experiment 1A. Plotting conventions are identical to those of Figure 1. Participants were asked to report the colour of the stream’s animal word. Key feature processing speed was modulated, since high-frequency (HF) animal words (e.g., dog) are detected faster than low-frequency (LF) ones (e.g., iguana). Botella et al. (2001) observed a pre-target intrusion response shift going from the LF to the HF condition.

Figure 8 presents an example stream of Botella et al.’s (2001) Experiment 2. Participants were asked to report the uppercase word, which again were high-(TABLE in Figure 8, HF) or low-frequency words (PRATE in Figure 8, LF). This modulates response feature processing speeds, since the key feature is fixed (uppercase-ness), while HF words are reported faster than LF ones. Botella et al. (2001) observed a post-target shift of intrusion errors going from LF to HF. We replicated Botella et al.’s (2001) Experiment 2 (Figure 9B) setting τK = 0 / τR = 0 to resemble the HF and τK = 0 / τR = 2 to resemble the LF condition. A larger τR value means a delayed onset of response pathway processing, consistent with the *identity* of the uppercase word (*response* feature) being processed slower (i.e., WM encoding later) when it was less frequent. A LF to HF post-target intrusion shift is in line with the 2f-ST^2^ model (see Figure 6), since HF response types will progress earlier through the model’s response pathway. Therefore, at the fixed time of blaster firing (since the key feature (uppercase-ness) is held constant), it is likely that early response TFL/ type neurons’ have already decayed in activation strength and subsequent, e.g., post-target intruder, TFL/ type neurons have the highest activation and are thus bound to the currently active token at Stage 2.

**Figure 8.**
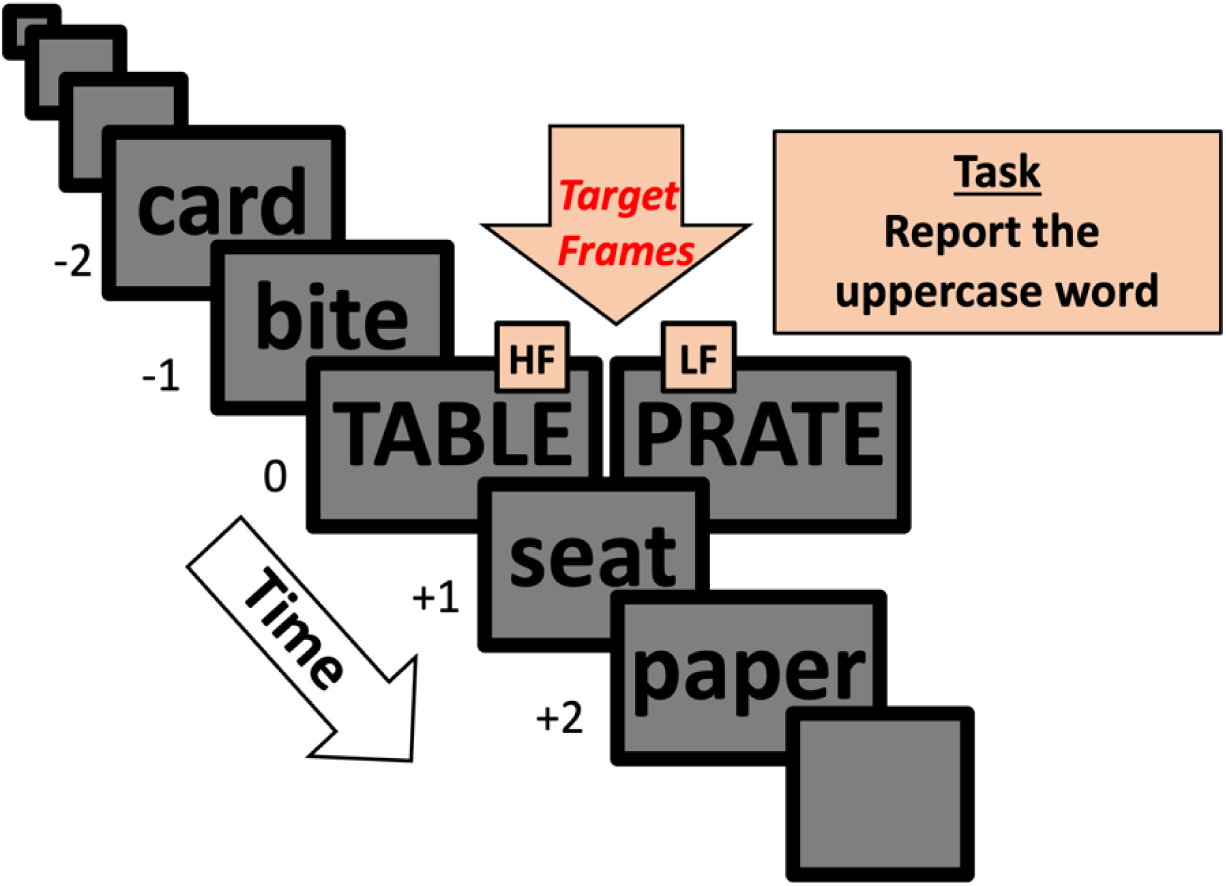
Experimental design of Botella et al.’s (2001) Experiment 2. Plotting conventions are identical to those of Figure 1. Participants were asked to report the uppercase word. Response feature processing speed was modulated, since high-frequency (HF) uppercase words (e.g., TABLE) are reported faster than low-frequency (LF) ones (e.g., PRATE). Botella et al. (2001) observed a post-target response shift in the HF condition.

Figure 9 illustrates how the 2f-ST^2^ model qualitatively replicates Botella et al.’s (2001) Experiments 1A (Figure 9A) and 2 (Figure 9B). Experiment 1A is replicated since our model generates the same increase in *pre-target intrusions* as more frequent animal words are used as the key feature (larger teal than dark blue bars in Figure 9A*’s left* column). This pre-target shift was accompanied with empirical API values of −0.306 (HF) and 0.043 (LF) (Botella et al., 2001), which was replicated by the 2f-ST^2^ model with API values of −0.11 (HF, τK = 0 / τR = 0) and 0.69 (LF, τK = 8 / τR = 0). Experiment 2 is replicated since our model generates the same increase in *post-target intrusions* as more frequent uppercase words are used as the response feature (larger teal than dark blue bars in Figure 9B’*s right* column). The empirical API values of Botella et al.’s (2001) Experiment 2 were 0.131 (HF) and −0.034 (LF), which were again replicated by the 2f-ST^2^ model: −0.11 (HF, τK = 0 / τR = 0) and −0.45 (LF, τK = 0 / τR = 2).

**Figure 9.**
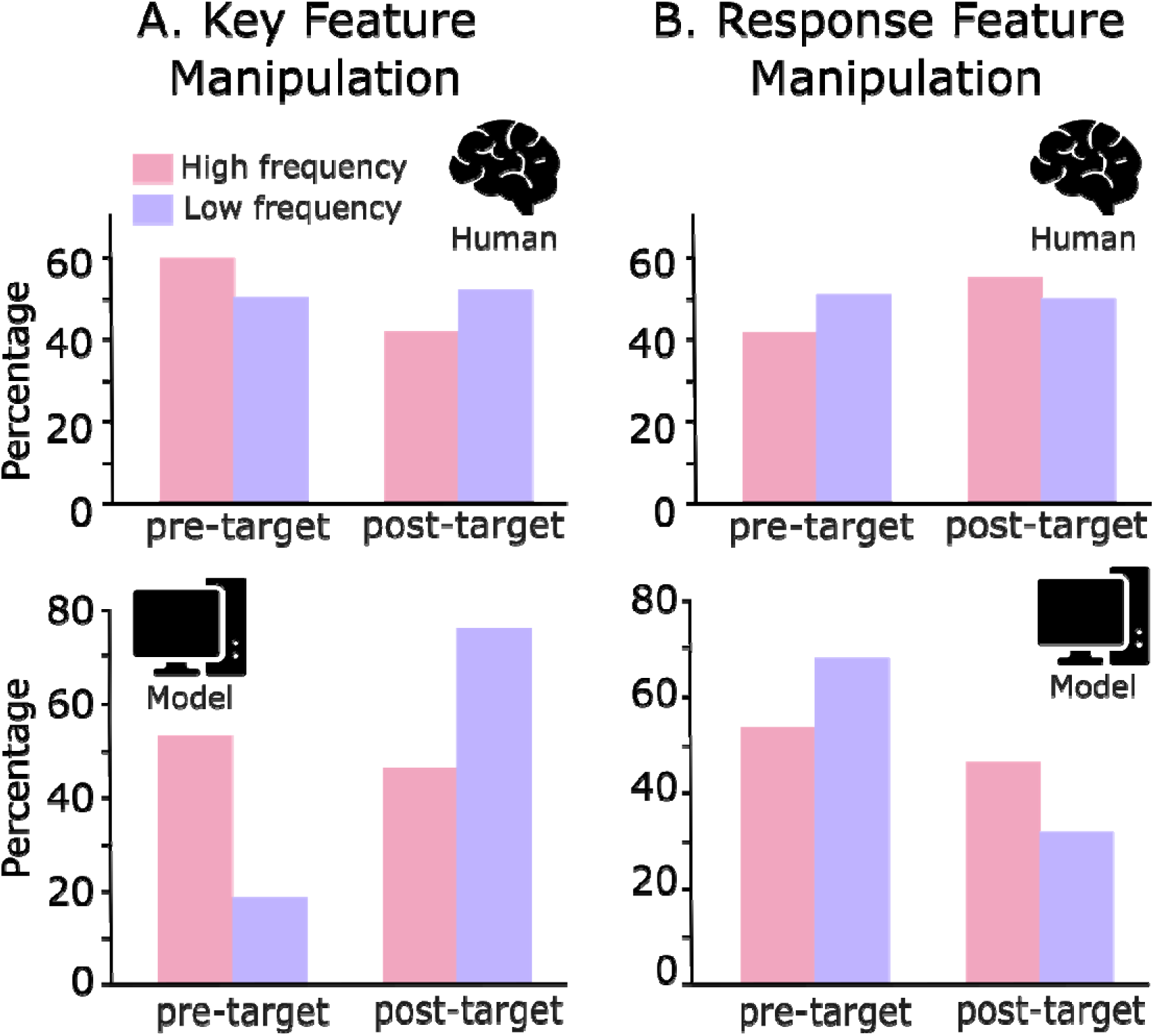
The 2f-ST^2^ model replicates the response distributions obtained in Botella et al.’s (2001) Experiments 1A and 2. Botella et al. (2001) varied word frequency to obtain an isolated modulation of key feature processing speeds in Experiment 1A (Panel A), which was accompanied with an increase in pre-target intrusion responses for more salient key features (higher frequency words). The 2f-ST^2^ model replicates this pattern (Panel A). Botella et al. (2001) modulated word frequency to obtain an isolated modulation of response feature processing speeds in Experiment 2 (Panel B). This was accompanied with an increase in post-target intrusion responses for more salient response features (higher frequency words). This figure is adapted from Chennu et al. (2011).

### Replicating Chennu et al.’s (2011) Fortunate Conjunctions Experiment

Chennu et al.’s (2011) fortunate conjunctions experiment presented coloured letter or symbol targets within RSVP streams of coloured digit distractors (SOA 94 ms). The key feature was target category (letter or symbol), and the response feature was colour; for details, see (Chennu et al., 2011). Importantly, in this experiment, key feature processing speed was manipulated since symbol targets were detected slower than letter targets. This resulted in a post-target shift of intrusions, generating API values of 0.18 (letter targets) and 0.31 (symbol targets). The 2f-ST^2^ model replicates this post-target shift with APIs of −0.97 (early key condition (EK), τK = −8 / τR = 0) and −0.11 (late key condition (LK), τK = 0 / τR = 0). Figure 10 illustrates that the 2f-ST^2^ model replicates these empirical results.

**Figure 10.**
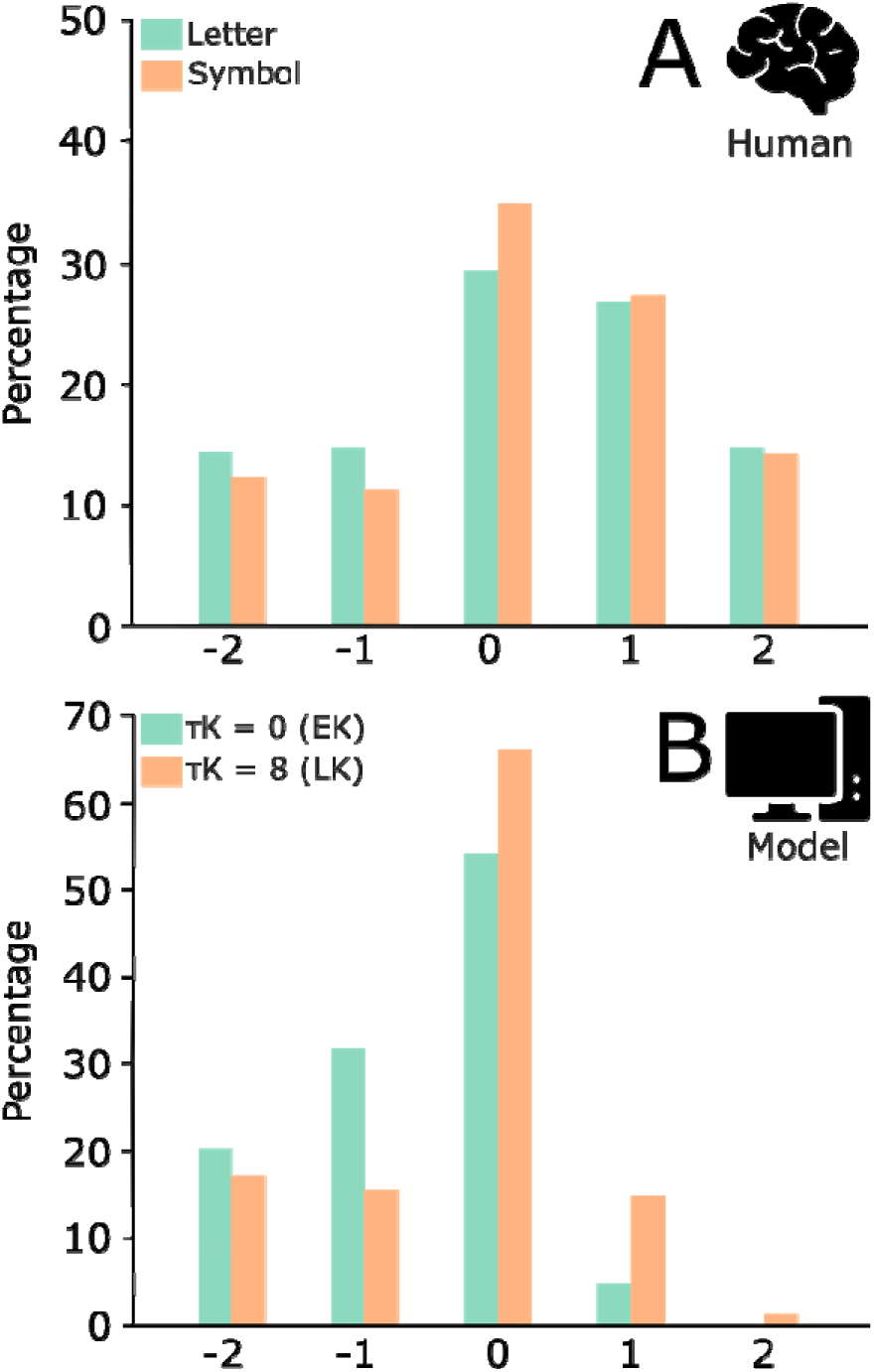
Chennu et al.’s (2011) fortunate conjunction experiment (Panel A) replicated with the 2f-ST^2^ model (Panel B). Letter and symbol experimental conditions were replicated with τKs of 0 & 8 time-steps (corresponding to early key (EK) & late key (LK)), respectively. This figure is adapted from Chennu et al. (2011) to reflect the current 2f-ST^2^ model’s behaviour in Panel B.

### Counterintuitive Reaction Time (RT) Pattern of Botella (1992)

As previously discussed, a central finding, and a critical test of the 2f-ST^2^ model, is a rather counterintuitive pattern of reaction times (RT) presented by Botella (1992). In this study (Botella, 1992), participants had to give a response as soon as the target stimulus was perceived. The central finding was that correct reports were provided the fastest, followed by pre-target errors and then post-target errors. This finding is striking and counterintuitive, since one would expect pre-target errors to be made quicker than correct reports, because the item that was reported by the participant was presented *earlier in the RSVP sequence than the correct item*.

Thus, the results of Botella (1992) suggested that the pattern of RTs in distractor intrusion experiments does not follow the sequence of item presentation in the RSVP streams. We modelled RTs in the 2f-ST^2^ model as the point in time at which the currently available token completes binding to the key as well as response neuron. An index of this event is provided by the currently active token’s trace neuron (Bowman & Wyble, 2007). Specifically, (as already used in Chennu et al. (2011)) the model provides its response in a given trial if the token’s trace neuron reaches 75% of its maximum amplitude.

Of course, a response to a stimulus, in fact, results from a processing hierarchy in which WM-encoding is an intermediate process that follows perceptual processes. In the 2f-ST^2^ model, the processes required for a response that follow WM-encoding (e.g. excitation of neural populations in motor-cortex) were not modelled. Nonetheless, the 2f-ST^2^ model replicates the counterintuitive pattern of RTs shown by Botella (1992); see Figure 11. Our model yields responses that are over 100 ms-equivalents later than the human data. However, as previously discussed, this is not surprising, since we maintained the parameter space of the original ST^2^ model, which was recognized to have delayed responses compared to human RSVP data (Bowman et al., 2008; Bowman & Wyble, 2007). The 2f-ST^2^ model moreover not only replicates Botella’s (1992) RT patterns qualitatively but also provides an explanation of why this pattern was observed in humans with its vERP components. This will be presented next.

**Figure 11.**
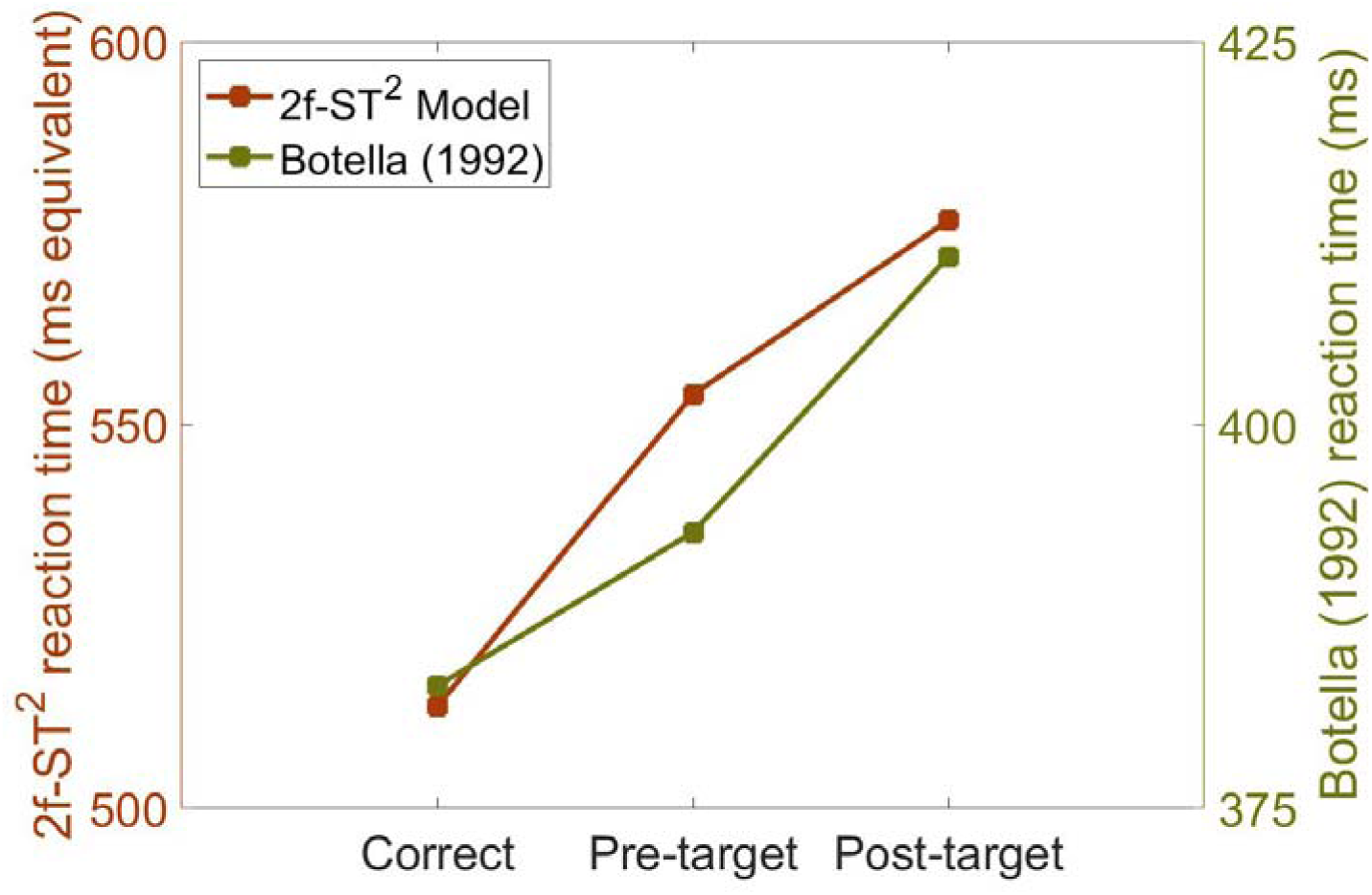
Reaction time patterns of Botella (1992) replicated by the 2f-ST^2^ model. Note that y-axes are different.

### 2f-ST2’s virtual ERPs explaining RT patterns

The counterintuitive pattern of reaction times presented in Figure 11 can be understood by looking at our model’s virtual ERP components. Figure 12 shows the virtual N2pc and P3 components. Green, black, and red traces correspond to pre-target intrusion, correct response, and post-target intrusion trials, respectively. Vertical coloured lines in all vERP plots indicate the *average* latency of blaster-firing for each response condition, with their exact numerical values in the legends. Blaster-firing latencies were defined, at the single-trial level, as the time point at which 50% of the area under the vN2pc (measuring the blaster’s output) curve was reached. The vP3 pattern is especially relevant to reaction times (RT), as we model RTs as the time-point at which the binding of a type to a token completes in the binding pool, a process that is generated by the layers that constitute the vP3. As displayed in Figure 12B, correct trials show an earlier vP3 than pre-as well as post-target intrusion trials, qualitatively fitting the reaction time patterns well. This can be explained as follows.

**Figure 12.**
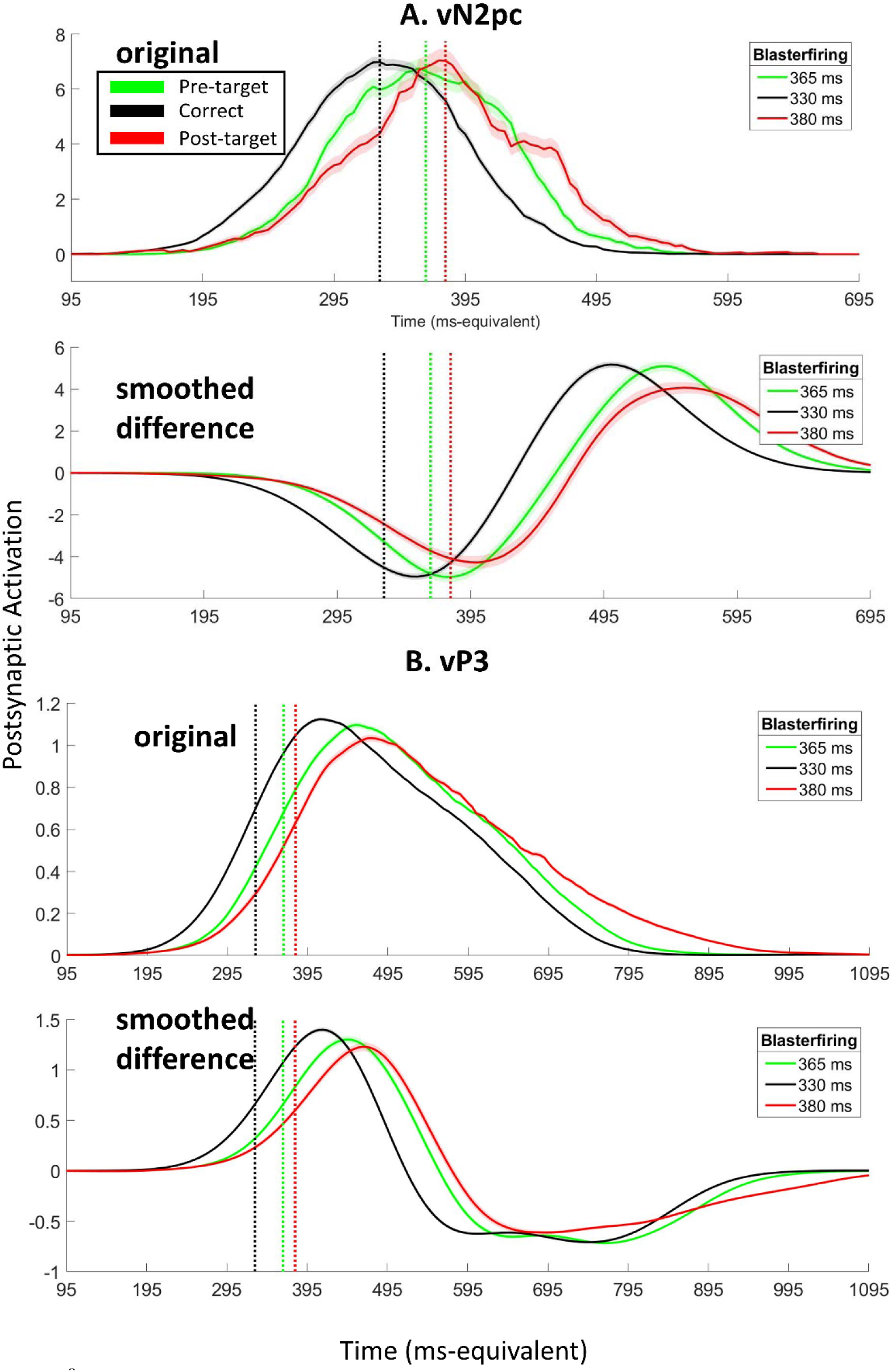
2f-ST^2^’s vERPs without response pathway task-filtering and no processing delays to either pathway explaining RT findings of Figure 11. Green, black, and red traces correspond to pre-target, correct, and post-target response conditions, respectively. Note that intrusion conditions collapse across +/- 1/2 intrusions, hence, the green trace corresponds to −2 as well as −1 intrusion trials. We present the vN2pc in Panel A and the vP3 in Panel B, plotting the original (model-output) vERPs as well as the smoothed difference vERPs. Vertical lines indicate *average* blaster-firing (with their numerical values being shown in the legend on the right).

In correct trials, the blaster fires close in time to the activation peak of a strongly active target response TFL/ type neuron. Consequently, the target response TFL/ type neuron quickly wins the competition with distractor response TFL/ type neurons, which arises from the lateral inhibition present in this layer (see Figure 5A, grey dashed arrows, 4^th^ layer, Stage 1, response pathway and Figure 6, bottom row). However, on intrusion trials, the target’s key and response features are generally weaker. The target’s key feature being weaker means a later blaster firing, which is shown as later intrusion vN2pcs in Figure 12A, since the vN2pc is the blaster’s output. Pre- and post-target intrusions occur if respective response TFL/ type neurons are stronger than the target’s response TFL/ type neuron. Importantly, in these cases, the blaster fires temporally further from the activation peak of the subsequently bound distractor type’s response TFL/ type neuron. This response TFL/ type neuron therefore requires more time to overcome the lateral inhibition of this layer and win the competition to the binding pool. The additional time this process requires in intrusion trials leads to delayed vP3s (which reflect activation levels of the response TFL/ type layer & the response binding pool) as well as increased mean RTs. We call this phenomenon a *loss of responsiveness* of these weak types, which is discussed in detail in Appendix 4. Notably, this delay is substantial enough for pre-target intrusion errors to be bound later than correct trials, even though their corresponding inputs were presented earlier in the virtual RSVP sequence, generating the counterintuitive RT pattern.

Essentially, the counterintuitive RT pattern follows from the blaster being fired by target activation in the key pathway. This yields a pattern of blaster firing whereby the blaster tends to fire closer to the optimal activation level in the response pathway for the target than non-target items (for example. the −1 distractor).

As an illustration, consider a −1-intrusion error that, on average, occurs late relative to a Correct response. This would have the following characteristics:

1. by the chance coincidence of random sampling of activation strengths and time constants, on some trials, target strengths are weak in key and response pathways, while the −1 item is stronger in the response pathway;
2. due to its weakness, the key pathway target fires the blaster relatively late;
3. at this time-point, due to its greater strength, the −1 item is the most active in the response pathway, but has entered what we call a low responsiveness regime, i.e. because of the delay before the blaster fires, the −1 item’s activation is not increasing rapidly and may even be reducing already;
4. this low responsiveness means that, despite blaster firing affecting all items the same, relatively speaking, more time is required to bring −1’s activation levels above threshold.
5. these factors ensure that, when a −1 item is encoded, it is encoded late, relative to a target item, and thus exhibits a slower reaction time.

### Experiments with response feature category non-matching distractors

We will now focus on replicating experimental paradigms in which only the distractor immediately following the target (i.e., the +1-intruder item) matched the target’s category along the response dimension. Specifically, we will account for behavioural as well as electrophysiological results from Zivony and Eimer’s (2021) Experiments 1 (combining Experiments 1B and 1A, since 1B was a direct replication of 1A) and 2. The dual-stream RSVP paradigms of the two experiments are illustrated in Figure 13. In Experiment 1, participants were presented with grey stimuli, with letters as distractors and digits as targets. The target digit was differentiated by a surrounding annulus or square. Participants had to report the target after each trial. In target frames, a distractor letter was also presented in the other RSVP stream surrounded by either an annulus or square (which shape identified the target digit was pre-specified). The frame preceding the target frame always consisted of two letters (one in each stream) and earlier pre-target frames were equally likely to contain two letters or one letter and one digit (to ensure that attentional allocation was placed according to the annulus or square rather than alphanumerical category, i.e., subjects did not just search for the first digit in the stream). The frame that followed the target frame included another digit at the same location on 75% of trials. In the remaining 25%, a distractor letter was presented instead (Zivony & Eimer, 2021). Hence, the annulus and the square were the key features in this setting and digit identity the response feature. Experiment 2 adopted a similar dual-stream paradigm. However, participants now had to report the first coloured digit. The following post-target distractor stimulus was either a digit or a letter and either presented in grey or coloured (Figure 13’s bottom panels A-D depict these scenarios). As a result, colour was the key and digit identity the response feature in Experiment 2.

**Figure 13.**
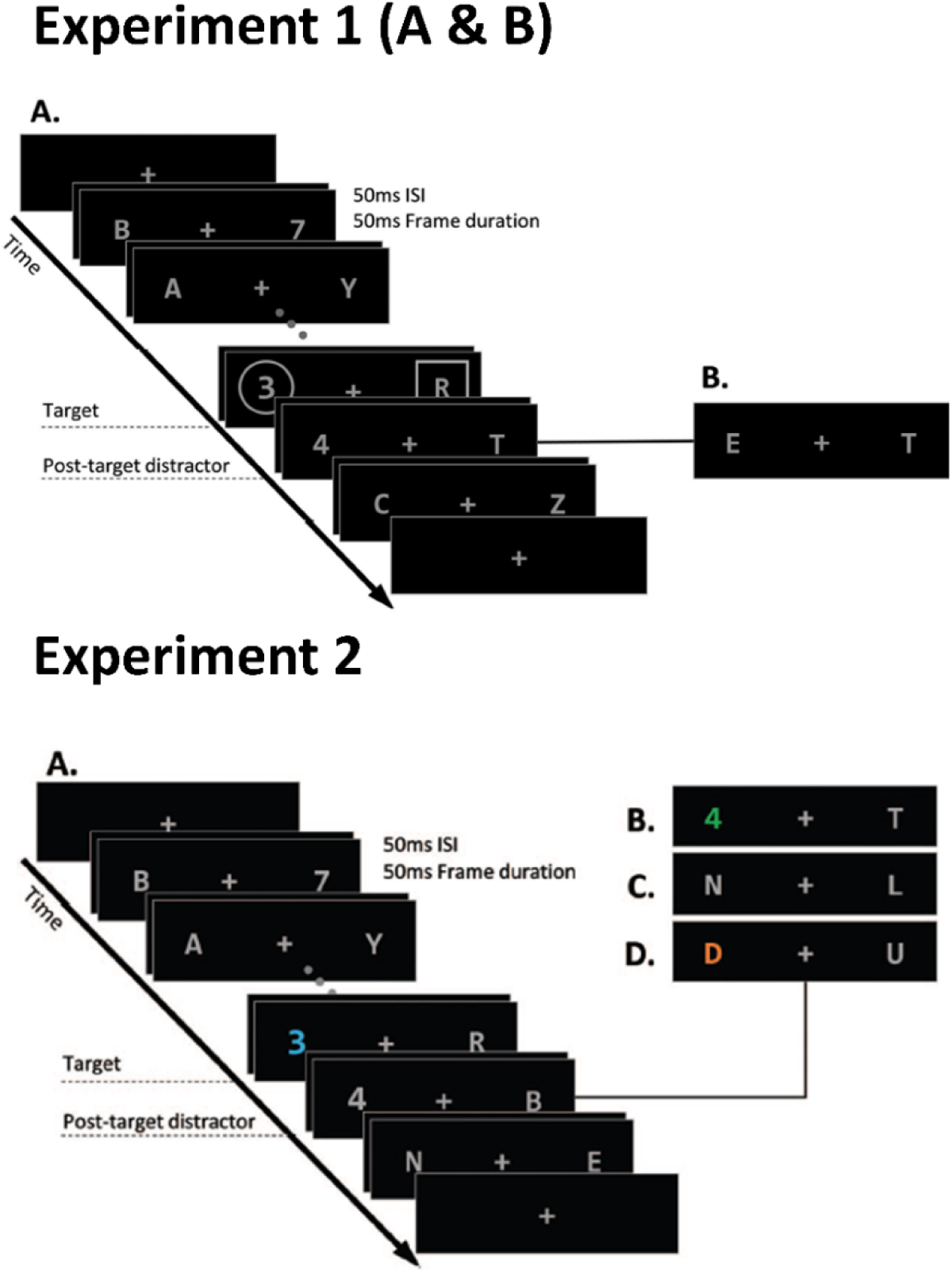
Experimental paradigms of Zivony and Eimer’s (2021) Experiments 1 and 2. Experiment 1 adopted an annulus or square surrounding the digit in target frames, whereas in Experiment 2, target stimuli were coloured. Hence, Experiment 2 implemented a more salient key feature and was simulated with the 2f-ST^2^ model using a lower key delay (τK) value (of 13 time-points) than that used to simulate Experiment 1 (τK = 36).

We computed event-related potentials (ERPs) using the electroencephalography (EEG) data of these two experiments to later reproduce them with the 2f-ST^2^ model’s vERPs. ERPs were computed separately for trials in which participants reported the target-digit correctly (correct trials) and for reports of the post-target digit-distractor stimulus (intrusion trials).

Incorrect trials, i.e., with reports of neither the target nor the post-target digit-distractor, were excluded in ERP analyses. N2pcs, suggested to index attentional processing (Eimer, 1996; Kiss et al., 2008; Luck & Hillyard, 1994), were computed as the contralateral – ipsilateral difference wave between PO7 & PO8 electrodes w.r.t. the location of the target (e.g., PO8 – PO7 if the target was presented in the left RSVP stream). The P3 component, suggested to index WM consolidation (Polich, 2007), was defined as the ERP amplitude at the Pz electrode. Hence, we retained the original paper’s (Zivony & Eimer, 2021) EEG methodology, with the addition of a 25 Hz low-pass filter for P3s and a larger time-window of interest (because the original paper did not analyse P3 components).

Zivony and Eimer’s (2021) experiments, particularly due to their dual-stream RSVP nature, adopted key features that were much harder to detect than those used in the early work of Botella and colleagues (Botella, 1992; Botella et al., 2001). Therefore, τK, the fixed delay to key pathway processing in the 2f-ST^2^ model, had to be set to rather large values to simulate the lower salience of key features. Specifically, we used τKs of 36 and 13 time-steps to simulate Zivony and Eimer’s (2021) Experiments 1 and 2, respectively. The τK value required to simulate Experiment 1 was higher, since a geometrical shape surrounding target digits is more difficult to notice (i.e., less salient / delayed key feature processing) than a colour change (in which, for example, attention does not have to be redirected spatially from the stream’s alphanumeric characters for the key feature to be detected). We furthermore implemented task-filtering in the response TFL/ type layer, which ensured that only two response types (target and +1-intruder) could progress to the binding pool. We also implemented a mechanism that ensures that the impact of our task-demand units (TDUs) on both pathways’ TFL/ type layer neurons, which implements the layers’ task-filter via a positive or negative bias, is only active *after* any delay in key pathway processing. This was done to prevent cases with a large key delay in which TDUs affected TFL/ type-layer neurons *before* the virtual RSVP-stream commenced.

### Replicating the Behavioural Data of Zivony and Eimer (2021)

The 2f-ST^2^ model replicates Zivony and Eimer’s (2021) human response distributions of non-missed trials well (Figure 14). Experiment 1’s (low key feature salience) distribution of 37.74% correct and 62.26% intrusion was replicated with model-responses of 40.9% correct and 59.1% intrusion with a τK of 36 (Figure 14A) time-steps. Experiment 2’s (high key feature salience) distribution of 58.54% correct and 41.46% intrusion was replicated with model-responses of 57.7% correct and 42.3% intrusion with a τK of 13 (Figure 14B) time-steps. The values mentioned for Zivony and Eimer (2021) are averages across Experiments 1A and 1B (being direct replications) for Experiment 1 and across gray and coloured distractors for Experiment 2.

**Figure 14.**
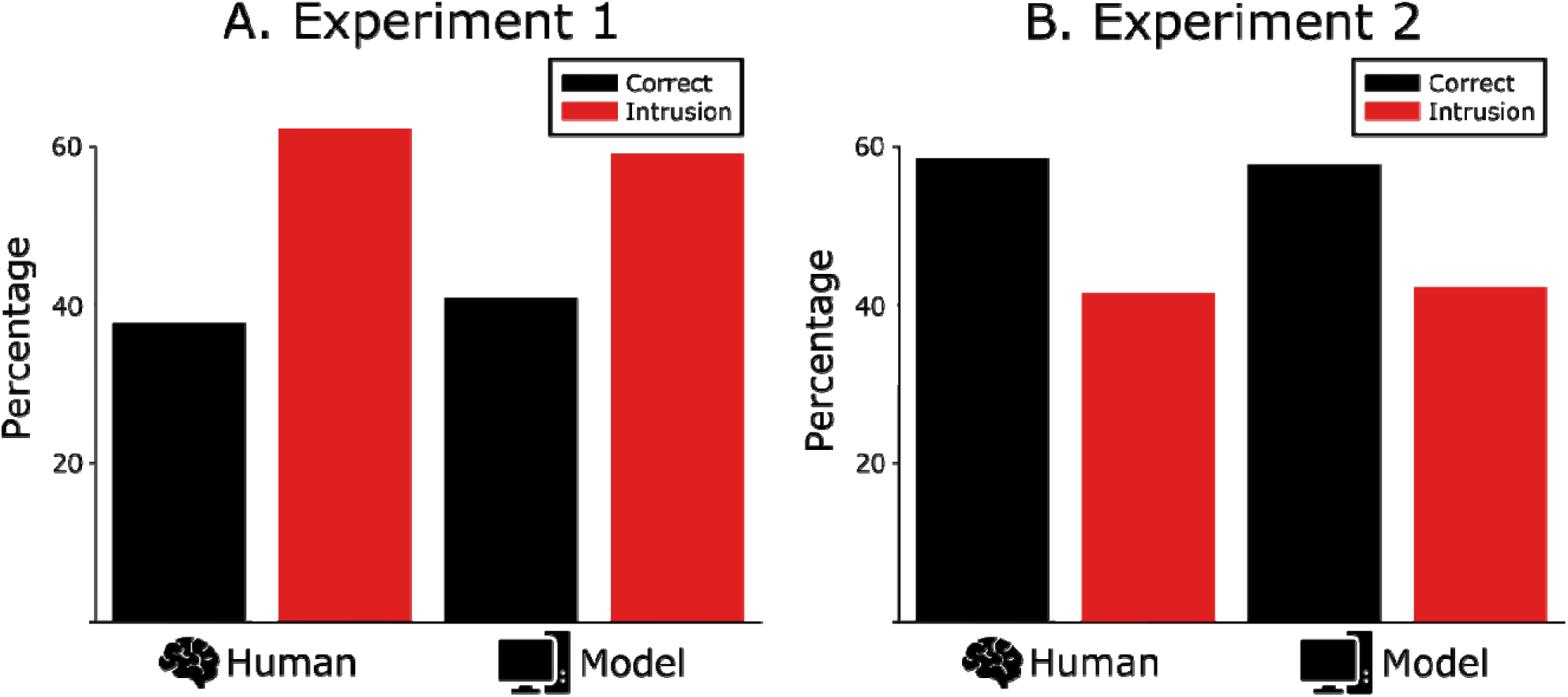
Zivony and Eimer’s (2021) response distributions next to the 2f-ST^2^ model’s. In all panels, black is correct and red intrusion. The 2f-ST^2^ model replicates the accuracy patterns of Experiment 1 with a τK of 36 (Panel A) and those of Experiment 2 with a τK of 13 time-steps (Panel B).

### Replicating the Electrophysiological Data of Zivony and Eimer (2021)

Figure 15 plots the human ERP components of Zivony and Eimer’s (2021) Experiments 1 (Figure 15 A & B) and 2 (Figure 15 C & D), with black and red lines representing correct and intrusion conditions, respectively. Zivony and Eimer (2021) ran two between-experiment ANOVAs on the N2pcs of the two experiments, which revealed that the N2pc emerged earlier in Experiment 2 (187 ms) than 1 (225.5 ms) and that N2pc amplitudes were larger in Experiment 2. The amplitude ANOVA furthermore yielded a significant interaction between experiment and response condition, implying that the enhancement of N2pcs for correct compared to intrusion trials was more substantial in Experiment 1 than in Experiment 2. The latter is easily observable in Figure 15, as the N2pc of correct (black) trials shows a much larger amplitude than intrusion (red) trials’ N2pc in Experiment 1 (Figure 15A). However, in Experiment 2 (Figure 15C) this difference in amplitudes is much weaker. Also, see Hosseini et al. (2024) for further empirical analyses on the N2pc and P3 components of Zivony and Eimer (2021).

**Figure 15.**
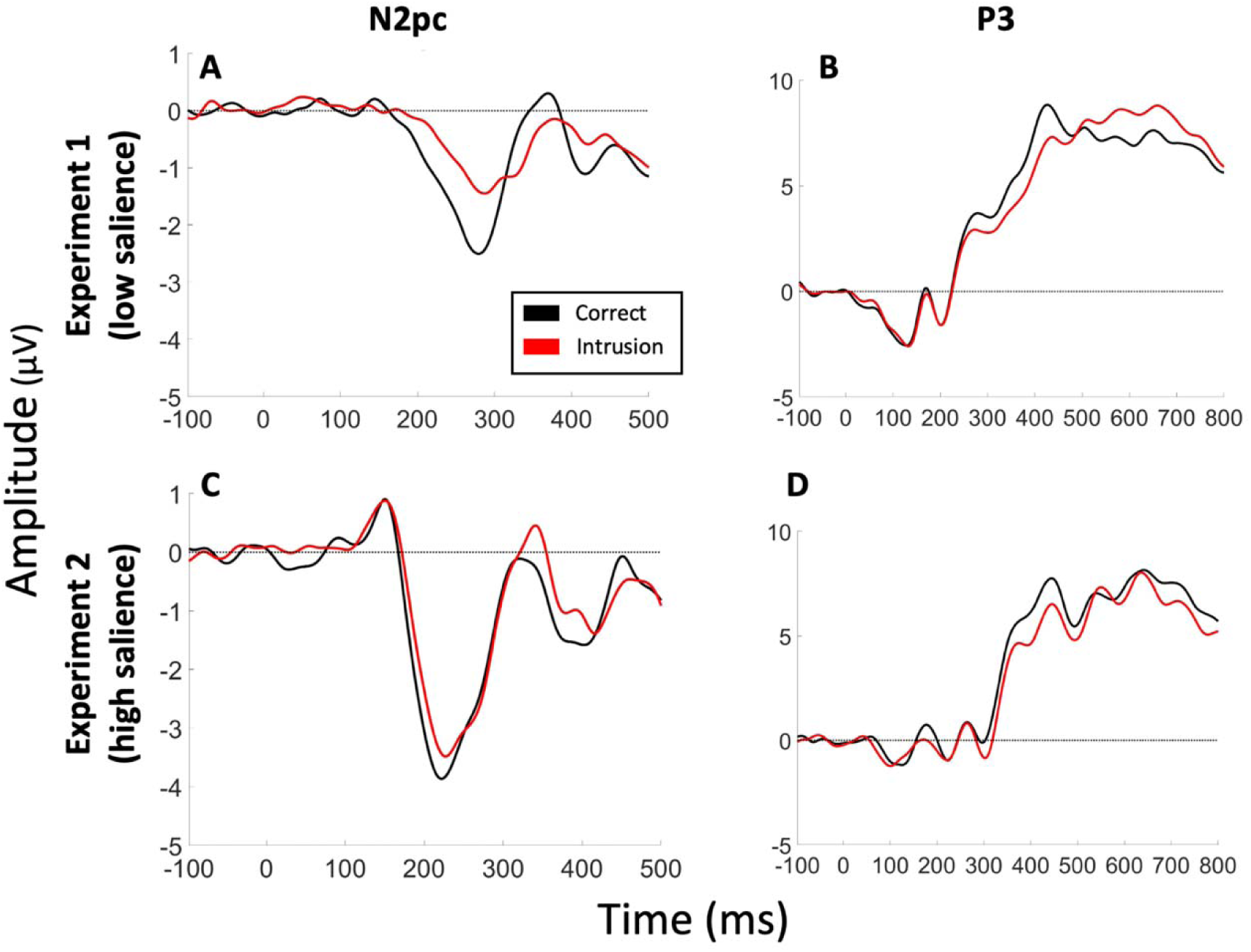
Human ERP Comparison plots. Panels A and C present the N2pc components of Experiments 1 and 2 of Zivony and Eimer (2021), respectively. Panels B and D present the P3 components of Experiments 1 and 2, respectively. Black and red lines correspond to correct and intrusion trials.

We present comparison plots of the 2f-ST^2^ model’s vERPs, with τK values of 36 and 13 time-steps, replicating Zivony and Eimer’s (2021) Experiments 1 and 2, respectively, in Figure 16 and Figure 17. These vERP plots follow the plotting conventions of Figure 12, with the only difference being that black and red vERP traces represent correct and (+1) intrusion trials, respectively, since no other response conditions were present in this case. Figure 16 illustrates that we qualitatively replicate the N2pc patterns found by Zivony and Eimer (2021) in humans: earlier and larger (i.e., higher amplitude) (v)N2pcs in Experiment 2/τK of 13 (Figure 16 A & C) than 1/τK of 36 (Figure 16 B & D). The 2f-ST^2^ model also shows smaller amplitude-differences between correct and intrusion trials with a τK value of 13 than with a τK of 36 time-steps. We therefore also qualitatively replicate the significant interaction effect between experiments and response conditions demonstrated in Zivony and Eimer’s (2021) N2pc-amplitude ANOVA.

**Figure 16.**
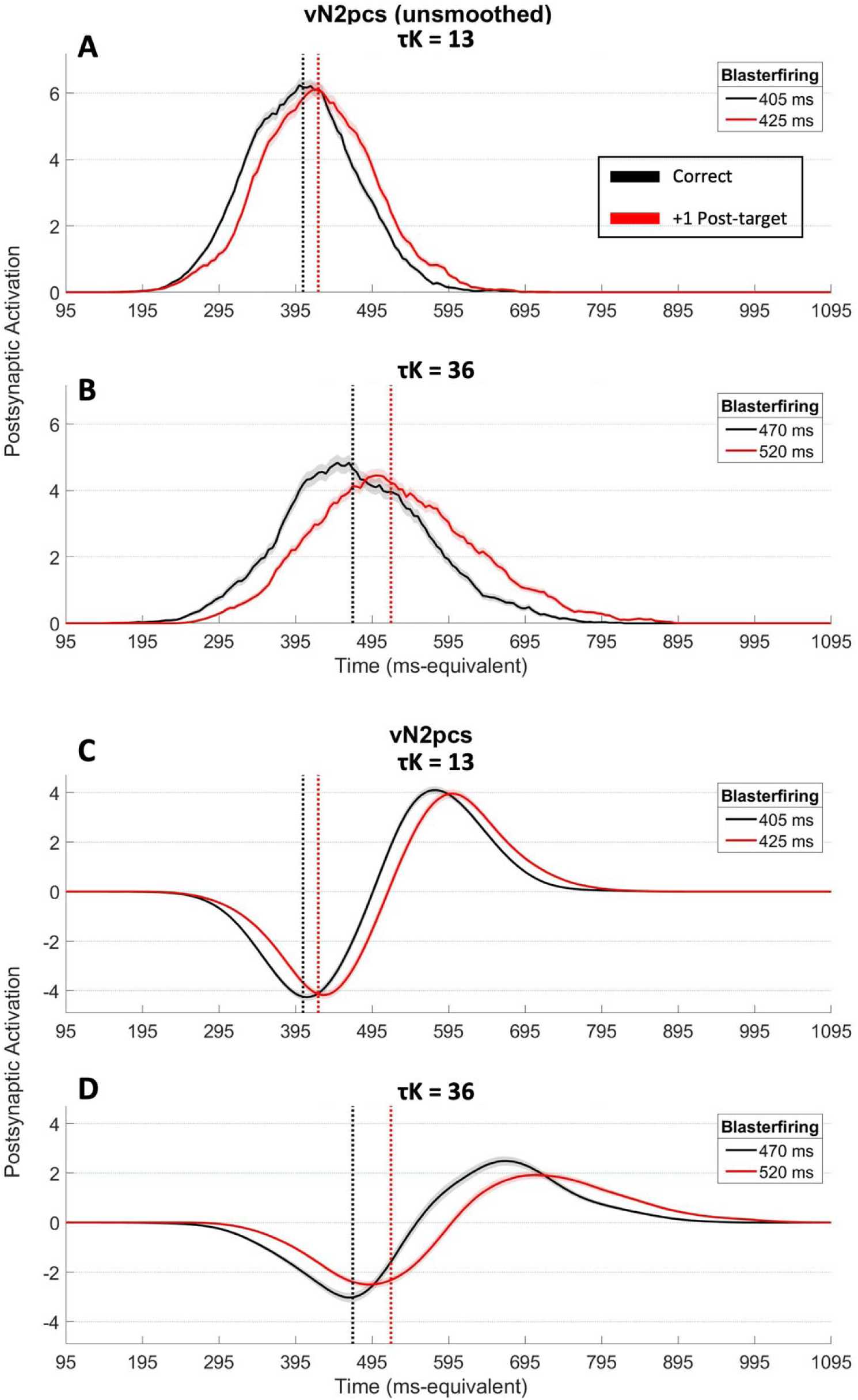
vN2pc Comparison Plots, with model run with active task-filtering in the response pathway. Panels A and C plot the unsmoothed and smoothed vN2pcs of the 2f-ST^2^ model with τK = 13 time-steps. Panels B and D show these with τK = 36. In all panels, black and red lines correspond to correct and +1 intrusion trials, respectively, and vertical dotted lines indicate average blaster-firing latencies, with respective numerical values in plots’ legends. The 2f-ST^2^ model qualitatively replicates the human N2pc results shown in Zivony and Eimer’s (2021) Experiments 1 and 2.

**Figure 17.**
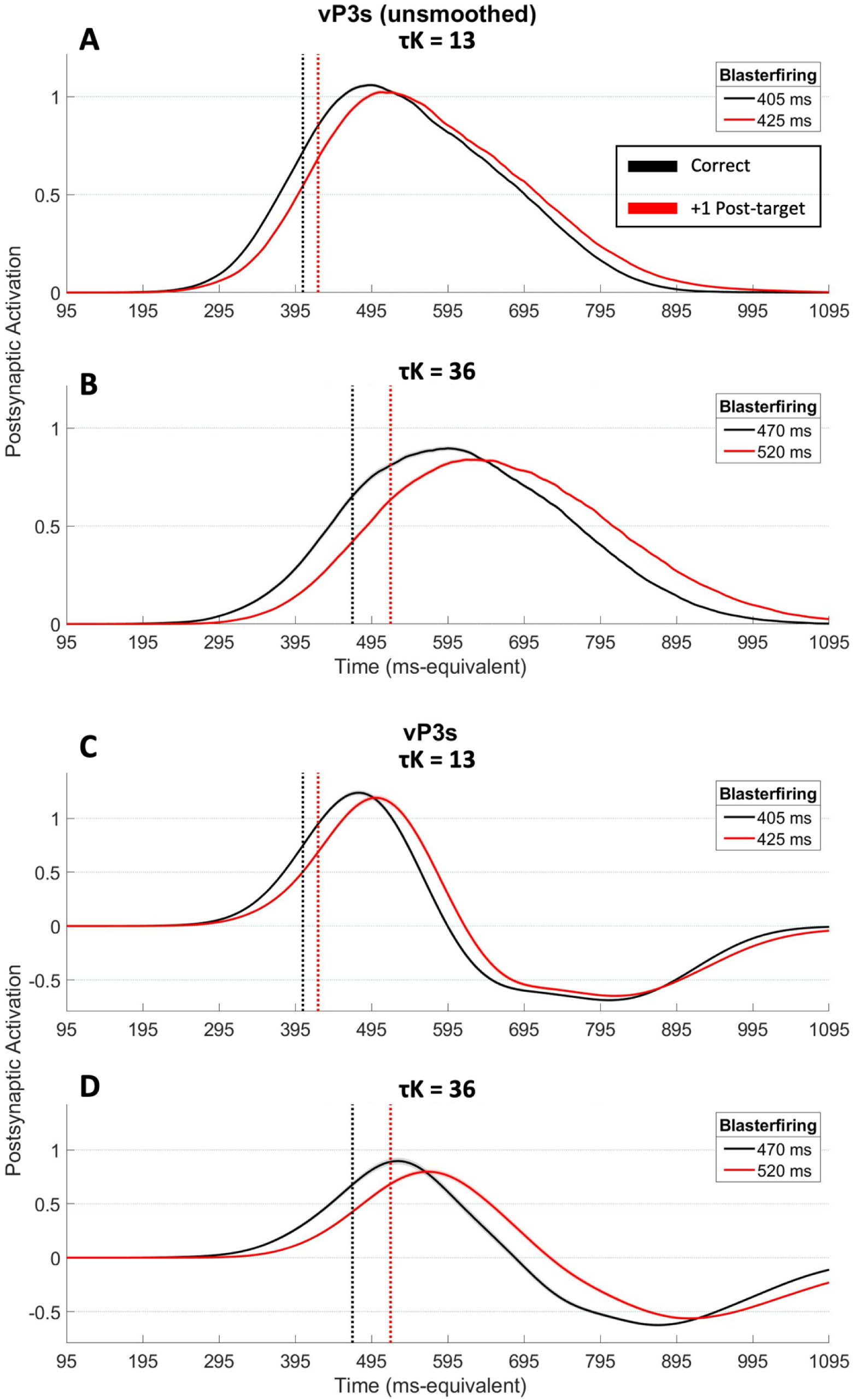
vP3 Comparison Plots. Plotting conventions are identical to those of Figure 16.

Figure 17 displays the vP3s after running the 2f-ST^2^ model with τKs of 13 (Figure 17 A & C) and 36 (Figure 17 B & D) time-steps. Again, the vP3s of both response conditions were earlier and larger amplitude with a τK of 13 time-steps. This is in line with the patterns shown in Figure 16, which is to be expected because of the direct impact the blaster (vN2pc) has on later model-layers captured in the vP3 (see Hosseini et al. (2024) for evidence of a temporal correlation between the N2pc and P3 components in humans, supporting this facet of the 2f-ST^2^’s model architecture).

### Modelling the temporal variability underlying transient attentional enhancement (TAE) with Gamma Noise

The 2f-ST^2^ model’s *loss of responsiveness* phenomenon, introduced earlier and presented in detail in Appendix 4, prevented the accurate replication of Zivony & Eimer (2021) electrophysiological data with the model’s vERPs at first (despite facilitating the replication of the important and counterintuitive RT data provided by Botella (1992)). To overcome this limitation as well as to increase the neurophysiological plausibility of our computational model, we added a new source of temporal variability to the model. In particular, gamma distributed variability (henceforth called Gamma Noise) was added to each trial’s onset of key pathway processing, which in turn added temporal variability to blaster firing latencies. The extent of this variability was furthermore modulated by τK. For a complete run of the model, τK was fixed, simulating different processing speeds of different kinds of key features determined by their salience. This (globally fixed) τK value determined the extent to which trial-level τKs (denoted τK^tr^) varied due to Gamma Noise, adding more Gamma Noise to the model’s key pathway as τK increases (Appendix 5).

The concept of temporal variability of the TAE underlying intrusion errors is central to theoretical accounts, such as those of Zivony and Eimer (2021), Vul et al. (2009) (see their selection window, which is initialised by detecting the target key feature) or Botella and colleagues (Botella et al., 2001) (albeit only for the sophisticated guessing process). The time-point of the blaster impacting later Stage 1 layers of the 2f-ST^2^ model is likewise the main mechanism leading to intrusion errors.

Importantly, the initial 2f-ST^2^ model (Chennu et al., 2011) only manipulated the blaster’s temporal variability via τD, the random delay added to each item’s presentation length.

Critically, even though this delay was different for different items, it was held constant *across the two pathways* for a given item and trial. This approach kept the model’s two pathways synchronised (which carries a set of important implications; see Appendix 4). Implementation of the blaster’s onset variability via τD assumes a very early (in terms of the brain’s processing pathway) source of temporal variability that affects key and response feature processing identically, thereby proposing that these two kinds of features do not have separate sources of temporal variability. However, it would be surprising if such a limited incorporation of temporal jitter would offer a completely satisfactory model of intrusion errors. The addition of Gamma Noise, presented in detail in Appendix 5, increased the neurophysiological plausibility of the 2f-ST^2^ model, whilst preserving its capacity to simulate the main distractor intrusion findings (also see related considerations in the context of the 2f-ST^2^ model’s loss of responsiveness in Appendix 4 and Hosseini (2022)).

### Responding more than once – Replicating Vul et al.’s (2009) Experiment

Vul et al. (2009) ran a distractor intrusion RSVP experiment in which subjects had to guess the identity of the target stimulus four times: best guess, second best, and so on. The authors formulated a theoretical (contiguous-graded selection) account, proposing that once the target key feature is detected, a time-window of selection is placed around the target stimulus. Item (i.e., target as well as proximal distractor) representations presented in this time-window are temporarily stored in short-term memory with one item being subsequently selected for further processing, allowing its identification and report. Selecting one item for further processing out of the candidate items involves sampling from the stored probability distribution. Critically, the authors demonstrated that the distribution of Guess 2 reports was not modulated by which item was provided as Guess 1. This motivated the conclusion that the temporal position of the selection window does not vary considerably across trials and that both guesses are sampled from the *same* probability distribution (Vul et al., 2009). Central to the authors’ explanation of this finding is that a probability distribution across potential responses is represented on each trial, with the probability associated with a response item determined by its temporal distance from the point of key feature selection. It is this probability distribution that they argue is sampled from, giving their contiguous-graded selection account (Vul et al., 2009). In this section, we replicate the findings of Vul et al. (2009) with the 2f-ST^2^ model.

In some trials of the 2f-ST^2^ model, more than one *response* type is bound to the currently active token. This is in line with earlier ST^2^ models (Bowman & Wyble, 2007), in which multiple types being bound to a token allowed lag-1 sparing of the attentional blink to be simulated. We will call these multi-bindings. Importantly, the 2f-ST^2^ model’s multi-bindings are binary rather than graded; that is, items are either encoded into tokens or not, but there is no sense to which these encodings have a strength, beyond zero or one. In probability distribution terms, 2f-ST^2^ only represents *uniform* distributions across the responses with non-zero probability (i.e. that are allocated to a token). So, this approach is more like the Contiguous All-or-None Selection that Vul et al. (2009) highlight. In particular, the non-uniform distribution that is observed only emerges when averaging across many trials, each exhibiting one of a variety of these distributions, with varying support. In the 2f-ST^2^ model, multi-bindings are resolved by randomly selecting one of the response types bound to the token for report.

The occurrence of multi bindings in our model depends on the strength of the lateral inhibition present in the response TFL/ type layer. This is because as lateral inhibition gets progressively weaker, it is more likely that more than one node in this layer has sufficient activation to excite their corresponding response binding pool unit, binding its type to the currently active token. Figure 18 demonstrates the interaction between lateral inhibition strength and likelihood of multi-bindings, plotting the proportions of misses (no type bound to the token), single bindings (only one type bound to the token) and multi bindings as a function of lateral inhibition in the range of -.05 (value used throughout this paper) to 0 (no lateral inhibition).

**Figure 18.**
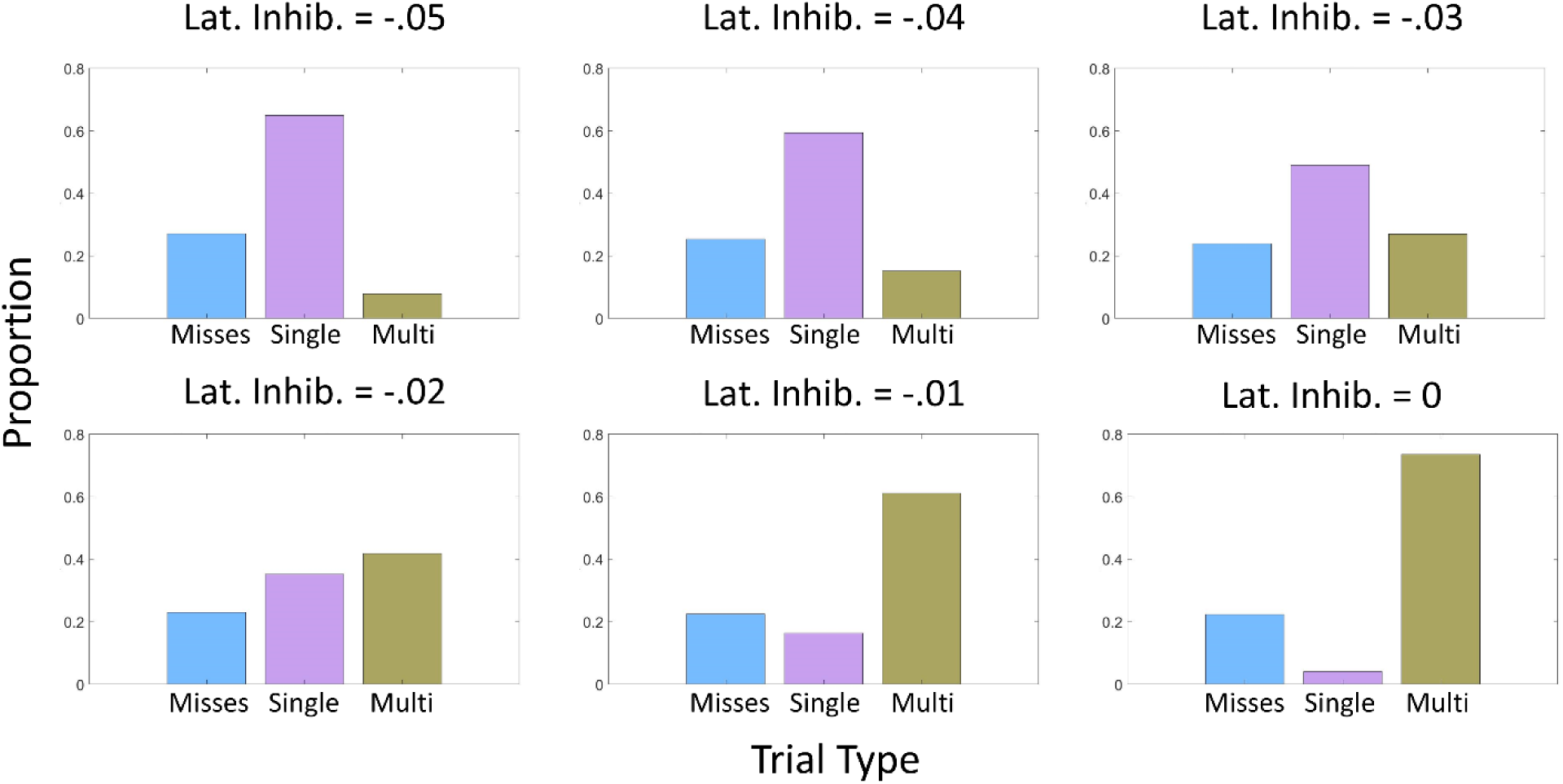
Proportions of misses, single bindings and multi bindings made by the 2f-ST^2^ model as a function of lateral inhibition. As lateral inhibition weakens (going from top left to bottom right), misses and single bindings decrease and multi bindings increase. Weakened lateral inhibition can be regarded as the equivalent of different top-down strategies adopted by the brain to adapt to different task demands.

These values were obtained after a single run of the model, without task-filtering in the response pathway and without any processing delays (i.e., τK & τR = 0).

For example, if multiple guesses after a trial are required by subjects, the threshold to WM encoding should be lower (weak or no lateral inhibition) than if subjects are asked to only report one response (strong lateral inhibition).

Different values of lateral inhibition in the response TFL/ type layer simulate different top-down strategies adopted based on task demands. This notion has been introduced as a flexible encoding threshold by Zivony and Eimer (2020) (see the authors’ Figure 9 and Experiment 4 in particular). In essence, if only one item has to be reported after a trial, the encoding threshold is set to a high (conservative) value, only allowing the strongest (i.e. maximum likelihood) item into WM. However, if more than one response is required as in many attentional blink studies and also that of Vul et al. (2009), the threshold will be set to a low value.

Since in Vul et al.’s (2009) experiment, subjects knew that multiple guesses will be provided, we removed lateral inhibition from the response TFL/ type layer to simulate a low encoding threshold. The following are the only results generated by the 2f-ST^2^ model without lateral inhibition, since in all other empirical paradigms replicated only one item was reported after a trial.

To simulate Vul et al.’s (2009) experiment, we introduced guessing to the 2f-ST^2^ model, resolving multi-bindings as done originally (i.e., randomly select and report one of the multiple types bound to the token) repeatedly until all types bound to a specific token were guessed, establishing a guess sequence. For example, if four types were bound to a token, Guess 1 was selected out of these four types, Guess 2 was selected out of the remaining three, and so on.

Selection of types was always uniformly random. The model was run 20 times, providing 27440 trials. Figure 19 presents the distribution of responses provided as Guess 1 (which is identical to what we call the response distribution in other configurations of the 2f-ST^2^ model). Figure 19 only includes the response range from −5 to +5 and demonstrates more pre-target than post-target intrusions, which is in line with the empirical data of Vul et al. (2009).

**Figure 19.**
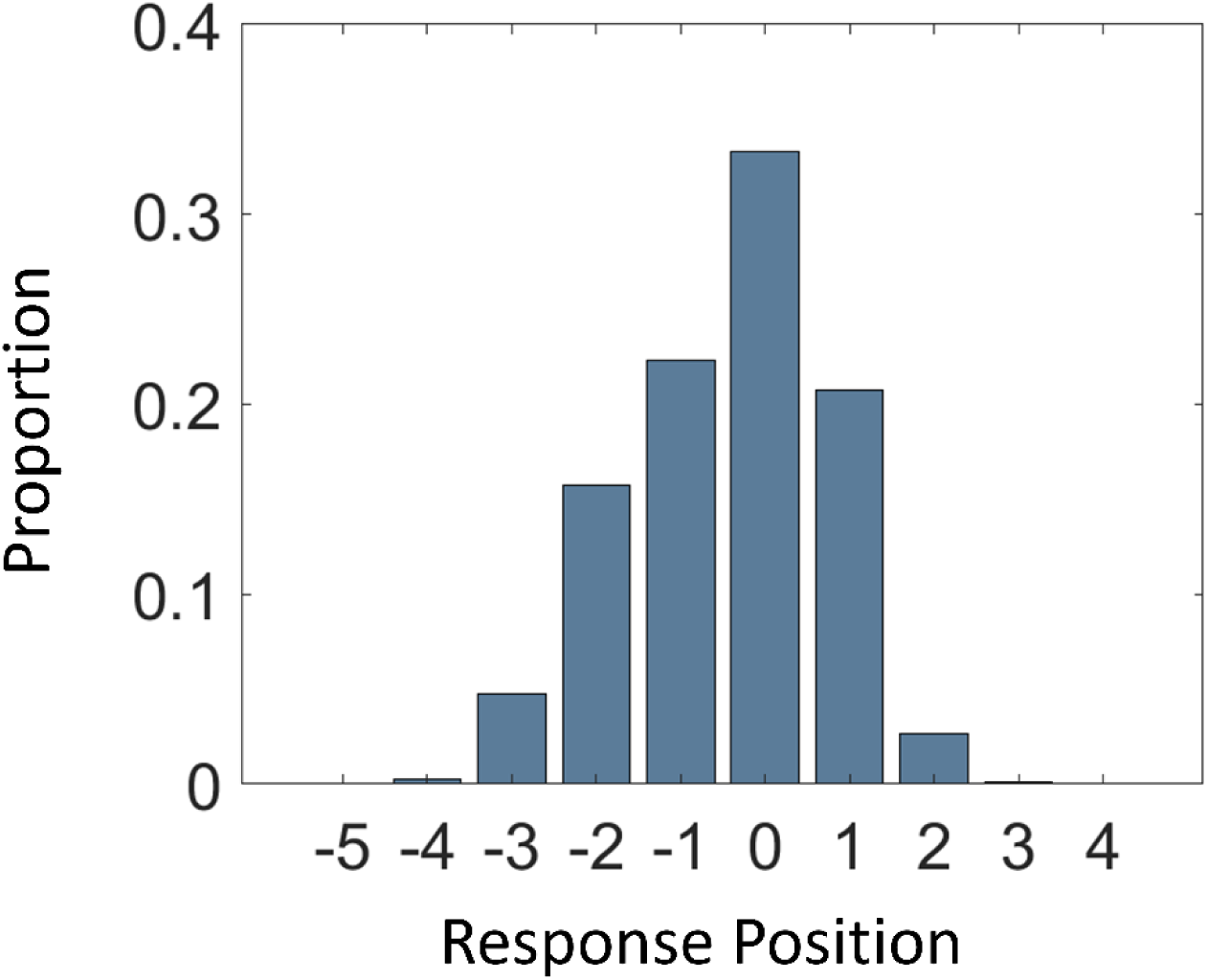
Guess 1 (i.e., response distribution) of the 2f-ST^2^ model without lateral inhibition in the response TFL/ type layer, task-filtering in the response pathway or any temporal delays to either pathway’s processing onset. Response position of 0 indicates a correct trial. 20 identical runs of the 2f-ST^2^ model, employing different rng-seeds, were combined (total number of trials 27440).

A main finding of Vul et al. (2009) was that the distributions of responses provided as Guess 2 were constant whatever the serial positions of Guess 1, and that Guess 2 distributions approximated the Guess 1 distribution. The authors stated that this pattern implied that the *same* probability distribution was used for sampling an item out of the candidate ones held in short-term memory and that the selection window did not exhibit substantial temporal across-trial variability. Figure 20A plots Vul et al.’s (2009) conditional Guess 2 frequencies (as coloured lines) as well as Guess 1 frequencies (as a thick grey line) as a function of serial position. Figure 20B displays this figure for the 2f-ST^2^ model. Differently coloured lines plot different conditional probabilities, with each line corresponding to different serial positions being provided as the first guess (ranging from −2 to +2, 0 = target).

**Figure 20.**
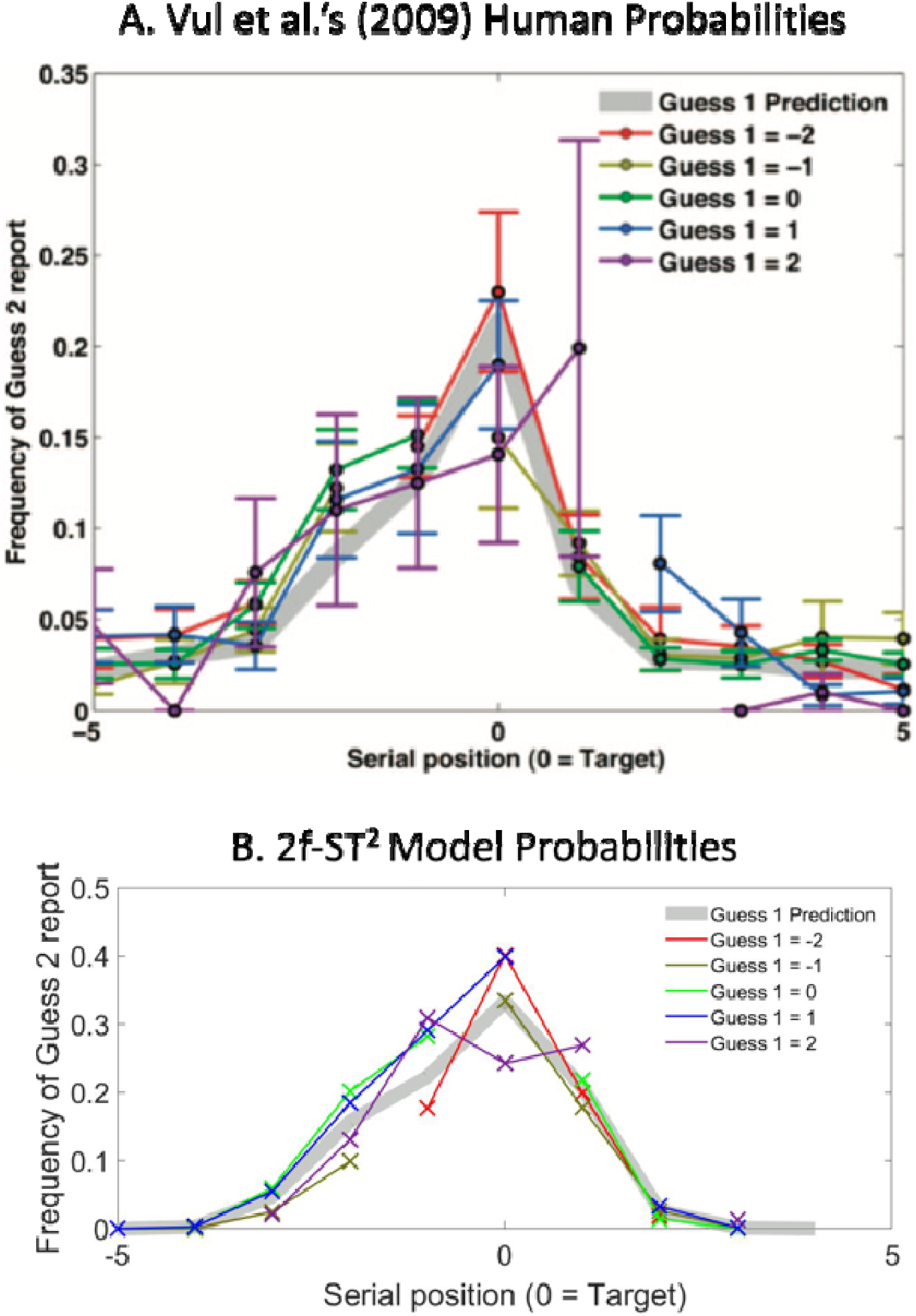
Panel A shows Vul et al.’s (2009) conditional Guess 2 frequencies (as coloured lines) as well as Guess 1 frequencies (as a thick grey line) as a function of serial position. Panel B plots the analogous plot for the 2f-ST^2^ model. We show the results of 20 runs of the model without lateral inhibition in the response TFL/ type layer, task-filtering in the response pathway or any temporal delays to either pathway’s processing onset. Nearest neighbour interpolation was used to estimate missing values so that conditional probabilities are comparable across Guess 1 responses. The 2f-ST^2^ model exhibits the pattern shown by Vul et al. (2009), i.e., that Guess 2 probabilities are independent of Guess 1 serial position and approximate the distribution of Guess 1 responses.

2f-ST^2^ natively generates histograms that have very substantial amplitude differences, due to the different probabilities of different serial positions for Guess 1; for example, −2 Guess 1’s are very rare, causing the histogram for *Guess 2, given Guess 1 = −2*, to have few data points. Consequently, we normalised these histograms to generate probability distributions. However, all these *Guess 2, given Guess 1* histograms have a gap (i.e. missing value) at the serial-position of Guess 1 and furthermore, the amplitude relative to other points in the histogram of this missing value varies with Guess 1’s serial position. For example, the gap at −1 for the olive-coloured line (Guess 1 = −1) will be of lower amplitude than that at 0 for the bright green-coloured line (Guess 1 = 0). Accordingly, directly normalising these histograms, while still containing a gap, will not produce meaningful results. To overcome this difficulty, we first approximated the missing point by interpolating it from the part of the histogram that was present using nearest neighbour interpolation with MATLAB’s interp1 function. We then normalised the gap-free histogram to obtain a probability distribution (i.e. with all elements adding to 1) and then removed the point at the Guess 1 serial position^4^. The result of this process is shown in Figure 20B. Figure 20B replicates Figure 20A, demonstrating the key findings that Guess 2 probabilities 1) are independent of Guess 1 and 2) follow the distribution of Guess 1.

The 2f-ST^2^ model replicates this pattern by resolving multi-bindings randomly, which resembles sampling from a uniform distribution of response type representations instead of the graded distribution proposed by Vul et al. (2009).

## Discussion

Solving the binding problem (Feldman, 2013; Treisman, 1998), i.e., binding stimulus-features together accurately, allows us to perceive the objects in our environment in an integrated manner. Conjunction errors, or illusory conjunctions, occur when features are bound together erroneously, leading to the perception of objects that were not in fact presented. This phenomenon has been investigated in RSVP experiments that focused on distractor intrusion errors, which are conjunction errors made in the temporal (as opposed to the spatial) domain.

Previous attempts to account for distractor intrusions in RSVP (Botella et al., 2001; Chennu et al., 2011; Vul et al., 2009; Zivony & Eimer, 2020, 2021) provided important contributions, but a complete explanation of these errors considering all empirical paradigms and their corresponding findings was lacking. In this paper we provide such a complete explanation with the 2f-ST^2^ neural model, which accounts for the most important empirical findings obtained in distractor intrusion work thus far. They include empirical response distributions (Botella et al., 2001; Zivony & Eimer, 2021), counterintuitive reaction time data (Botella, 1992), results from experimental paradigms involving multiple responses (Vul et al., 2009), and electrophysiological findings (Zivony & Eimer, 2021). These findings were obtained in experiments adopting two different paradigms, with either all distractors (Botella, 1992; Botella et al., 2001; Vul et al., 2009) or only the +1 distractor (Zivony & Eimer, 2021) sharing the response feature dimension of the target. The 2f-ST^2^ model accounted for both paradigms via the same basic architecture.

Moreover, we would argue that the 2f-ST^2^ model architecture is more parsimonious than the main competitor model, Botella et al.’s (2001) focussed-attention/ sophisticated-guessing model, by not requiring separate focussed attention and sophisticated guessing pathways. Moreover, the 2f-ST^2^ model benefits from fitting within a broad scope cognitive theory.

Much of computational modelling in cognitive neuroscience is piecemeal, with few over-arching, broadly applicable, models. That is, each different experimental phenomenon is often considered through the lens of a different model. While certainly useful, such modelling does not reflect the general-purpose nature of the brain, i.e., it is the *same* cognitive system that exhibits all of the phenomena. The attempt here to integrate our modelling of distractor intrusion errors into an existing “broad scope” model moves away from piecemeal theory development towards more unified modelling. As previously discussed, a large spectrum of temporal attention and working memory phenomena have been modelled within the ST^2^ framework (Bowman et al., 2022; Bowman & Wyble, 2007; Chennu et al., 2009; Craston et al., 2008; Jones et al., 2020; Wyble et al., 2009; Wyble et al., 2011), suggesting a single common mechanism across a range of temporal attention phenomena. ST^2^ particularly emphasizes the importance of episodic binding enforcing seriality and unity of experience, following a parallel Stage 1 (Avilés et al., 2020; Bowman & Avilés, 2021, 2022; Treisman & Gelade, 1980).

### The critical role of Transient Attentional Enhancement (TAE)

There is plenty of evidence for the temporal spotlight of attention being a central mechanism for perception *in general* (Bowman & Wyble, 2007; Olivers & Meeter, 2008; Reeves & Sperling, 1986; Wyble et al., 2009; Zivony & Eimer, 2020, 2021). We support this notion further, proposing the variability of TAE as the main reason for distractor intrusion errors: on average, early TAE deployment leads to pre-target errors and late TAE deployment to post-target errors. Modelling the *nature* of TAE-variability accurately has been highlighted as a critical issue for obtaining the effects we observe. In this context, we have shown a set of interesting effects that modelling TAE-variability either via τD, as in the early work of Chennu et al. (2011)^5^, or via the new Gamma Noise approach, which affects τK directly, had on the 2f-ST^2^ model’s processes and results.

Whether TAE should be modelled as an additive or multiplicative process remains an important question. In the model presented here it is additive, but there are important multiplicative implementations, for example in the episodic STST model (Wyble et al., 2009). Multiplicative enhancement has a sharpening effect, i.e. differentially benefitting those units that are already strongly active, while additive enhancement produces a similar amplitude “across the board” increase, even to units that were not active at all. However, the benefit of the additive enhancement is that its realisation in neural circuity is straightforward, consistent with a fast-acting attentional enhancement.

### Competition between response feature representations

We further propose that the competition between response feature representations (at the end of the response pathway in the 2f-ST^2^ model) plays a critical role for the percept experienced. As ambiguity increases (i.e., the competitive period is longer and winners win less decisively), the system’s precision decreases and percepts are perceived less vividly, with less confidence. The impact of stimulus-competitions on subjective visibility and confidence scores has been well documented across the field. Visibility effects have been demonstrated in backward masking (Bae et al., 2025; Binder et al., 2017; Jimenez et al., 2019), meta-contrast masking (Lau & Passingham, 2006), crowding (Levi, 2008; Pelli & Tillman, 2008), and attentional blink (Pincham et al., 2016) studies. Confidence effects were demonstrated in backward masking studies (Balsdon et al., 2024; Jaeger et al., 2020; Rausch et al., 2018). In the context of distractor intrusions, in which stimulus-competition occurs due to backward masking, too, the 2f-ST^2^ model implies that the strength of lateral inhibition of response feature representations plays a big role. It is suggested that top-down task-dependent influences likely regulate its strength (being low if subjects guess multiple times, as in Vul et al. (2009), and high if reporting just once, for example). Future work should consider the competition between response features and top-down influences on the nature of this competition when designing and explaining related experiments.

### Relationship to the global workspace framework

The critical contribution of this paper is to provide a complete, unifying, neural implementation of the mechanisms leading to intrusion errors and explicitly show that it can reproduce the most important experimental phenomena in the literature. This includes both behavioural and electrophysiological findings, as well as both intuitive and counter-intuitive findings, particularly, the counter-intuitive reaction time finding of Botella (1992). Furthermore, the 2f-ST^2^ model makes a particular claim about the point at which key and response pathways come together. If one imposes the natural interpretation on the model’s second stage (see Figure 6A), it suggests that the following concepts coincide: occurrence of the (breakthrough-)P3 (see Hosseini et al., 2024 for a discussion on the breakthrough P3, also see Alsufyani et al, 2019, 2021; Bowman et al, 2014), encoding into working memory, conscious perception of a feature conjunction (e.g. of the colour red with the letter K) and the generation of a global brain-scale state^6^. This “global” state would involve the TFL/type layer, the binding pool and tokens, and might be related to the global workspace (Dehaene et al., 1998), but where the functional role of the workspace is to bind types to token through a shared binding resource (the binding pool). This contrasts with Dehaene et al.’s (1998)’s definition of the global workspace, where it mediates competition between brain representations; see Figure 21.

**Figure 21:**
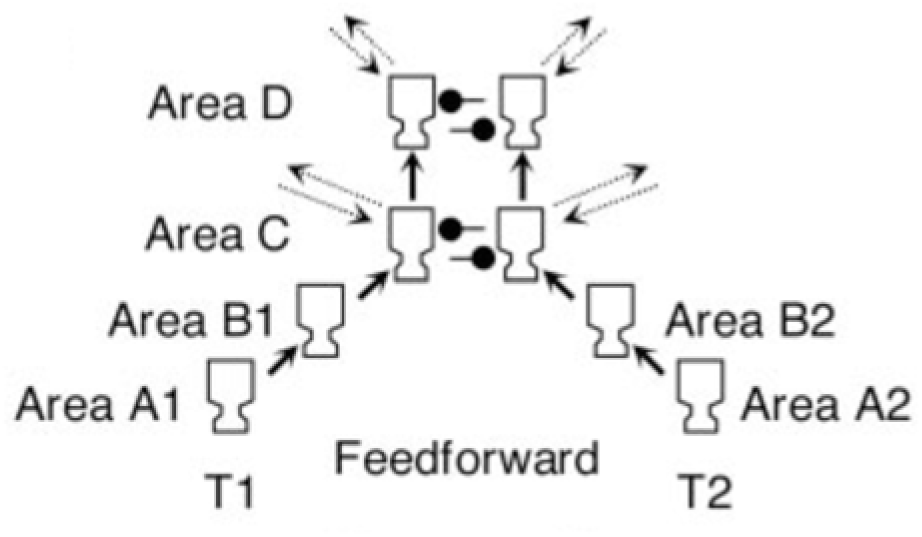
Panel from Figure 1B of Dehaene et al. (2003) showing the competitive nature of the global workspace: see lateral inhibition in Areas C and D.

**Figure 22.**
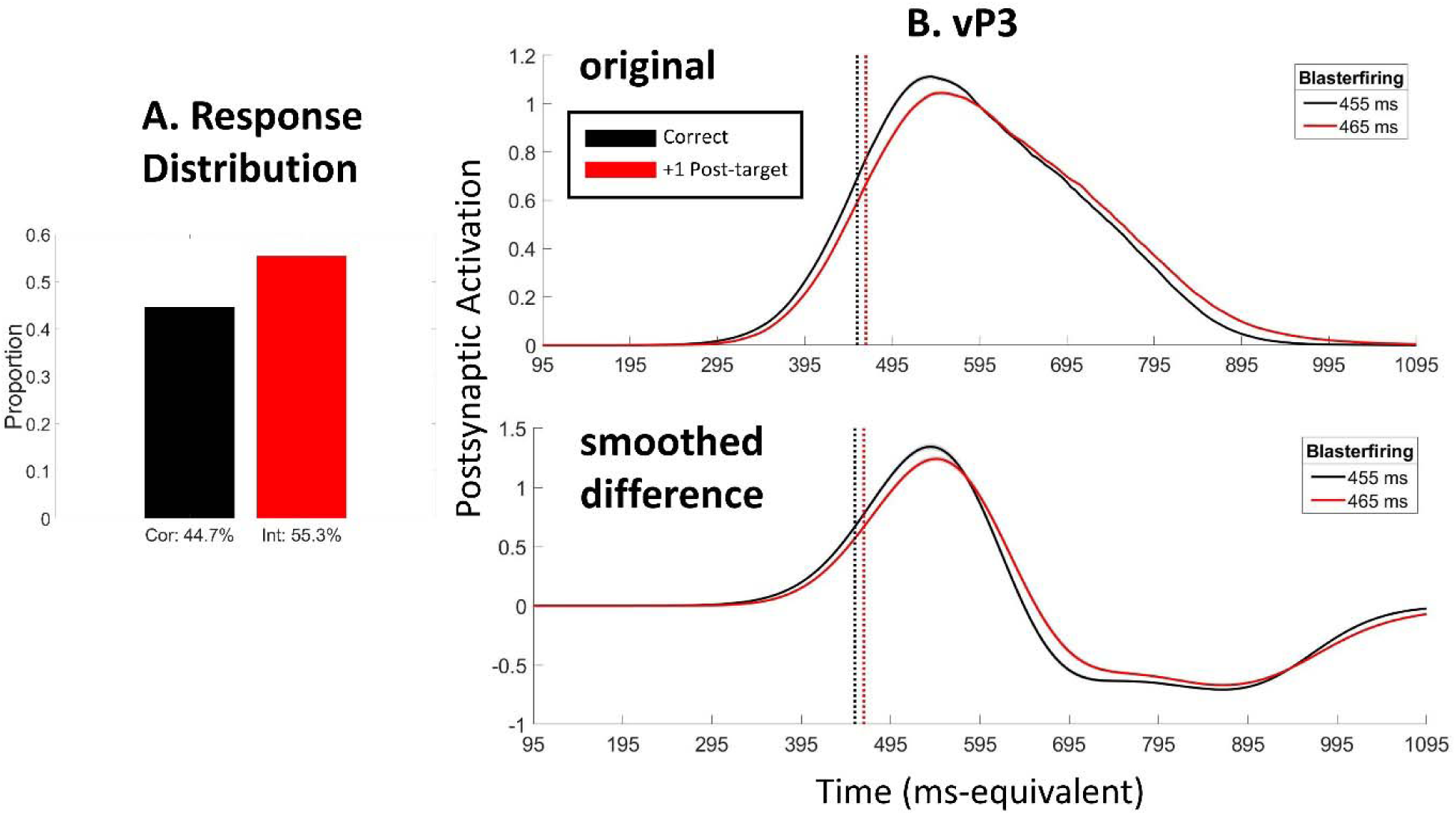
Results of running the intermediate 2f-ST^2^ model with task-filtering in the response pathway and a τK of 24 time-steps, to model Zivony and Eimer’s (2021) Experiment 1. We present the 2f-ST^2^ model’s response distribution and vP3s in Panels A & B, respectively.

**Figure 23.**
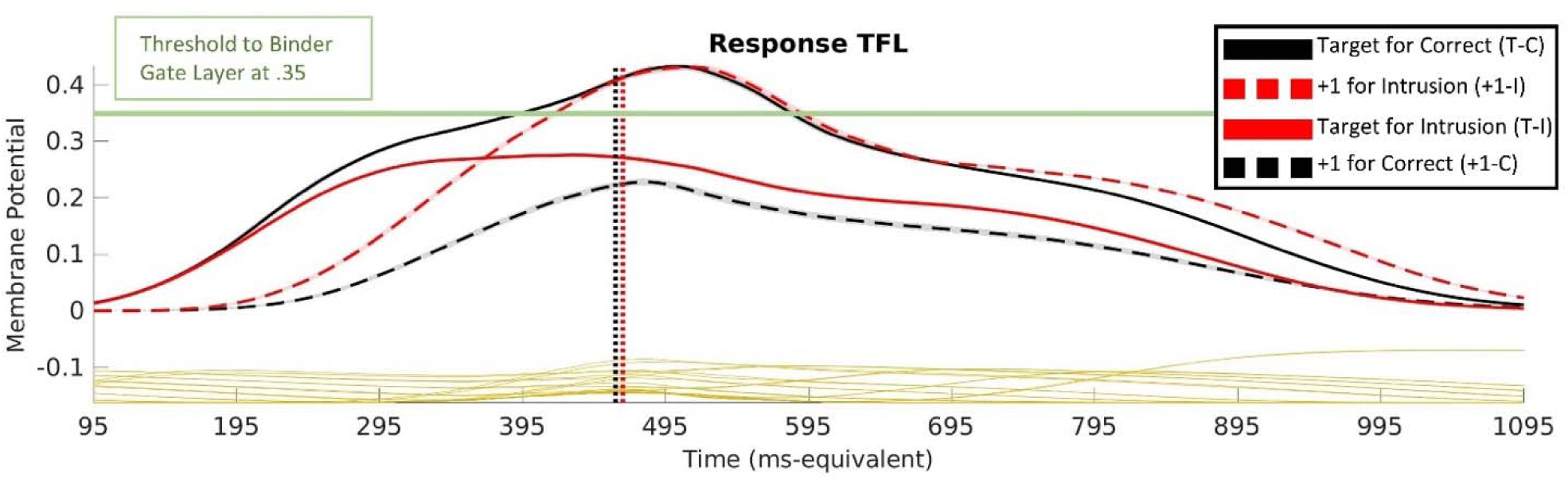
Response TFL/ type layer’s neuronal membrane potentials (VMs). Black = correct-& Red = intrusion-trials. Solid = target unit & dashed = +1 unit. Therefore, black solid, red dashed, red solid, and black dashed lines represent VMs of target unit in correct trials (T-C), +1 unit in intrusion trials (+1-I), target unit in intrusion trials (T-I), and +1 unit in correct trials (+1-C); see legend. The green horizontal line indicates the VM threshold to respective binding pool units at .35. Vertical dotted lines indicate average blaster firing latencies of correct (black) and intrusion (red) trials. Gold traces indicate category-nonmatching distractor types, which do not meet task demands.

**Figure 24.**
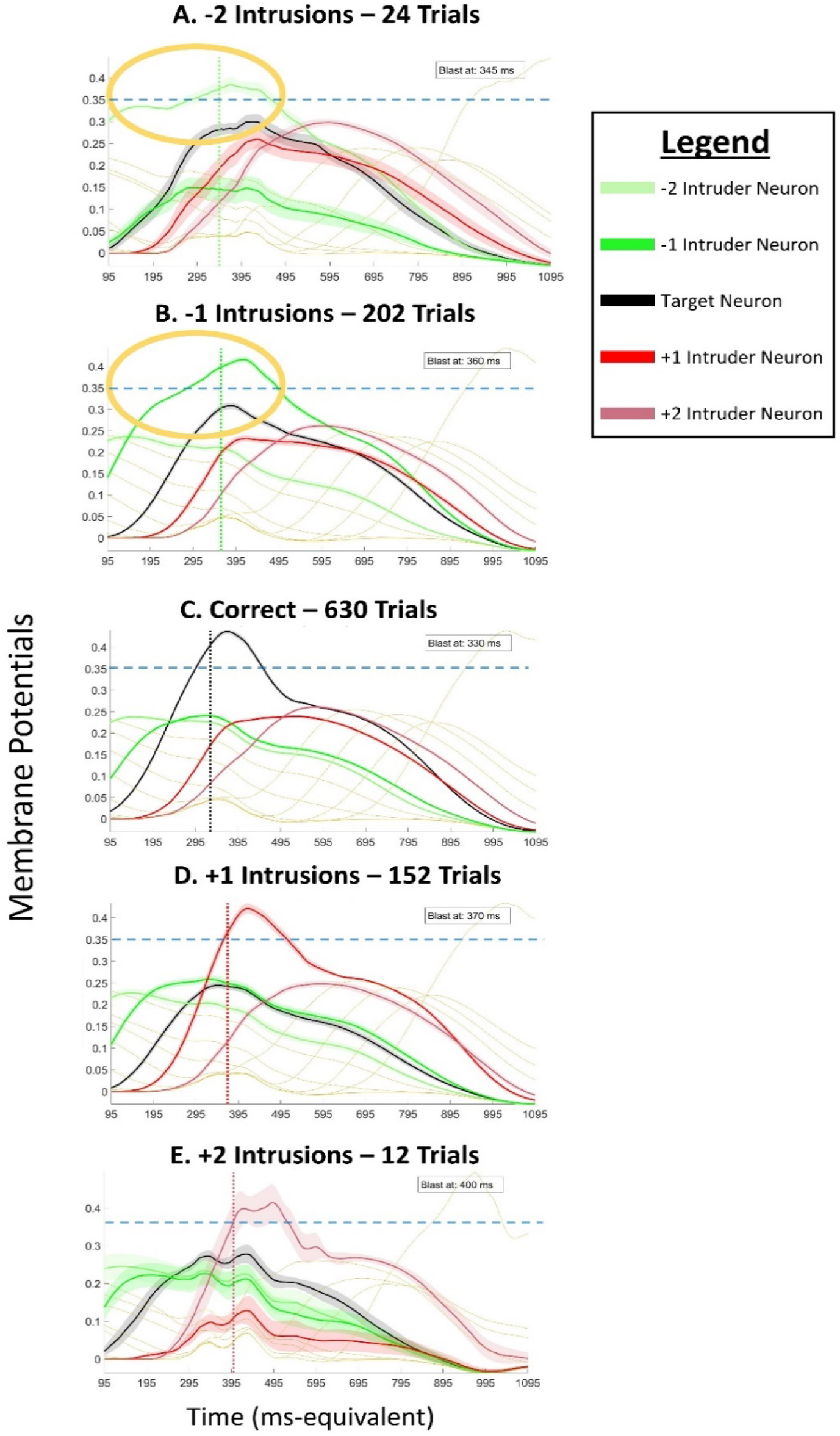
Loss of responsiveness explaining counterintuitive RT patterns. The membrane potentials of the response TFL with the response filter turned off and without any delays to either pathway are plotted. Light green, green, black, red, and light red lines correspond to the VMs of −2, −1, target, +1, and +2 neurons, respectively. The number of trials of each condition is provided, vertical lines indicate average blaster-firing latencies (respective numerical values being provided in plots’ top-right textboxes) and horizontal lines indicate the threshold to the binding pool at 0.35. Note, since these plots are averages of many trials and average blaster firings, even though it looks in some cases as if the blaster fires after the encoding threshold has been crossed, this may not be the case at the single trial level. Indeed, in some trials where an encoded trace rises early, that rise could still follow blaster firing.

**Figure 25.**
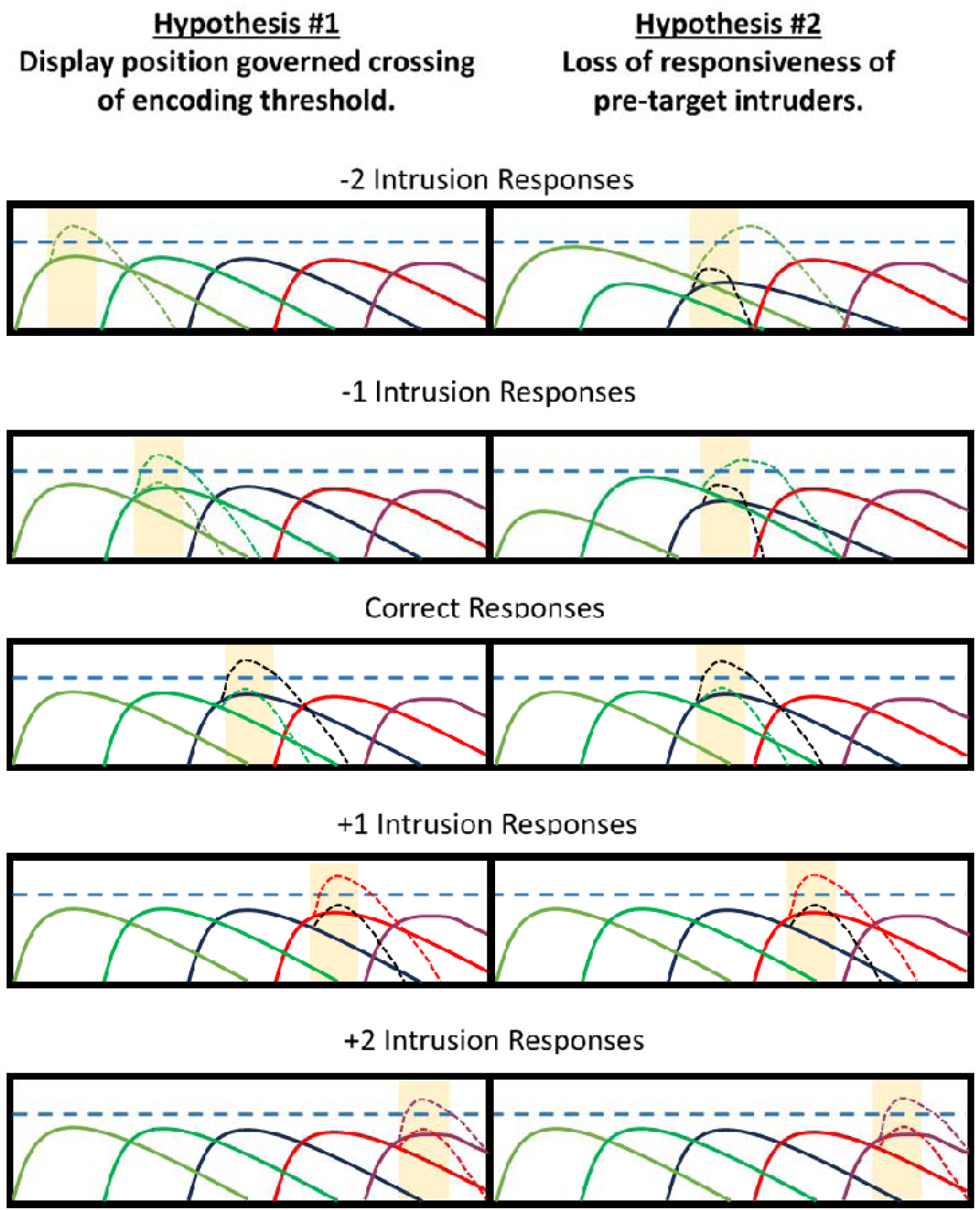
Schematic outline of two possible scenarios (hypotheses) of intrusion errors occurring in human cognition. Light green, green, black, red, and light red curves indicate item-representations of −2, - 1, 0 (target), +1, and +2 items, respectively. The blue dashed lines indicate the encoding threshold, which item-representations must cross to be encoded into WM and subsequently provided as the response. Orange rectangles indicate transient attentional enhancement (TAE). Dashed curves indicate the trajectory an item response would take as a result of blaster firing. For simplicity of presentation, we only show these for the two responses that are most impacted by the blaster. The trajectory for losing items (e.g. black dashed in second row) is due to initially being boosted by blaster firing and then suppressed by competition with winning item. Item representations specifically refer to items’ response dimension and time-intervals of TAE are the only key-feature related element in these plots. Rows illustrate how −2 to +2 intrusions as well as correct responses are generated. Hypothesis 1 (left column) states that item-representations (coloured curves) rise and fall sequentially and that the time-interval of encoding is governed by items’ temporal display positions in the RSVP stream. The second hypothesis (right column) states that the TAE’s timing is more centred around the target-item’s representation. According to Hypothesis 2, item-representations of pre-target intruders in pre-target intrusion trials (top two rows) cross the encoding threshold later than target items in correct trials (third row), which in turn leads to the counterintuitive RT patterns shown by Botella (1992). Time intervals of TAE deployment depicted for the second hypothesis are based on the blaster-firing latencies found with the 2f-ST^2^ model. Due to random variability, there will be amplitude differences in item-representations. As shown, these play an important role in the mechanism we believe leads to pre-target errors.

**Figure 26.**
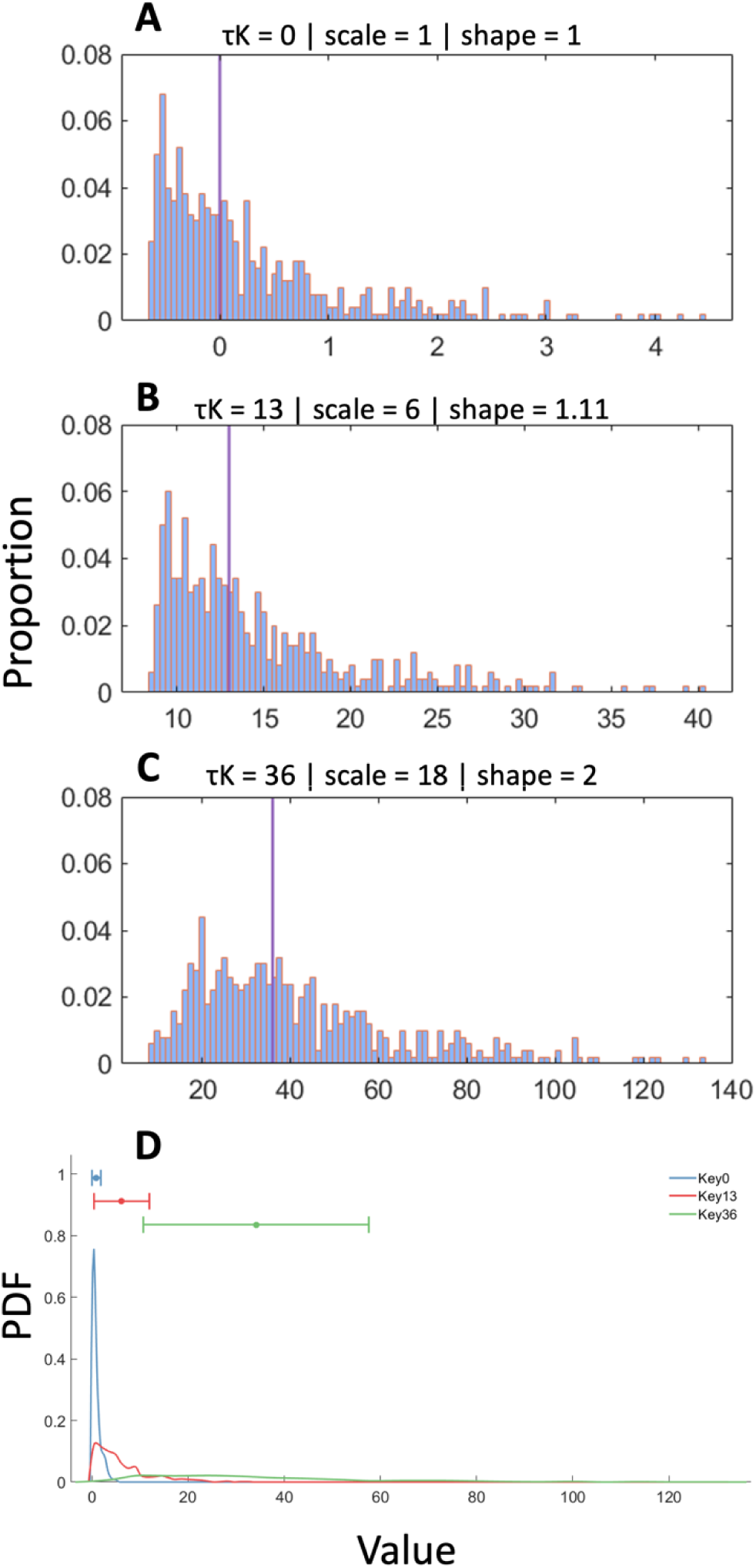
Panels A, B, and C show τK^tr^ distributions for τKs of 0, 13 & 36, respectively. Panel D plots the three corresponding Gamma distributions (denoted G in the equations above) on the same axis to highlight the impact of τK on Gamma distributions’ scale and shape parameters. Blue, red and green lines correspond to Gamma distributions after τKs of 0, 13 and 36, respectively. τK^tr^ distributions are obtained by sampling randomly from the respective Gamma distribution, subtracting the median of G and adding τK.

Indeed, in general, while the global workspace is a seminal contribution to the study of conscious perception, one can debate the neural mechanisms by which the workspace is realised. For example, the competitive interpretation of Dehaene et al. (1998) does not give a mechanism by which multiple feature dimensions can be bound, as required to model distractor intrusions.

The competition in Dehaene et al. (1998)’s workspace would seem to only enable single features to be perceived/encoded at a time. This exclusivity of access to the workspace also makes lag-1 sparing in the attentional blink difficult to model, since it also requires multiple items to pass through the encoding “bottleneck” together; see, for example, Bowman & Wyble (2007), Wyble et al (2009) and Olivers & Meeter (2008).

The ST^2^ family of models propose that the analogue of the workspace is focused on a binding process (c.f. the binding pool). This enables lag-1 sparing and feature conjunctions to be modelled, since temporally proximal items (features of a single or multiple stimuli) can bind together and thereby encode together. Constraints on conscious access then arise through a “latch” on entry into the binding process, implemented as the blaster in the ST^2^ family, which itself is inspired by the TAE idea. Blaster firing opens the gate to encoding, while suppression of the blaster closes it. However, commencement of this suppression can be delayed (see in particular Wyble et al (2009)), potentially allowing multiple items to enter the binding process together. Accordingly, blaster dynamics are central to determining what is bound via the binding pool and thereby encoded, a central theme explored in this paper. We believe this offers a more complete interpretation of encoding and access awareness arising from fast stimulus presentations than that provided by the global workspace model of Dehaene et al. (1998).

### Assessing item representations at the single trial level

A key remaining question concerns item representations at the single trial level. A critical difficulty in behavioural studies is that it is hard to observe the system in its “native state” at the single-trial level, e.g. to know the nature of the representation across response items in a single-trial. Interrogating this representation by looking at the frequency of responding (as the seminal work of Vul et al., 2009), can only be done across many single-trial repetitions (i.e. one can only calculate a proportion/probability across many repetitions, which would correspond to many single-trials). Our modelling in the section focusing on Vul et al. (2009) suggests that the single trial representation in working memory could be uniform across encoded items. This provides an alternative explanation to Vul et al.’s interpretation of their results, but it does not definitively answer the question. We propose that this uncertainty might be resolved in a decoding study that assesses the strength of representation of different items at the single-trial level.

### Predictions for Subjective Visibility

A central objective for this paper has been to provide a more parsimonious alternative to the focused attention-sophisticated guessing model (Botella et al., 2001). This has led to the 2f-ST^2^ model, which suggests a continuum of responding across corrects and intrusion errors. The choice between these two models could be informed by empirical work on the nature of experience during RSVP intrusion error experiments. In particular, the focused attention-sophisticated guessing model (Botella et al., 2001) suggests a qualitative difference between the experience of corrects and errors at the Binding stage and after it. This could be assessed by adding subjective visibility ratings (Bowman et al., 2022; Pincham et al., 2016) to the responding in intrusion error experiments. Botella et al.’s (2001) model would predict a bimodality in subjective visibility responses, with responses from the focussed-attention route generating a (possibly more concentrated) higher mode. The 2f-ST^2^ model predicts that subjective visibility decreases with increased ambiguity with which the competition of response types for access to the Binding Pool was won. This should further hold irrespective of whether the response was correct or not, since there is only one binding mechanism in 2f-ST^2^.

Percepts should be less visible, whenever types are encoded for cortical responses (early Stage 1 layers in the 2f-ST^2^ model) that are already decaying. This is related to the loss of responsiveness phenomenon (Appendix 4), whereby certain types in certain model configurations (and, thus, empirical settings) typically win the competition at the response TFL/ type layer by only a small margin. Appendix 4 provides an in-depth explanation of why −1 intrusion errors in the context of Botella’s RT paradigm (1992) are an example of such types, which, as a result, should be accompanied with low subjective visibility.

While Zivony and Eimer (2020)’s Experiment 2 asked subjects to report confidence in an intrusion error experiment, all other related findings have been obtained in somewhat different paradigms to the classic two feature experiment considered here. Simione et al. (2017) found evidence that errors in integration in RSVP experiments were experienced subjectively in a similar way to correct percepts. Furthermore, in Bowman et al. (2022), Jones et al. (2020) and Pincham et al. (2016), a reduction in subjective visibility and meta-experience was observed when two targets were presented close in time. However, in the experiment considered in Bowman et al. (2022), Jones et al. (2020) and Pincham et al. (2016), an integrated percept was not possible, i.e., the two targets were reported as separate items. Thus, it seems that when errors of integration do generate a coherent object, as they do in Simione et al. (2017) and in intrusion error experiments, they are vividly experienced. This does not fit well with the focused attention-sophisticated guessing model (Botella et al., 2001) idea of a qualitative difference between correct and incorrect responses. In contrast, these findings suggest that correct and incorrect responses are part of the same continuum, with corrects as merely fortunate conjunctions, while still being in line with the notion that the subjective visibility of intrusion errors depends on how clearly intruder items won the competition to enter WM against the target.

### Predictions for Human Electrophysiology

With respect to the 2f-ST^2^ model with no task-filtering in the response TFL/ type layer (replicating the experimental paradigm of Botella and colleagues (e.g., Botella et al. (2001)), a straightforward direction for future research would be to measure the N2pc as well as P3 ERP components. The potential discovery of pre-target intrusion trials being accompanied with *later* ERP components would provide strong evidence in favour of our model architecture and particularly the loss of responsiveness phenomenon generating the RT pattern shown by Botella (1992). See Figure 11, Figure 12, as well as Appendix 4 for more detail. The only relevant ERP study in this context was Botella et al. (2008) and their results were broadly in line with the 2f-ST^2^ model. The authors recorded the P3 component at Pz and showed increased amplitude after correct trials compared to intrusion trials and later P3s for post-target than pre-target intrusions.

Even though these findings clearly fit the 2f-ST^2^ model, a study specifically comparing N2pc or P3 latency of pre- and post-target intrusion *as well as* correct trials is still lacking, as Botella et al. (2008) combined the two types of intrusions into a single condition, when comparing their P3s to those of correct trials.

### Limitations of the 2f-ST^2^ model

We highlight two issues limiting the 2f-ST^2^ model. First, the human response distributions that were replicated by the 2f-ST^2^ model (Figure 9 & Figure 14) reflected only trials where a target or intruder was reported (i.e., non-missed trials). Since the field has yet to explicitly investigate cases where either no response was made or where the responded item was not presented in the trial, which together comprise misses, in intrusion error experiments, our present computational efforts did not account for misses either, and they are not fitted to human data. Future research should consider such responses. Second, in the experiments of Zivony and Eimer (2021) simulated in this paper, grey digit stimuli which lacked the key feature, were presented in pre-target frames to ensure that attentional resources were deployed due to the key feature and not the categorical shift from letters to digits. In the 2f-ST^2^ model, this would mean that in each trial, some types in the response TFL/ type layer meet response pathway task demands prior to the key-target-type. Types corresponding to these non-target digit stimuli would affect other response TFL/ type neurons more strongly because of the lateral inhibition implemented in this layer. These are not currently modelled in the 2f-ST^2^ model.

### Conclusion

We presented the 2f-ST^2^ model, the first neural model accounting for numerous findings reported in the distractor intrusion error literature. The 2f-ST^2^ model explains a wide range of behavioural as well as electrophysiological findings obtained in experiments across three different paradigms (with all distractors or only the +1-distractor sharing the response feature dimension of the target and with subjects being required to provide multiple guesses after each trial). This was achieved via the same underlying model architecture, stressing the significance of the temporal dynamics with which TAE is deployed in the human brain after the task-relevant key feature was detected. In general, different empirical paradigms yield different distributions of correct and error responses due to a shift in the timing of key or response feature processing according to our model. We finally placed our findings into the context of broader cognition and provided predictions made by our computational model to be tested by future empirical studies. These would require meta-cognitive/ meta-experiential judgements to be added to RSVP intrusion error experiments.

Notably, our model proposes that temporal conjunction errors can be accurately modelled within the Simultaneous Type/ Serial Token framework, a broad-scope theory of perception, temporal attention and working memory, without the need for separate pathways for correct performance and errors. Accordingly, correct responses can simply be viewed as fortunate conjunctions and errors as unfortunate ones, arising from the same (single) binding process and the same sources of temporal variability.

## Appendix 1: Stage 1 and 2 Parameter Modifications

### Stage 1

ST^2^’s Stage 1 was duplicated to generate 2f-ST^2^’s key and response pathways. The following changes were made to 2f-ST^2^’s key pathway, in comparison to ST^2^’s Stage 1. Firstly, to decrease TFL/ type layer activation strengths, the item to TFL/ type layer weights were reduced from 0.015 to 0.013. In addition, the TFL/ type layer leak current was increased from 0.07 to 0.0715. Secondly, to ensure that TFL/ type layer ON neurons were suppressed at the correct times, weights from TFL/ type layer OFF neurons to ON neurons were decreased from - 0.12 to −0.15. Finally, to activate the blaster more strongly from the target type node, the TFL/ type layer to blaster Input weight was increased from 0.02003 to 0.02803. Leak current and connection weight settings for the layers in Stage 1 of the response pathway are presented in Tables 2 and 3 of Chennu (2009). Weak lateral inhibition was added between TFL/ type layer on neurons (connection weight −0.05) in the response pathway. Two slight changes were made to Stage 1, since the initial 2f-ST^2^ model (Chennu et al., 2011). The item layer’s bias was set to zero, as this distorted neurons’ membrane potentials, especially when introducing a substantial time-delay to key pathway processing, which was required in novel delay-configurations of the model. Response pathway item layer neurons were furthermore not inhibited by their corresponding off-neurons. We implemented this change to simplify the inhibitory dynamics of this pathway, which due to the lateral inhibition present in its TFL/ type were already rather complex.

### Stage 2

The Stage 2 binding pool was expanded to represent associations between response pathway types and tokens (one per combination of 4 tokens and 25 response types). To compensate for the altered activation dynamics at the token level, Stage 2 weights were adjusted as in Table 3 of Chennu (2009).

## Appendix 2: Smoothing the 2f-ST2 model’s vERPs

The 2f-ST2 model’s smoothed vERPs were computed by first calculating the approximate derivative of original vERPs (i.e. generating a time-series where each point is the difference of adjacent time-points in the original vERP), since (modulo a multiplicative constant, the capacitance) current is the time derivative of the voltage, i.e., how it changes through time.

We further 1-D smoothed the resulting time-series with a gaussian kernel (using MATLAB’s ‘gausswin’ and ‘filter’ functions). We decided against zero-phase filtering (e.g., using MATLAB’s filtfilt function), as it shifted vERP’s peak-latencies more substantially. This property of using zero-phase filtering resulted from filtering derivatives, as the peaks of the original vERPs meant the derivative being close to zero. Having no phase shifts filtering the derivatives therefore meant that time-windows of maximum amplitudes of the original vERPs were transformed to time-windows of minimal (i.e., close to zero) amplitudes. Using the ‘filter’ function solved this issue and led to smoothed vERP time series that approximated time-intervals of maximal amplitudes of original vERPs. The utilized gaussian kernels were standardized to have an area-under-curve (AUC) equal to one prior to any smoothing. The denominator coefficient *a* of the filter-function was furthermore chosen in a way that led to amplitudes similar to those exhibited in our original vERPs (*a* being .05 and .025 for the vN2pc and vP3, respectively). Importantly, this coefficient was constant across different analyses of the same vERP component and *only differed between components*.

## Appendix 3: 2f-ST2 Model Glossary

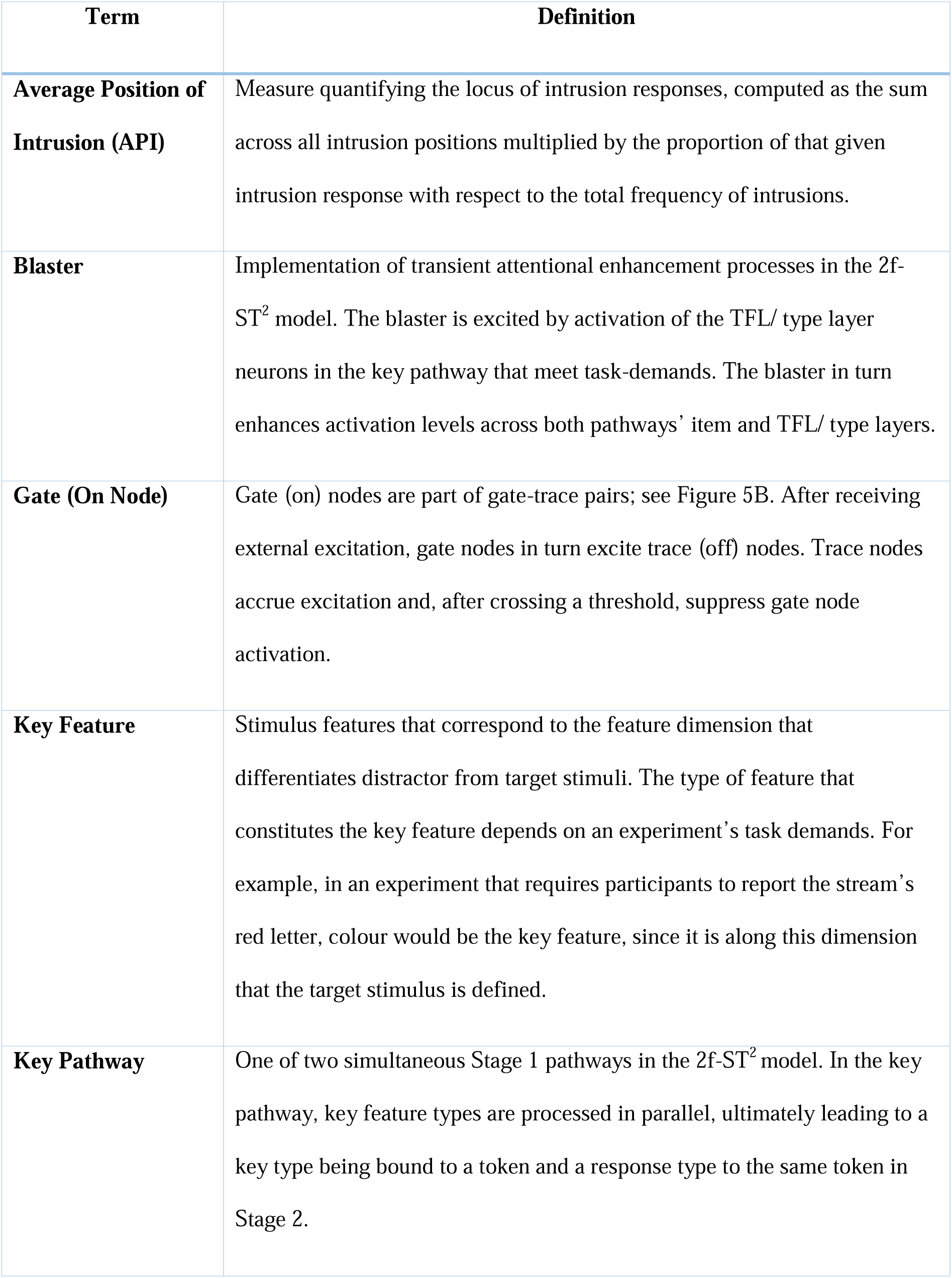
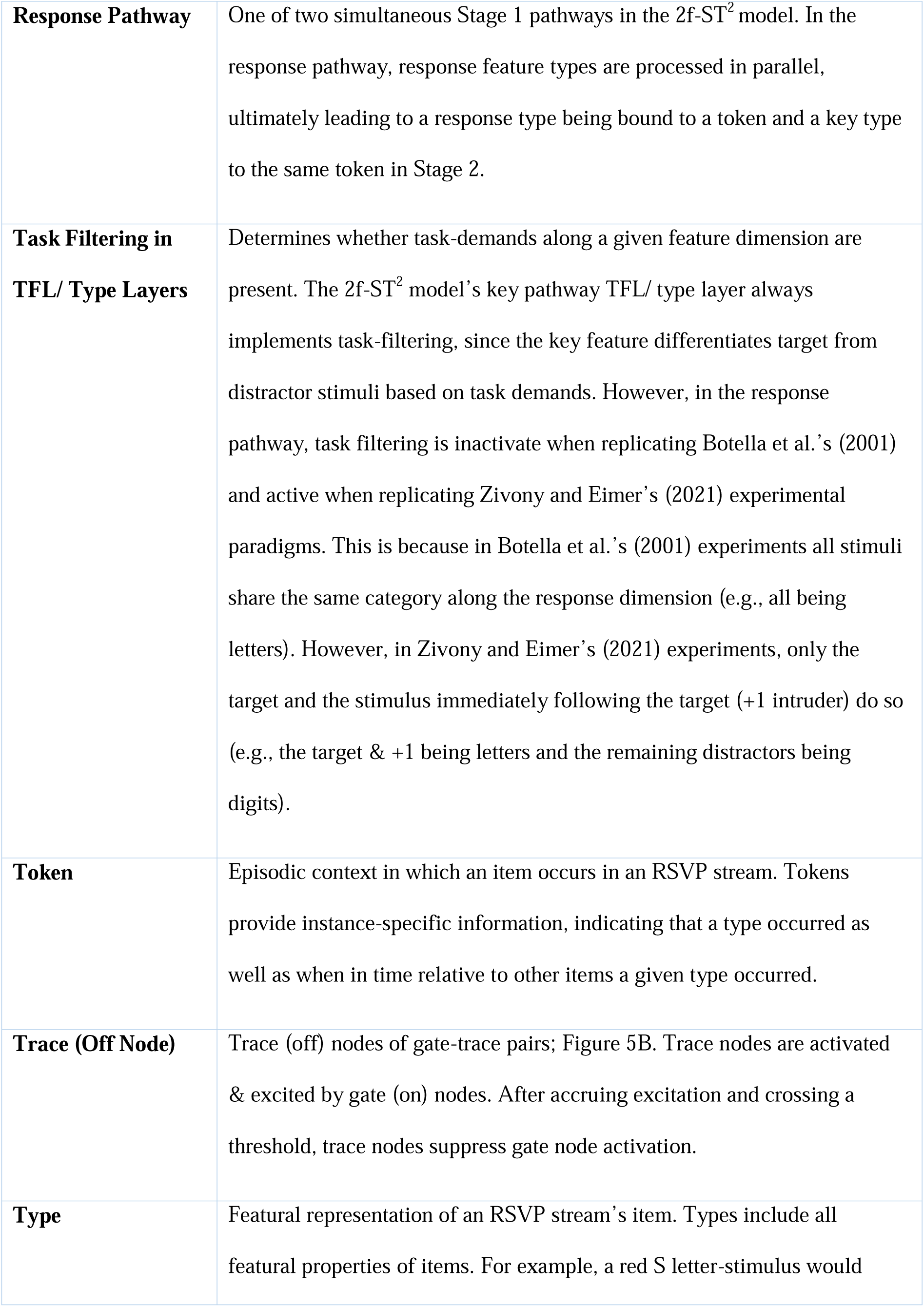
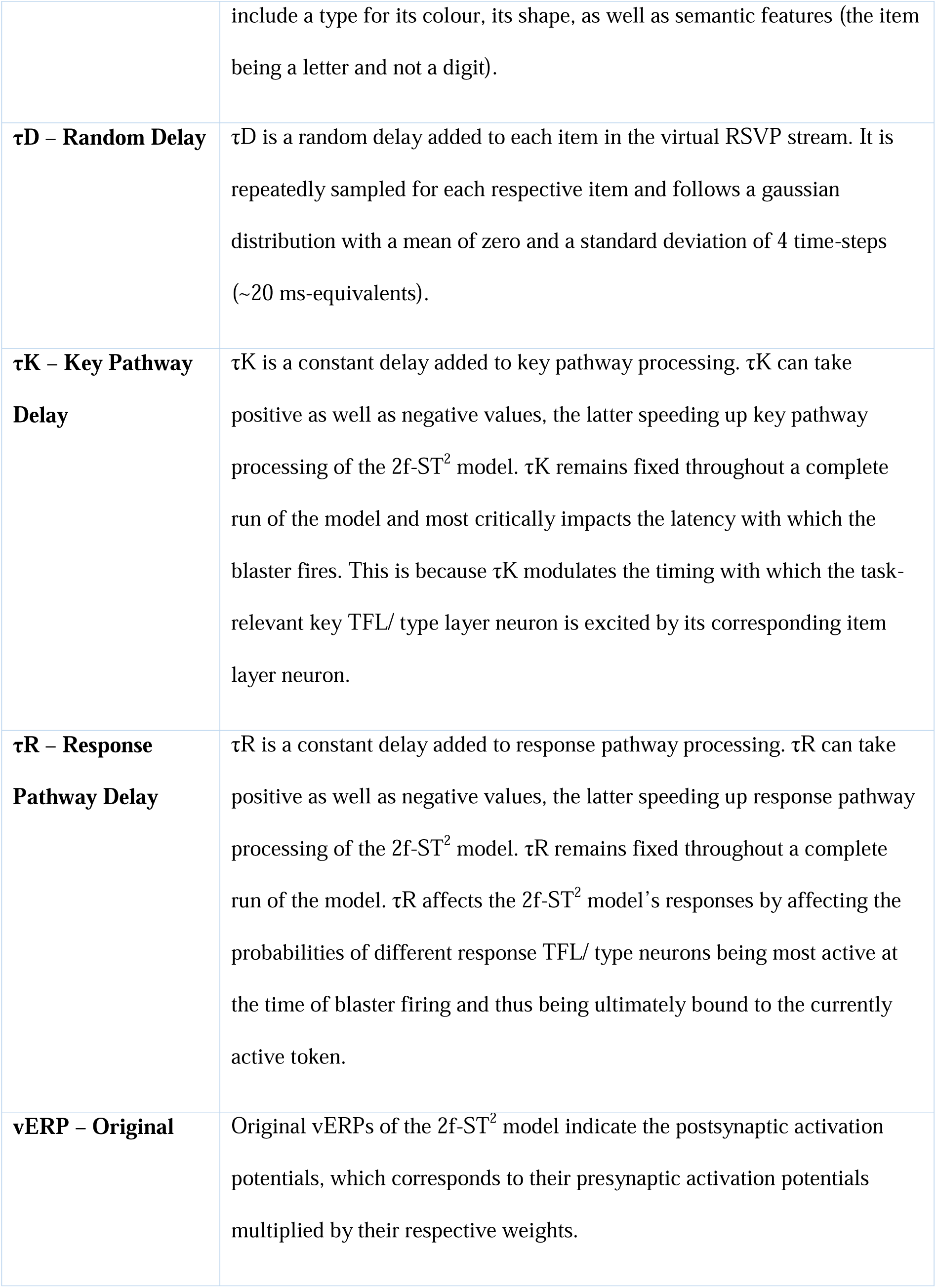
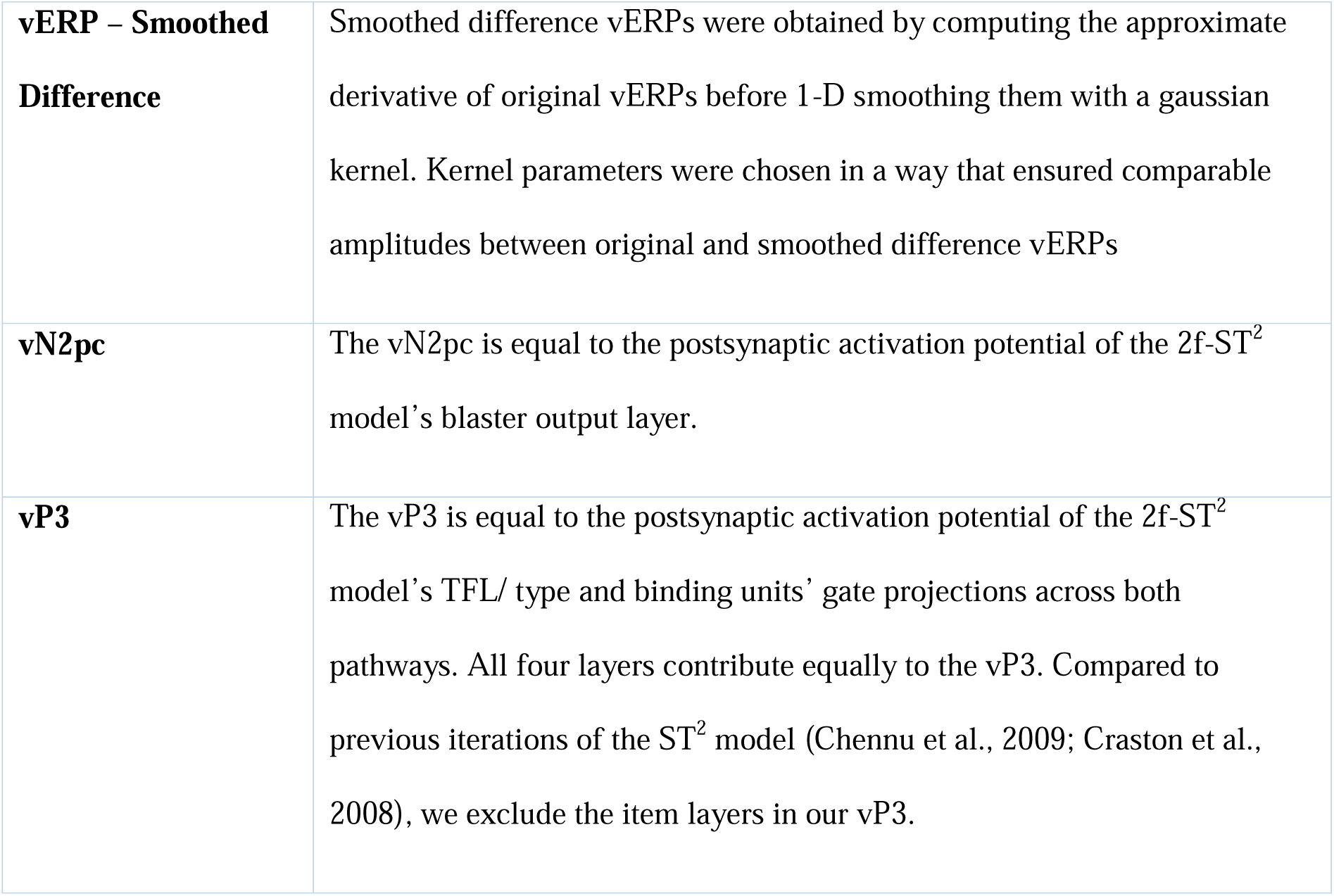

## Appendix 4: The 2f-ST2 Model’s Loss of Responsiveness

This appendix will introduce the 2f-ST^2^ model’s loss of responsiveness issue and explain how it limited capacity to account for Zivony and Eimer’s (2021) electrophysiological results before explaining why it simultaneously allowed us to replicate the counterintuitive reaction times of Botella (1992). *Everything* presented in this appendix was generated by the 2f-ST^2^ model *without Gamma Noise*. As a result, the value of τK used in this appendix to replicate Zivony and Eimer’s (2021) Experiment 1 was smaller than that mentioned in the text body.

The initial 2f-ST^2^ model (Chennu, 2009; Chennu et al., 2011) was developed to replicate the experimental paradigm employed by Botella et al. (2001), in which all stimuli could have been the target stimulus as they all shared the same response dimension (e.g., all being digits). We extended this initial 2f-ST^2^ model to the experiments of Zivony and Eimer (2021), in which *only one* distractor stimulus shared the response dimension of the target, representing the only possibility of committing an intrusion error. We call the 2f-ST^2^ model that lacks Gamma Noise but that can, via a flexible response-pathway filter, replicate both experimental paradigms the *intermediate 2f-ST^2^ model*.

### Loss of responsiveness leading to minimal vP3 differences replicating the paradigm of Zivony and Eimer (2021)

**Error! Reference source not found.** presents the results of running the intermediate 2f-ST^2^ model configured to simulate Zivony and Eimer’s (2021) Experiment 1, via task-filtering in the response TFL/ type layer and a τK of 24. **Error! Reference source not found.**A shows the response distributions of the model, which were in line with those found empirically. However, the between conditions amplitude and latency-differences in the vP3 (**Error! Reference source not found.**B) were small and constituted a limitation of the intermediate 2f-ST^2^ model.

The main factor leading to these small between-condition vP3 differences is that neurons in the response pathways’ TFL/ type layer require the blaster’s enhancement for their membrane potentials (VM) to cross the effective threshold of that layer and in turn excite their respective binding pool units. **Error! Reference source not found.** demonstrates this, plotting the response TFL/ type layer’s VMs of the target- and +1-neuron in correct and intrusion trials. The blaster’s enhancement is uniquely important in correct trials to enable the target neuron to cross the threshold to the response binding pool (henceforth called T-C trials and plotted as a black solid line in **Error! Reference source not found.**). This is because the target neuron in correct trials has a VM close to, but just below, the threshold (at .35, indicated by the green horizontal line) for a rather long time-period prior to the blaster firing.

**Error! Reference source not found.** shows that the target neuron for T-C outcomes (black solid line) starts to ascend *earlier* than the +1-neuron in intrusion trials (called +1-I, red dashed line). For example, at around 150 milliseconds, the black solid line is considerably above the red dashed line. If the latency difference of these two neurons’ VM-onsets would translate to their postsynaptic potentials, it would lead to vP3s that more clearly resemble the latency-differences between correct and intrusion trials shown in Zivony and Eimer’s (2021) human ERPs (Figure 15). However, because the target neuron for T-C outcomes struggles to cross the threshold (at a VM of 0.35) without help from the blaster, it crosses the threshold at about the same time as the +1 neuron in +1-I (the +1-intruder being bound in intrusion trials) outcomes (see solid black and dashed red lines crossing green line at around 395 milliseconds), leading to the vP3s of **Error! Reference source not found.**. Note that blaster-firing latencies in vERP or neurons’ postsynaptic activation plots, such as those of **Error! Reference source not found.** & **Error! Reference source not found.**, depict *averages* across all trials of a given response condition, and for some trials, the firing will be a good deal earlier or indeed later.

In Hosseini (2022) we provide a more thorough analysis of these dynamics, demonstrating that 1) decreasing excitation from the item layer is the main reason for the flattening of the T-C’s VM-trace at around 300 ms, which is simply due to the sequential presentation of items to the model, and 2) τK is the parameter that modulates the extent of responsiveness being lost. To summarise the loss of responsiveness of the response TFL/ type target-& +1-neurons: these neurons competed very little with each other without a key-delay and it was usually unambiguous which response feature will be encoded to a given token. However, once τK is large (24 time-steps), an important competition (due to lateral inhibition) emerges between them, and the winning neuron usually beats its competitor by just a small margin.

Hence, the range of trials that are now categorized as correct or intrusion changes and become more similar to one another. This is why their vP3s are so similar: because the underlying trials’ characteristics (in terms of blaster-firing latencies and input stimulus strengths) are so similar.

### Loss of responsiveness explaining the counterintuitive reaction times of Botella (1992)

To reiterate, in Figure 11 we demonstrated that the 2f-ST^2^ model is able to replicate counterintuitive reaction time (RT) patterns provided by Botella (1992). These RT patterns are counterintuitive because correct responses were given faster than pre-target intrusion errors. To understand why the 2f-ST^2^ model generates this pattern too, we show the membrane potentials of the response TFL/ type layer for the intermediate 2f-ST^2^ model that replicated Botella (1992) (i.e., with response pathway task-filtering being inactive and no delays to either pathway) in **Error! Reference source not found.**. Light green, green, black, red, and light red lines in **Error! Reference source not found.** correspond to the VMs of −2, −1, target, +1, and +2 neurons, respectively. Panels A, B, D and E show −2-, −1-, +1- and +2-intrusions, respectively.

Panel C shows the correct condition. **Error! Reference source not found.** can therefore be viewed as the analogue of **Error! Reference source not found.**. **Error! Reference source not found.**B shows the membrane potential of the −1-neuron in −1-trials, which is strikingly similar to that of the target-neuron in correct trials (previously called T-C), shown in **Error! Reference source not found.**. As described with respect to the T-C neuron previously, the −1-neuron in −1-trials demonstrates a similar decrease in its membrane potential’s gradient just before the blaster enhances neurons’ activation-levels (indicated with a yellow circle in **Error! Reference source not found.**B). This membrane potential gradient-decrease is even more visible looking at the −2-neuron in −2-trials (**Error! Reference source not found.**A). Thus, this also seems to be a case of pre-target neurons losing their responsiveness: pre-target intrusions occur after the blaster enhances the respective pre-target neuron when its driving item-layer excitation has already started to decline. As a result, the pre-target neuron requires comparably more time to receive enough enhancement from the blaster to excite its corresponding binding pool unit sufficiently for its type to be bound to the currently active token. This additional time ultimately results in the 2f-ST^2^ model replicating the counterintuitive RT patterns found by Botella (1992) in humans (Figure 11).

**Two Hypotheses of Intrusion Errors:** We provide a schematic illustration of how the loss of responsiveness of pre-target intruder-neurons is not only helpful in this configuration of the 2f-ST^2^ model, but presumably necessary for our model to replicate the counterintuitive RT patterns. **Error! Reference source not found.** displays two hypotheses of how intrusion errors might occur in each of its columns. Each row of **Error! Reference source not found.** depicts how different responses are provided according to the two hypotheses. In these plots, curved and coloured lines represent item-representations along the response dimension, where in this material, item is used in its general sense not with regard to the item-layer in the model. Light green, green, black, red, and light red colours correspond to −2 intruder-, −1 intruder-, target-, +1 intruder, and +2 intruder-items, respectively. The encoding threshold, as proposed by Zivony and Eimer (2020) and as modelled in the 2f-ST^2^ model as the connection between response TFL/ type layer and the binding pool, is represented as a blue dashed line. Time-intervals of transient attentional enhancement (TAE, the blaster in 2f-ST^2^ model) are depicted as orange areas. Item-representations specifically refer to response features and the time-interval of TAE (or the blaster-firing) is the only aspect of the key feature (or key pathway) present in these plots. Importantly, in this illustration, items need to cross the encoding threshold to be encoded into WM.

Hypothesis 1 (left column of **Error! Reference source not found.**) displays probably the simplest explanation of how intrusion errors occur. According to Hypothesis 1, intruders’ (response) item-representations cross the encoding threshold completely governed by their display position (i.e., temporal position in the RSVP stream). Therefore, which item is encoded into WM is only a matter of *when* the encoding process itself takes place. Importantly, this presumably still involves TAE-mechanisms. This concept has similarities to the sophisticated guessing route of Botella et al.’s (2001) model, with the critical time, t_c_, determining which response feature would be bound. Indeed, Botella et al.’s (2001) Figure 6A looks strikingly similar to **Error! Reference source not found.**’s left plots. However, such dynamics would not lead to the RT patterns observed by Botella (1992). Instead, RT patterns reflecting the order of item-representations would be the result: RT (pre-target intrusions) < RT (correct) < RT (post-target intrusions). The counter-intuitive RT patterns observed in humans (Botella, 1992) in addition to the response TFL/ type layer’s VMs shown in **Error! Reference source not found.**, however, strongly support the second hypothesis illustrated in **Error! Reference source not found.** (right column).

According to Hypothesis 2, the range of TAE-timings is more centred around the correct item. We based the TAE-intervals shown in **Error! Reference source not found.**’s right column on the average blaster-firing latencies presented in **Error! Reference source not found.**. Importantly, for **Error! Reference source not found.**’s blaster-firing latencies correct trials on average occur earlier than those of pre-target intrusions. This is because in correct trials, the target key type is generally stronger and thus activates the blaster circuit earlier. Hypothesis 2 proposes that pre-target intrusions are associated with later TAE deployment, when pre-target intruder item’s representations have already started to decrease in magnitude. This can be seen in the top two rows of **Error! Reference source not found.**’s right column, in which both green curves are declining before the time-interval of TAE (yellow box)^7^. This is in contrast to how correct reports are provided according to Hypothesis 2 (third row of **Error! Reference source not found.**’s right column). TAE impacts the target item (black curve) in correct trials at a point in time at which its item representation is increasing in magnitude. Therefore, less assistance is required from TAE to cross the encoding threshold in this case (see rapid increase of dashed black curve), which leads to the target item in correct trials being encoded faster than pre-target intruder items in pre-target intrusion trials. This is the case despite the target item itself being presented after pre-target intruder items in the RSVP stream. This latency-difference in encoding is illustrated by the time-interval at which the black and green curves cross the encoding threshold in the first three rows of **Error! Reference source not found.**’s right column. The last two rows of **Error! Reference source not found.**’s right column illustrate how post-target intrusions are made according to Hypothesis 2. +1-intrusions are made because TAE impacts processes at a time interval at which the target-item’s representation has been decreasing, but the +1-intruder’s representation has been increasing, leading to the +1-intruder winning the competition and being encoded into WM. +2-intrusions are made similarly.

## Appendix 5: Gamma Noise Methods

The Gamma Noise distribution was created prior to a run of the model. For each trial in a given run, one value of Gamma Noise was sampled from this distribution. We present example distributions in **Error! Reference source not found.** (panels A, B and C), for which a shape and a scale parameter had to be set. We also present equations below formalising the procedure.

The shape parameter determines how right-skewed (low shape value) or symmetric (high shape value) the distribution is. For our purposes, a low shape value was desired because selective attention has an absolute bound on *how quickly* it can be deployed, but not on *how slowly*. For example, the TAE will never be deployed *before previous cognitive processes have completed* (e.g. the N2pc typically carries an onset latency of 150-200 ms (Eimer, 1996; Zivony & Eimer, 2021)). However, it does not depend on subsequent cognitive processes in the same manner. Our variation of the shape parameter was motivated by speed-accuracy RT trade-offs (Draheim et al., 2019). RTs are distributed more symmetrically as one instructs participants to respond more accurately. RT distributions become more right skewed (i.e., more probability mass for fast RTs) if participants are instructed to respond as fast as possible. We therefore let Gamma Noise’s shape parameters depend on τK, as low τKs lead to fast RTs (i.e. a low shape value) and large τKs lead to slow RTs (i.e. higher shape values).

The scale parameter enabled us to control the extent of variability added to τK and, thereby, the temporal jitter added to blaster firing latencies. Thus, larger τK values (simulating less salient key-features in experiments) should imply more variability with which the blaster fires. We therefore set Gamma Noise’s scale to 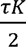, rounding down to the nearest natural number.

*τK^tr^* denotes the key delay imposed on a particular trial, *tr*. We sample a value for *τK^tr^* anew on each trial. This is done according to the equations below, which ensured that τK was the median of the distribution and thus, that it was equally likely to sample a τK^tr^ that was smaller or larger than the run’s fixed τK. For example, if τK was 36, the value of τK^tr^ adopted for an individual trial would be sampled from the distribution shown in **Error! Reference source not found.**C. Critically, even though values smaller or larger than τK both made up 50% of each distribution, the difference of values > τK to τK was generally larger than that of values < τK, simulating the characteristic of the hard limit to fast TAE-deployment. **Error! Reference source not found.**D plots the three distributions’ probability density functions on the same axis to illustrate how the values added to τK were modulated by its fixed value, being smaller (lower scale) and more right-skewed (lower shape) for low values of τK (simulating very salient key features).

More formally, on each trial, we sample a new key feature time constant, denoted *τK^tr^*, where indicates a number of time units, as follows:

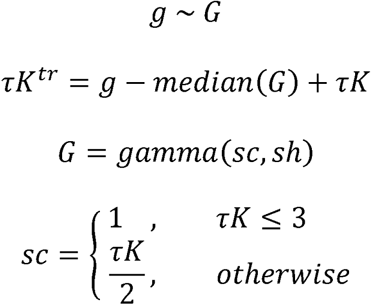

and

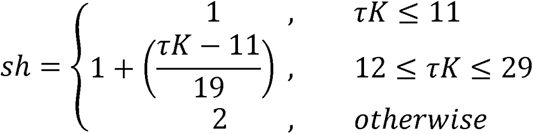

## Acknowledgements

This project has received funding from the European Union’s Horizon 2020 research and innovation programme under (Grant Agreement No. 896192 to A.Z.) and from the Economic and Social Research Council (Grant No. ES/V002708/1 to M.E.).

## Footnotes

1 Of course, the selection is from an activation pattern across items in short-term memory, which might be being treated as if it is a probability distribution, but on a single-trial it cannot be one, since each item in the stream has only been presented once.

2 We will also consider situations in which more than one unit in each pathway can encode, but we postpone consideration of that possibility for now.

3 Although, unlike for the key pathway, the influence of response task-demand is not illustrated in Figure 5A.

4 As an illustration of the issue, imagine there were 100 trials with this distribution: ***-2*** *= 10, **-1** = 20, **0** = 40, **+1** = 20, **+2** = 10*. If you take out the *40* at ***0*** and normalise the other entries (with some rounding adjustments to give whole numbers) to a probability distribution, you obtain, *17%, 33%, gap, 33%, 17%*. However, if you take out the *10* at ***+2*** and normalise you obtain *11%, 22%, 45%, 22%, gap*. That is, the *10* trials (really *10%*) for ***-2*** become *17%* in one case and *11%* in the other? This is the problem. If we estimate the gap with interpolation, we obtain *∼10%* for the ***-2*** in both cases.

5 Chennu et al. (2009) contains a minor reporting error, since τD was sampled from a gaussian distribution with mean of 0 ms and standard deviation of 20 ms as opposed to the 15 ms reported.

6 Although, Jones, Pincham, Gootjes-Dreesbach & Bowman (2020); Bowman, Jones, Pincham, Fleming, Cleeremans & Smith (2022); and Pincham, Bowman & Szucs (2016) that strictly there is a dissociation between conscious perception and working memory encoding in this binding process.

7 When comparing **Error! Reference source not found.** and **Error! Reference source not found.**, do note that the former is an average across many individual trials, while the latter is imagining an individual trial. Thus, the **Error! Reference source not found.** pattern will not show exactly the same pattern as **Error! Reference source not found.** because it averages across many different individual trial patterns with different blaster firing latencies and loss of responsiveness dynamics.

## References

Akyürek, E. G., Leszczyński, M., & Schubö, A. (2010). The temporal locus of the interaction between working memory consolidation and the attentional blink. Psychophysiology. 10.1111/j.1469-8986.2010.01033.x

Alsufyani, A., Hajilou, O., Zoumpoulaki, A., Filetti, M., Alsufyani, H., Solomon, C. J.,…& Bowman, H. (2019). Breakthrough percepts of famous faces. Psychophysiology, 56(1), e13279.

Alsufyani, A., Harris, K., Zoumpoulaki, A., Filetti, M., & Bowman, H. (2021). Breakthrough percepts of famous names. Cortex, 139, 267–281.

Avilés, A., Bowman, H., & Wyble, B. (2020). On the limits of evidence accumulation of the preconscious percept. Cognition, 195, 104080. 10.1016/j.cognition.2019.104080

Bae, J., Jung, K., James, O., Suzuki, S., & Kim, Y. J. (2025). Frontal engagement in perceptual integration under low subjective visibility. NeuroImage, 305, 120984.

Balsdon, T., Wyart, V., & Mamassian, P. (2024). Metacognitive evaluation of postdecisional perceptual representations. Journal of vision, 24(4), 2. 10.1167/jov.24.4.2

Battelli, L., Pascual-Leone, A., & Cavanagh, P. (2007). The ‘when’ pathway of the right parietal lobe. Trends in Cognitive Sciences, 11(5), 204–210. 10.1016/j.tics.2007.03.001

Binder, M., Gociewicz, K., Windey, B., Koculak, M., Finc, K., Nikadon, J., Derda, M., & Cleeremans, A. (2017). The levels of perceptual processing and the neural correlates of increasing subjective visibility. Consciousness and cognition, 55, 106–125. 10.1016/j.concog.2017.07.010

Botella, J. (1992). Target-specified and target-categorized conditions in RSVP tasks as reflected by detection time. Bulletin of the Psychonomic Society, 30(3), 197–200. 10.3758/BF03330440

Botella, J., Barriopedro, M. I., & Suero, M. (2001). A model of the formation of illusory conjunctions in the time domain. Journal of Experimental Psychology: Human Perception and Performance, 27(6), 1452–1467. 10.1037/0096-1523.27.6.1452

Botella, J., & Eriksen, C. W. (1992). Filtering versus parallel processing in RSVP tasks. Perception & Psychophysics. 10.3758/BF03211627

Botella, J., Privado, J., de Liaño, B. G. G., & Suero, M. (2011). Illusory conjunctions reflect the time course of the attentional blink. Attention, Perception, and Psychophysics, 73(5), 1361–1373. 10.3758/s13414-011-0112-z

Bowman, H., & Avilés, A. (2021). Fragile Memories for Fleeting Percepts. 10.31234/osf.io/fr3bg

Bowman, H., & Avilés, A. (2022). No Subliminal Memory for Spaced Repeated Images in Rapid-Serial-Visual-Presentation Streams. Psychological Science, 33(11), 1959–1965. 10.1177/09567976221100229

Bowman, H., Filetti, M., Alsufyani, A., Janssen, D., & Su, L. (2014). Countering countermeasures: Detecting identity lies by detecting conscious breakthrough. PloS one, 9(3), e90595.

Bowman, H., Jones, W., Pincham, H., Fleming, S., Cleeremans, A., & Smith, M. (2022). Modelling the simultaneous encoding/serial experience theory of the perceptual moment: A blink of meta-experience. Neuroscience of Consciousness, 2022(1), niac003. 10.1093/nc/niac003

Bowman, H., & Wyble, B. (2007). The simultaneous type, serial token model of temporal attention and working memory. Psychological Review, 114(1), 38–70. 10.1037/0033-295X.114.1.38

Bowman, H., Wyble, B., Chennu, S., & Craston, P. (2008). A reciprocal relationship between bottom-up trace strength and the attentional blink bottleneck: Relating the LC-NE and ST2 models. Brain Research, 1202, 25–42. 10.1016/j.brainres.2007.06.035

Chennu, S. (2009). The temporal spotlight of attention: Computational and electrophysiological explorations. [University of Kent]. http://www.cs.kent.ac.uk/pubs/2009/3054

Chennu, S., Bowman, H., & Wyble, B. (2011). Fortunate Conjunctions Revived: Feature Binding with the 2f-ST2 Model. Proceedings of the 33rd Annual Conference of the Cognitive Science Society, 1992, 182–196.

Chennu, S., Craston, P., Wyble, B., & Bowman, H. (2009). Attention Increases the Temporal Precision of Conscious Perception: Verifying the Neural-ST2 Model. PLOS Computational Biology, 5(11), e1000576.

Chun, M. M. (1997). Temporal binding errors are redistributed by the attentional blink. Perception & Psychophysics, 59(8), 1191–1199. 10.3758/BF03214207

Chun, M. M., & Potter, M. C. (1995). A two-stage model for multiple target detection in rapid serial visual presentation. Journal of Experimental Psychology: Human Perception and Performance, 21(1), 109–127. 10.1037/0096-1523.21.1.109

Craston, P., Wyble, B., Chennu, S., & Bowman, H. (2008). The Attentional Blink Reveals Serial Working Memory Encoding: Evidence from Virtual and Human Event-related Potentials. Journal of Cognitive Neuroscience, 21(3), 550–566. 10.1162/jocn.2009.21036

Dehaene, S., Kerszberg, M., & Changeux, J. P. (1998). A neuronal model of a global workspace in effortful cognitive tasks. Proceedings of the National Academy of Sciences, 95(24), 14529–14534.

Dehaene, S., Sergent, C., & Changeux, J. P. (2003). A neuronal network model linking subjective reports and objective physiological data during conscious perception. Proceedings of the National Academy of Sciences, 100(14), 8520–8525.

Draheim, C., Mashburn, C. A., Martin, J. D., & Engle, R. W. (2019). Reaction time in differential and developmental research: A review and commentary on the problems and alternatives. Psychological Bulletin, 145(5), 508.

Eimer, M. (1996). The N2pc component as an indicator of attentional selectivity. Electroencephalography and Clinical Neurophysiology. 10.1016/0013-4694(96)95711-9

Eimer, M. (2014). The neural basis of attentional control in visual search. Trends in Cognitive Sciences, 18(10), 526–535. 10.1016/j.tics.2014.05.005

Feldman, J. (2013). The neural binding problem(s). Cognitive Neurodynamics, 7(1), 1–11. 10.1007/s11571-012-9219-8

Henderson, C. M., & McClelland, J. L. (2020). Intrusions into the shadow of attention: A new take on illusory conjunctions. Attention, Perception, & Psychophysics, 82(2), 564–584. 10.3758/s13414-019-01893-3

Hosseini, M. (2022). My mind’s playing tricks on me: Understanding event integration in rapid stimulus streams. [University of Kent]. https://kar.kent.ac.uk/id/eprint/97369

Hosseini, M., Zivony, A., Eimer, M., Wyble, B., & Bowman, H. (2024). Transient Attention Gates Access Consciousness: Coupling N2pc and P3 Latencies Using Dynamic Time Warping. The Journal of Neuroscience, 44(26), e1798232024. 10.1523/JNEUROSCI.1798-23.2024

Jaeger, C., Glim, S., Dimulescu, C., Ries, A., Sorg, C., & Wohlschläger, A. (2020). Segregated Co-activation Patterns in the Emergence of Decision Confidence During Visual Perception. Frontiers in systems neuroscience, 14, 557693. 10.3389/fnsys.2020.557693

Jimenez, M., Villalba-García, C., Luna, D., Hinojosa, J. A., & Montoro, P. R. (2019). The nature of visual awareness at stimulus energy and feature levels: A backward masking study. Attention, perception & psychophysics, 81(6), 1926–1943. 10.3758/s13414-019-01732-5

Jones, W., Pincham, H., Gootjes-Dreesbach, E. L., & Bowman, H. (2020). Fleeting Perceptual Experience and the Possibility of Recalling Without Seeing. Scientific Reports, 10(1), 1–19. 10.1038/s41598-020-64843-2

Kanwisher, N. G. (1987). Repetition blindness: Type recognition without token individuation. Cognition, 27(2), 117–143. 10.1016/0010-0277(87)90016-3

Kiss, M., Van Velzen, J., & Eimer, M. (2008). The N2pc component and its links to attention shifts and spatially selective visual processing. Psychophysiology. 10.1111/j.1469-8986.2007.00611.x

Klein, R. M., Ishigami, Y., & Murray, N. E. (2023). Slippage of the attentional beam when searching in space and in time. Cognition, 241, 105610. 10.1016/j.cognition.2023.105610

Kristjánsson, Á., & Egeth, H. (2020). How feature integration theory integrated cognitive psychology, neurophysiology, and psychophysics. Attention, Perception, & Psychophysics, 82(1), 7–23.

Lau, H. C., & Passingham, R. E. (2006). Relative blindsight in normal observers and the neural correlate of visual consciousness. Proceedings of the National Academy of Sciences, 103(49), 18763–18768.

Lawrence, D. H. (1971). Two studies of visual search for word targets with controlled rates of presentation*. Perception & Psychophysics, 10(2), 85–89. 10.3758/BF03214320

Levi D. M. (2008). Crowding--an essential bottleneck for object recognition: a mini-review. Vision research, 48(5), 635–654. 10.1016/j.visres.2007.12.009

Luck, S. J., & Hillyard, S. (1994). Spatial filtering during visual search: Evidence from human electrophysiology. Journal of Experimental Psychology: Human Perception and Performance, 20(5), 1000–1014. 10.1037/0096-1523.20.5.1000

Nieuwenhuis, S., Holmes, B. D., Gilzenrat, M. S., & Cohen, J. D. (2005). The role of the locus coeruleus in mediating the attentional blink: A neurocomputational theory. Journal of Experimental Psychology: General, 134(3), 291–307. 10.1037/0096-3445.134.3.291

Nunez, P. L., & Srinivasan, R. (2007). Electroencephalogram. Scholarpedia, 2, 1348.

Olivers, C. N. L., & Meeter, M. (2008). A Boost and Bounce Theory of Temporal Attention. Psychological Review, 115(4), 836–863. 10.1037/a0013395

Op De Beeck, H. P., Haushofer, J., & Kanwisher, N. G. (2008). Interpreting fMRI data: Maps, modules and dimensions. Nature Reviews Neuroscience, 9(2), 123–135. 10.1038/nrn2314

Pelli, D. G., Palomares, M., & Majaj, N. J. (2004). Crowding is unlike ordinary masking: Distinguishing feature integration from detection. Journal of Vision, 4(12), 12–12. 10.1167/4.12.12

Pelli, D. G., & Tillman, K. A. (2008). The uncrowded window of object recognition. Nature neuroscience, 11(10), 1129–1135. 10.1038/nn.2187

Pincham, H. L., Bowman, H., & Szucs, D. (2016). The experiential blink: Mapping the cost of working memory encoding onto conscious perception in the attentional blink. Cortex, 81, 35–49. 10.1016/j.cortex.2016.04.007

Polich, J. (2007). Updating P300: An integrative theory of P3a and P3b. Clinical Neurophysiology, 118(10), 2128–2148. 10.1016/j.clinph.2007.04.019

Potter, M. C. (1976). Short-term conceptual memory for pictures. Journal of Experimental Psychology. Human Learning and Memory, 2(5), 509–522.

Rausch, M., Hellmann, S., & Zehetleitner, M. (2018). Confidence in masked orientation judgments is informed by both evidence and visibility. Attention, perception & psychophysics, 80(1), 134–154.

Reeves, A., & Sperling, G. (1986). Attention gating in short-term visual memory. Psychological Review, 93(2), 180–206. 10.1037/0033-295X.93.2.180

Roberts, S., & Pashler, H. (2000). How persuasive is a good fit? A comment on theory testing. Psychological Review, 107(2), 358–367. 10.1037/0033-295X.107.2.358

Scholte, H. S., & De Haan, E. H. F. (2025). Beyond binding: From modular to natural vision. Trends in Cognitive Sciences, 29(6), 505–515. 10.1016/j.tics.2025.03.002

Seibold, V. C., Balke, J., & Rolke, B. (2023). Temporal attention. Frontiers in Cognition, 2. https://www.frontiersin.org/journals/cognition/articles/10.3389/fcogn.2023.1168320

Simione, L., Akyürek, E. G., Vastola, V., Raffone, A., & Bowman, H. (2017). Illusions of integration are subjectively impenetrable: Phenomenological experience of Lag 1 percepts during dual-target RSVP. Consciousness and Cognition, 51, 181–192. 10.1016/j.concog.2017.03.004

Shih, S. I., & Sperling, G. (2002). Measuring and modeling the trajectory of visual spatial attention. Psychological review, 109(2), 260.

Su, L., Wyble, B., Zhou, L.-Q., Wang, K., Wang, Y.-N., Cheung, E. F. C., Bowman, H., & Chan, R. C. K. (2015). Temporal perception deficits in schizophrenia: Integration is the problem, not deployment of attentions. Scientific Reports, 5, 9745.

Tallon-Baudry, C. (2003). Oscillatory synchrony and human visual cognition. Journal of Physiology-Paris, 97(2-3), 355–363.

Treisman, A., & Schmidt, H. (1982). Illusory conjunctions in the perception of objects. Cognitive Psychology, 14(1), 107–141. 10.1016/0010-0285(82)90006-8

Treisman, A. (1998). Feature binding, attention and object perception. Philosophical Transactions of the Royal Society of London. Series B: Biological Sciences, 353(1373), 1295–1306. 10.1098/rstb.1998.0284

Treisman, A. M., & Gelade, G. (1980). A feature-integration theory of attention. Cognitive Psychology, 12(1), 97–136. 10.1016/0010-0285(80)90005-5

Vul, E., Hanus, D., & Kanwisher, N. (2009). Attention as Inference: Selection Is Probabilistic; Responses Are All-or-None Samples. Journal of Experimental Psychology: General. 10.1037/a0017352

Woodman, G. F., & Luck, S. J. (1999). Electrophysiological measurement of rapid shifts of attention during visual search. Nature. 10.1038/23698

Wyble, B., Bowman, H., & Nieuwenstein, M. (2009). The Attentional Blink Provides Episodic Distinctiveness: Sparing at a Cost. Journal of Experimental Psychology: Human Perception and Performance, 35(3), 787–807. 10.1037/a0013902

Wyble, B., Potter, M. C., Bowman, H., & Nieuwenstein, M. (2011). Attentional episodes in visual perception. Journal of Experimental Psychology: General, 140(3), 488–505. 10.1037/a0023612

Zeki, S. M. (1978). Functional specialisation in the visual cortex of the rhesus monkey. Nature, 274(5670), 423–428. 10.1038/274423a0

Zivony, A., & Eimer, M. (2022). The diachronic account of attentional selectivity. Psychonomic Bulletin & Review, 29(4), 1118–1142. 10.3758/s13423-021-02023-7

Zivony, A., & Eimer, M. (2021). Distractor intrusions are the result of delayed attentional engagement: A new temporal variability account of attentional selectivity in dynamic visual tasks. Journal of Experimental Psychology. General, 150(1), 23–41. 10.1037/xge0000789

Zivony, A., & Eimer, M. (2020). Perceptual competition between targets and distractors determines working memory access and produces intrusion errors in rapid serial visual presentation (RSVP) tasks. Journal of Experimental Psychology: Human Perception and Performance, 46(12), 1490–1510. 10.1037/xhp0000871

Zivony, A., & Eimer, M. (2024). Attention and feature binding in the temporal domain. Psychonomic Bulletin & Review, 31(6), 2599–2610. 10.3758/s13423-024-02493-5

